# Antihypertensive effect of brain-targeted mechanical intervention with passive head motion

**DOI:** 10.1101/2020.09.21.305706

**Authors:** Shuhei Murase, Naoyoshi Sakitani, Takahiro Maekawa, Daisuke Yoshino, Kouji Takano, Ayumu Konno, Hirokazu Hirai, Taku Saito, Sakae Tanaka, Keisuke Shinohara, Takuya Kishi, Yuki Yoshikawa, Takamasa Sakai, Makoto Ayaori, Hirohiko Inanami, Koji Tomiyasu, Atsushi Takashima, Toru Ogata, Hirotsugu Tsuchimochi, Shinya Sato, Shigeyoshi Saito, Kohzoh Yoshino, Yuiko Matsuura, Kenichi Funamoto, Hiroki Ochi, Masahiro Shinohara, Motoshi Nagao, Yasuhiro Sawada

## Abstract

Physical exercise is known to be beneficial for various brain functions. However, the mechanisms behind the positive effects of exercise on the brain remain to be elucidated. Here we show that passive head motion in hypertensive rats, which reproduces the mechanical accelerations generated in their heads during moderate-velocity treadmill running, decreases the expression of angiotensin II type 1 receptor (AT1R) in astrocytes in the rostral ventrolateral medulla (RVLM), thereby lowering blood pressure. Passive head motion generates interstitial fluid movement that is estimated to exert shear stress with an average magnitude of <1 Pa on the cells in the rat medulla. Fluid shear stress of a sub-Pa magnitude decreases AT1R expression in cultured astrocytes. In hypertensive rats, inhibition of interstitial fluid movement following hydrogel introduction to the RVLM eliminates the antihypertensive effects of passive head motion and treadmill running. Furthermore, vertically oscillating chair riding by hypertensive adult humans, which reproduces the mechanical accelerations generated in their heads during light jogging or fast walking, lowers their blood pressure. Our findings indicate that moderate mechanical intervention can have antihypertensive effects by modulating the function of RVLM astrocytes through interstitial fluid shear stress. We anticipate that mechanical regulation is responsible for a variety of the positive effects of physical exercise on human health, particularly those related to brain functions.

## Introduction

Exercise is known to be effective as a therapeutic/preventative measure for numerous physical disorders and diseases including hypertension^1, 2^, which is a major cause of stroke and cardiovascular diseases making the biggest risk factor for death worldwide^3^. However, the mechanisms underlying the antihypertensive effect of exercise remain to be elucidated.

Whereas the majority (>90%) of human hypertension comprises essential hypertension, the cause of which is unidentifiable^4^, long-term regulation of blood pressure (BP) has been recognized to be largely dependent on sodium excretion adjusting systems mainly involving kidney functions^5^. Elevated activity of the sympathetic nervous system also importantly contributes to the development of hypertension^6–8^. Rostral ventrolateral medulla (RVLM), which is located in the brainstem, plays a critical role in determining the basal activity of the sympathetic nervous system, and its functional integrity is essential for the maintenance of basal vasomotor tone and regulation of BP^6, 9^. Angiotensin II (Ang II) is the major bioactive peptide of the renin-angiotensin system (RAS), and is known to regulate BP as well as other biological processes such as cell growth/apoptosis/migration, inflammation, and fibrosis^10^. The biological effects of Ang II are mediated by its interaction with two distinct high-affinity G protein-coupled receptors, Ang II type 1 receptor (AT1R) and type 2 receptor. Of these receptors, AT1R is responsible for most of the known physiological and pathophysiological processes related to Ang II. Whereas RAS is involved in the functional regulation of various “peripheral” organs and tissues such as kidney and vessels, it also regulates brain functions within the blood-brain barrier, including the control and maintenance of sympathetic nerve activity and cognitive ability^11^. In particular, the role of AT1R signaling in the RVLM in cardiovascular regulation has been extensively studied and demonstrated. For example, the pressor and depressor responses to Ang II and Ang II antagonists, respectively, injected into the RVLM have been reported to be enhanced in spontaneously hypertensive rats (SHRs)^12, 13^. We have previously demonstrated that treadmill running at moderate velocities alleviates the sympathetic nerve activity, involving attenuation of AT1R signaling in the RVLM of stroke-prone spontaneously hypertensive rats (SHRSPs)^14^, a substrain of SHRs that exhibit severer hypertension as compared with SHRs^15^. However, the details about the changes in AT1R signaling in the RVLM of these hypertensive rats have yet to be elucidated. It remains unclear what type(s) of cells (e.g., neurons or astrocytes) are primarily responsible for the altered AT1R signaling in the RVLM of SHRs or SHRSPs. Furthermore, the causal relationship between the increased AT1R signal activity in the RVLM and high BP in SHRs or SHRSPs in their steady state (i.e., apart from their responses to pharmacological interventions) is left unrevealed.

AT1R has also been shown to play a vital role in regulating a variety of physiological or pathological processes, including cellular responses to mechanical perturbations^16, 17^. For example, mechanical stretching of cardiac myocytes activates AT1R signaling^18^, and fluid shear stress (FSS) of average 1.5 Pa lowers AT1R expression in human vein endothelial cells^19^. Although intervening the Ang II-AT1R system through pharmacological approaches, such as administration of angiotensin-converting enzyme inhibitor or selective AT1R blocker, has been established as an effective therapeutic strategy for hypertension^20^, mechano-responsive attenuation of AT1R signaling has not been clinically utilized as an antihypertensive measure.

Many of physical workouts, particularly aerobic exercises, involve vertical body motions, which generate mechanical accelerations in the head at the time of foot contact with the ground (i.e., landing). The importance of mechanical loads is well established in the physiological regulation of bones, the stiffest organ that only allows tiny deformation^21^. Osteocytes, the mechanosensory cells embedded in bones^22^, are assumed to undergo minimal deformations under physiological conditions. We have reported that FSS on osteocytes derived from interstitial fluid flow induced upon physical activity plays an important role in maintaining bone homeostasis^23^. Given that the brain is not a rigid organ, minimally deforming forces or stress distribution changes in the brain during exercise or even activity of daily living (e.g., walking) mthese results support our hypothesis ay produce beneficial effects. We have previously shown that in the prefrontal cortex (PFC) of rodents, moderate mechanical intervention-induced FSS modulates serotonin signaling in the neurons in situ^24^. Based on these previous findings together with the distribution of interstitial fluid throughout the whole brain, we hypothesized that moderate mechanical intervention might have antihypertensive effects involving FSS-mediated modulation of AT1R signaling in the RVLM.

## Results

### Application of cyclical mechanical intervention to the head by passive motion lowers the BP in SHRSPs

To determine the effects of a mechanical intervention of a moderate intensity on BP, we first sought to develop an experimental system that reproduces the acceleration generated in the head during rats’ treadmill running at a modest velocity (20 m/min), a typical experimental intervention to test the effects of physical exercise on rats^25, 26^. In a recent study, we observed that treadmill running of rats (20 m/min) generated 5-mm vertical oscillation of their heads with ∼1.0 × *g* peak accelerations and 2-Hz frequency; therefore, we developed a “passive head motion” (PHM) system to produce 2-Hz 5-mm vertical oscillation exerting 1.0 × *g* acceleration peaks in the heads of rats^24^. In the current study, we examined the effects of mechanical intervention on BP in SHRSPs, using the PHM system. Similar to the antihypertensive effect of treadmill running on SHRs or SHRSPs that we and others reported previously^15, 25, 27^, application of PHM (30 min/day, 28 consecutive days; see Fig. 1a) significantly lowered their BP as compared to their controls (Fig. 1b,c), whereas heart rate (HR) was not significantly affected by PHM (Fig. 1d). Anesthesia alone (daily 30 min) did not significantly alter the BP in SHRSPs (Extended Data Fig. 1a), indicating that the antihypertensive effect resulted specifically from PHM. The anti-cardiac hypertrophy effect of PHM on SHRSPs (Fig. 1e) as well as the lack of these PHM effects on control normotensive rats (Wistar-Kyoto: WKY) (Fig. 1b,c,e) were also consistent with previous reports describing treadmill running as an antihypertensive intervention for SHRs^27^. As was observed in our treadmill running experiments^14^, PHM decreased 24-h urinary norepinephrine excretion of SHRSPs (Fig. 1f). This suggests that PHM mitigates the sympathetic hyperactivity^28^. Collectively, these results support our hypothesis that cyclical application of moderate mechanical intervention to the head has an antihypertensive effect. Notably, ≥4-week PHM significantly decreased or delayed the stroke incidence in the SHRSPs (Extended Data Fig. 1b−d).

**Fig. 1.**
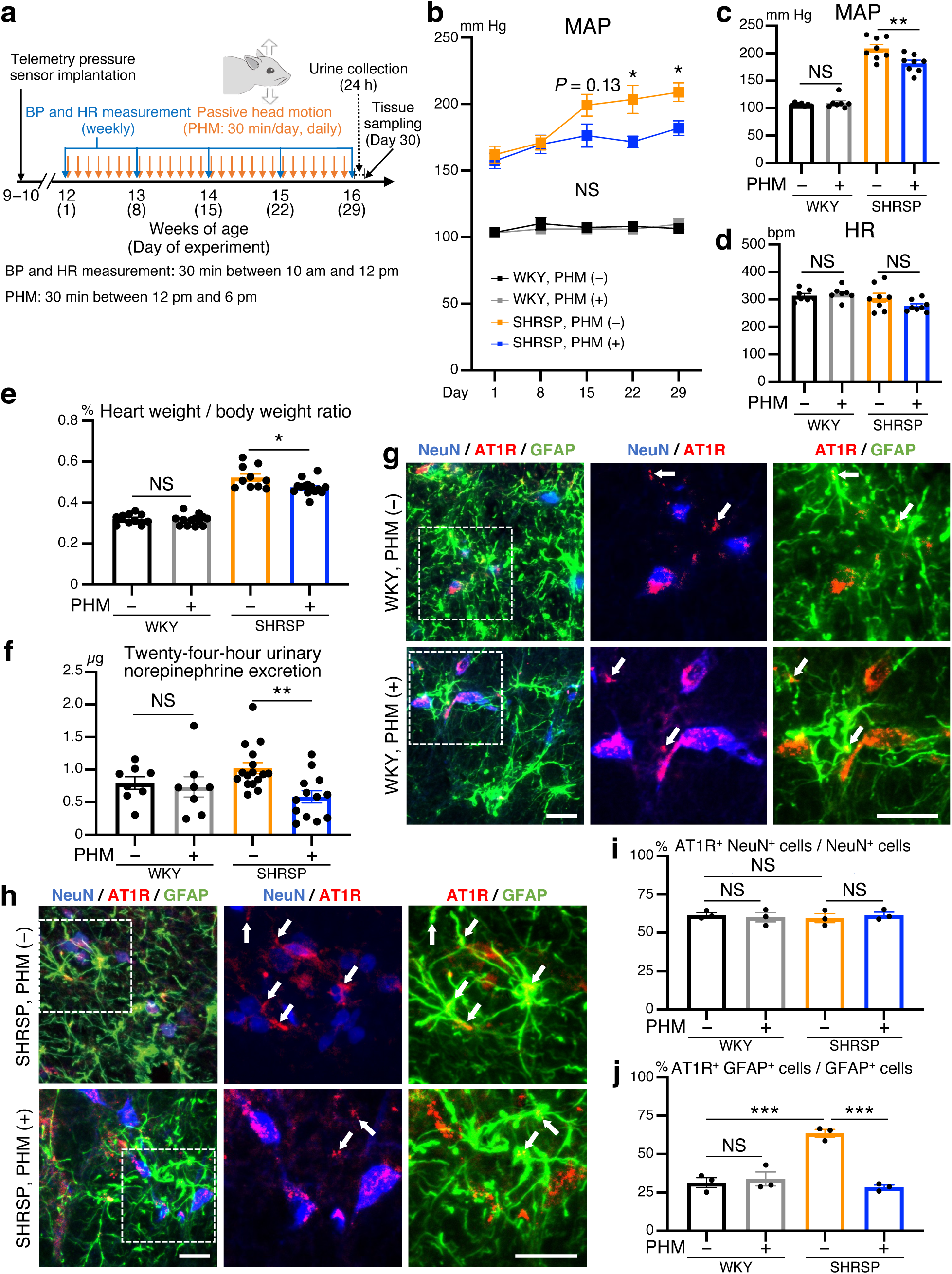
Application of cyclical mechanical intervention to the head by passive motion lowers the BP in SHRSPs, and AT1R expression in their RVLM astrocytes. **a,** Schematic representation of the experimental protocol to analyze the effects of PHM on the BP in rats. b,c, Time courses (b) and values on Day 29 (c) of the MAP in WKY rats and SHRSPs, subjected to either daily PHM or anesthesia only [b, SHRSP PHM (−) vs. (+): *P* = 0.1344 for Day15, *P* = 0.0110 for Day 22, *P* = 0.0463 for Day 29; WKY PHM (−) vs. (+): *P* > 0.9999 for Day 15, Day 22, and Day 29. c, *P* = 0.9739 for column 1 vs. 2, *P* = 0.0046 for column 3 vs. 4. *n* = 7 rats for each group of WKY; *n* = 8 rats for each group of SHRSP]. d, HR values on Day 29 (*P* = 0.9650 for column 1 vs. 2, *P* = 0.2362 for column 3 vs. 4. *n* = 7 rats for each group of WKY; *n* = 8 rats for each group of SHRSP). e, Relative heart weight (heart weight / whole body weight) measured on Day 30 [*P* = 0.9866 for column 1 vs. 2, *P* = 0.0152 for column 3 vs. 4. *n* = 10 rats for WKY, PHM (−); *n* = 13 rats for WKY, PHM (+); *n* = 10 rats for SHRSP, PHM (−); *n* = 14 rats for SHRSP, PHM (+)]. f, Twenty-four-hour (Day 29 to Day 30) urinary norepinephrine excretion [*P* = 0.9854 for column 1 vs. 2, *P* = 0.0085 for column 3 vs. 4. *n* = 8 rats for each group of WKY; *n* = 16 rats for SHRSP, PHM (−); *n* = 13 rats for SHRSP, PHM (+)]. g,h, Micrographic images of anti-NeuN (blue), anti-GFAP (green) and anti-AT1R (red) immunostaining of the RVLM of WKY rats (g) and SHRSPs (h), either left sedentary (top) or subjected to PHM (bottom) under anesthesia (30 min/day, 28 days). Higher magnification images (center and right) refer to the areas indicated by dotted rectangles in low magnification images (left). Arrows point to anti-AT1R immunosignals that overlap with anti-GFAP, but not anti-NeuN, immunosignals in merged images. Scale bars, 50 µm. Images are representative of three rats. i,j, Quantification of AT1R-positive neurons (i) and astrocytes (j) in the RVLM of WKY rats and SHRSPs, either left sedentary or subjected to PHM. Fifty NeuN-positive (NeuN^+^) cells and 100 GFAP-positive (GFAP^+^) cells were analyzed for each rat (i: *P* = 0.9602 for column 1 vs. 2, *P* = 0.9215 for column 1 vs. 3, *P* = 0.9313 for column 3 vs. 4. j: *P* = 0.9455 for column 1 vs. 2, *P* = 0.0004 for column 1 vs. 3, *P* = 0.0002 for column 3 vs. 4. *n* = 3 rats for each group). Data are presented as mean ± s.e.m. \**P* < 0.05; \*\**P* < 0.01; \*\*\**P* < 0.001; NS, not significant; two-way repeated measures ANOVA with Bonferroni’s post hoc multiple comparisons test (b) or one-way ANOVA with Tukey’s post hoc multiple comparisons test (c−f,i,j).

Next, we characterized PHM as an antihypertensive intervention by testing for various directions, frequencies, and amplitudes (peak magnitudes of acceleration). PHM that generated acceleration peaks of 1.0 × *g* in the rostral-caudal direction, but not in the left-right direction, had antihypertensive effects on SHRSPs (Extended Data Fig. 2a–c), suggestive of the directional selectivity. Regarding frequency, vertical PHM (peak magnitude of 1.0 × *g*) of 0.5 Hz, but not of 0.2 Hz, lowered the BP in SHRSPs to the approximately same extent as 2-Hz PHM did (Extended Data Fig. 3a–c). Furthermore, PHM generating a peak magnitude of 0.5 × *g*, but not 0.2 × *g*, was approximately as antihypertensive as 1.0-×-*g* PHM (Extended Data Fig. 3a,d,e). These results suggest the existence of a threshold and a plateau phase of frequency and amplitude (magnitude) of PHM in terms of its antihypertensive effects.

### PHM down-regulates AT1R expression in RVLM astrocytes of SHRSPs

We then looked into the mechanism of how PHM alleviated the development of hypertension in SHRSPs. We previously reported that down-regulation of the AT1R signaling in the RVLM is responsible for the treadmill running-induced sympathoinhibition in SHRSPs^14^. Given the mechanical regulation of AT1R expression in endothelial cells^19^, we examined whether PHM modulated the AT1R expression in RVLM neurons and astrocytes of SHRSPs. In our histochemical analysis, we defined neuronal nuclei (NeuN)-positive cells as neurons^29^ and glial fibrillary acidic protein (GFAP)-positive cells as astrocytes^30^. PHM (30 min/day, 28 days) did not significantly change the relative population of AT1R-expressing neurons and astrocytes in the RVLM of WKY rats (Fig. 1g). In contrast, 4-week PHM significantly decreased the expression of AT1R in the astrocytes, but not in the neurons, of SHRSPs’ RVLM (Fig. 1h). Notably, the AT1R expression in the RVLM neurons was comparable between WKY rats and SHRSPs, either with or without PHM (Fig. 1i). In contrast, the AT1R expression was significantly higher in the RVLM astrocytes of SHRSPs without PHM (Fig. 1j, columns 1 and 3). PHM lowered the AT1R expression in the RVLM astrocytes of SHRSPs to the level equivalent to that of WKY rats (Fig. 1j, columns 1, 2, and 4). Taken together, the AT1R expression in RVLM astrocytes appeared to be correlated with the antihypertensive effect of PHM on SHRSPs. In line with this observation, 4-week treadmill running of SHRSPs also decreased the AT1R expression in their RVLM astrocytes, but not neurons (Extended Data Fig. 4a−c).

### PHM alleviates the sensitivity of the RVLM in SHRSPs to Ang II or Ang II antagonist

We next sought to examine whether the PHM-induced decrease in AT1R expression in the RVLM astrocytes of SHRSPs (Fig. 1j, columns 3 and 4) was functionally relevant to the suppression of AT1R signaling. To this end, we analyzed the pressor responses to Ang II injected into the unilateral RVLM of WKY rats and SHRSPs, either subjected to 4-week PHM or left sedentary under anesthesia (30 min/day, 28 days) (Fig. 2a). As we previously reported^14^, SHRSPs without PHM exhibited significantly greater pressor response to Ang II administered to the RVLM than WKY rats (Fig. 2b, compare between top and bottom of left panels; Fig. 2c, compare columns 1 and 3). Four-week PHM alleviated the pressor response to Ang II injected into the RVLM of SHRSPs, but not of WKY rats (Fig. 2b, compare left and right; Fig. 2c, compare columns 1 vs. 2 and 3 vs. 4). Furthermore, the depressor response to Ang II antagonist injected into the unilateral RVLM^13^ was also mitigated by 4-week PHM in SHRSPs, but not in WKY rats (Fig. 2d,e). These results support the functional relevance of the PHM-induced decrease in the AT1R expression in the RVLM astrocytes of SHRSPs (Fig. 1j).

**Fig. 2.**
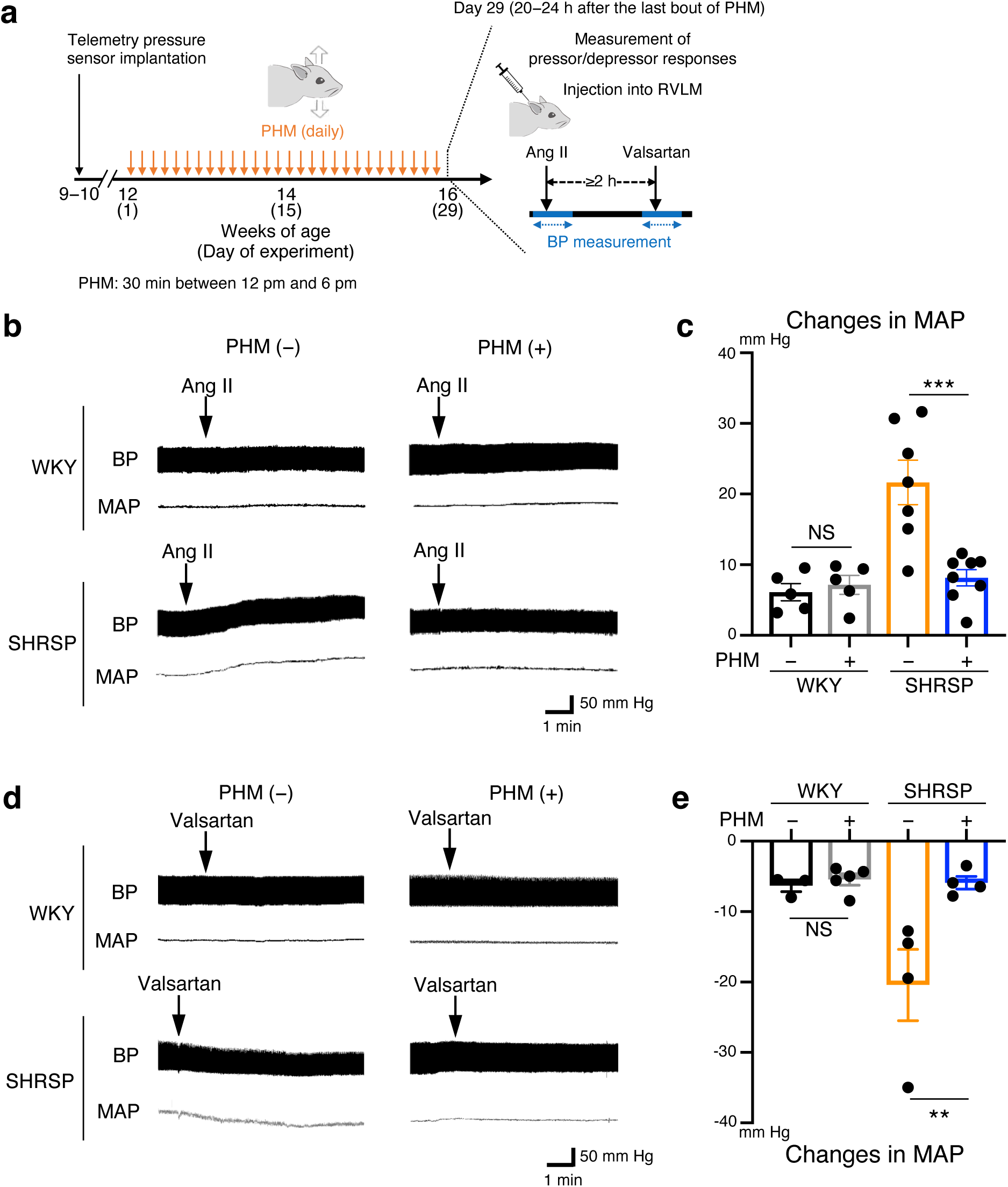
PHM alleviates the sensitivity of RVLM in SHRSPs to Ang II or valsartan. **a,** Schematic representation of the experimental protocol to analyze the effects of PHM on the sensitivity to Ang II or valsartan injected into the unilateral RVLM. Ang II (100 pmol) was injected into the unilateral RVLM of WKY rats and SHRSPs, either left sedentary (daily anesthesia) or subjected to PHM (30 min/day, 28 days), with their BP monitored under urethane anesthesia. Injection of valsartan (100 pmol) into the RVLM was conducted at least 2 h after the injection of Ang II. **b,** Representative trajectories of BP (top in each panel) and MAP (bottom in each panel). Arrows point to the time of the initiation of RVLM injection of Ang II. Right- angled scale bar, 1 min / 50 mm Hg. **c,** Quantification of MAP change caused by Ang II injection [*P* = 0.9876 for column 1 vs. 2, *P* = 0.0003 for column 3 vs. 4. *n* = 5 rats for each group of WKY; *n* = 7 rats for SHRSP, PHM (−); *n* = 8 rats for SHRSP, PHM (+)]. **d,e,** Effects of the RVLM injection of valsartan (100 pmol) examined as in **(b,c)** [**e**: *P* = 0.9953 for column 1 vs. 2, *P* = 0.0099 for column 3 vs. 4. *n* = 3 rats for WKY, PHM (−); *n* = 5 rats for WKY, PHM (+); *n* = 4 rats for each group of SHRSP]. Data are presented as mean ± s.e.m. \*\**P* < 0.01; \*\*\**P* < 0.001; NS, not significant; one-way ANOVA with Tukey’s post hoc multiple comparisons test.

To examine whether the increased AT1R expression in the RVLM astrocytes of SHRSPs was associated with their development of hypertension, we manipulated the AT1R signaling by introducing exogenous expression of AT1R-associated protein (AGTRAP), which interacts with AT1R and tempers the Ang II-mediated signals by promoting AT1R internalization^31^. To this end, we used an adeno-associated virus (AAV)-mediated gene delivery system^32^. AAV serotype 9 (AAV9) vectors were injected locally to transduce the RVLM cells (Fig. 3a and Extended Data Fig. 5a). To achieve astrocyte- and neuron-specific gene expression, we used the AAV9 vectors that harbored mouse GFAP promoter (AAV-GFAP) and rat neuron-specific enolase (NSE) promoter (AAV-NSE), respectively (Fig. 3a). Because these vectors contained a region encoding GFP and 2A sequence of porcine teschovirus-1 (P2A; self-cleaving peptides^33^) (Fig. 3a), observation of the green fluorescence allowed us to identify the cells in which transgene was expressed (Fig. 3b,c and Extended Data Fig. 5a−e). The AAV-mediated expression of AGTRAP in astrocytes (Fig. 3b) but not in neurons (Fig. 3c) of the bilateral RVLMs in SHRSPs significantly lowered the BP as compared with their control SHRSPs in which only GFP was virally expressed in the RVLM astrocytes or neurons (Fig. 3d,e).

**Fig. 3.**
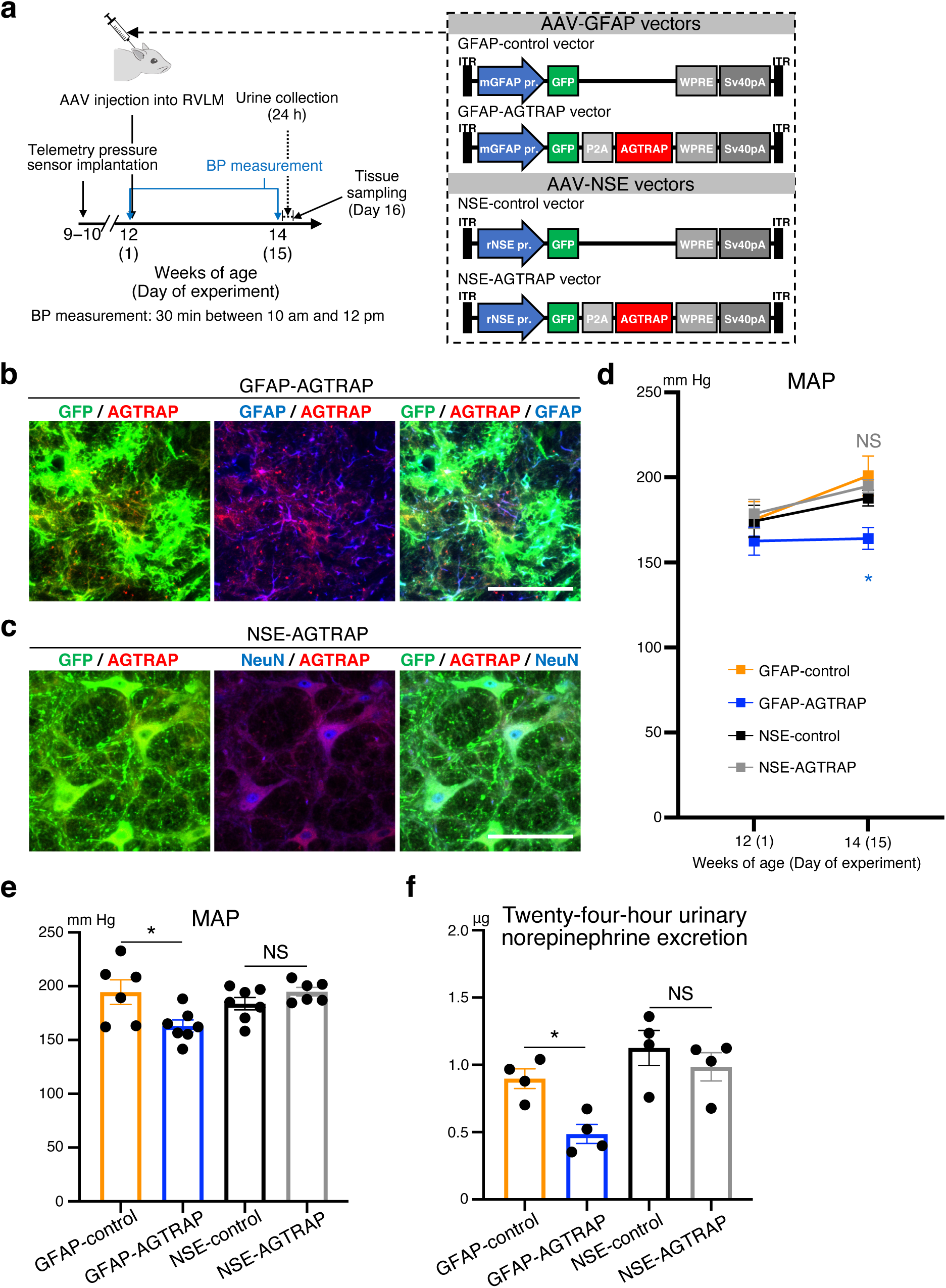
AAV-mediated expression of AGTRAP in RVLM astrocytes, but not neurons, lowers the BP in SHRSPs. **a,** Schematic representation of the experimental protocol to analyze the effects of AAV-mediated transduction of RVLM astrocytes or neurons with the AGTRAP gene. ITR; inverted terminal repeat. **b,c,** Astrocyte- **(b)** and neuron- **(c)** specific transgene expression by the RVLM injection of AAV9 vectors. Micrographic images of GFP (green) and anti-GFAP **(b)** or anti-NeuN **(c)** immunostaining (blue) of RVLM of SHRSPs 15 days after the injection of AAV9 vectors indicated at the top of each panel. Scale bars, 50 μm. Images are representative of three rats. **d−f,** Values just before (Day 1) and 2 weeks after (Day 15) the AAV injection into the RVLM **(d)**, and values on Day 15 **(e)** of MAP (**d**: *P* = 0.0222 for GFAP-control vs. GFAP-AGTRAP and *P* > 0.9999 for NSE-control vs. NSE-AGTRAP. **e**: *P* = 0.0229 for column 1 vs. 2, *P* = 0.6864 for column 3 vs. 4. *n* = 6 rats for GFAP-control; *n* = 7 rats for GFAP- AGTRAP; *n* = 7 rats for NSE-control; *n* = 6 rats for NSE-AGTRAP) and 24-h urinary norepinephrine excretion **(f)** (*P* = 0.0497 for column 1 vs. 2, *P* = 0.7455 for column 3 vs. 4. *n* = 4 rats for each group) of SHRSPs subjected to the RVLM injection of AAV9 vectors. Data are presented as mean ± s.e.m. \**P* < 0.05; NS, not significant, two-way repeated measures ANOVA with Bonferroni’s post hoc multiple comparisons test **(d)** or one-way ANOVA with Tukey’s post hoc multiple comparisons test **(e,f)**.

Furthermore, the AAV-mediated expression of AGTRAP in astrocytes, but not neurons, of SHRSPs’ bilateral RVLMs decreased the 24-h urinary norepinephrine excretion (Fig. 3f). Injection of the control AAV vector (GFAP-control or NSE-control) did not significantly affect the BP of SHRSPs (Extended Data Fig. 5f). These results support the importance of AT1R signal intensity in the RVLM astrocytes for SHRSPs’ development of hypertension and sympathetic hyperactivity, as well as the physiological relevance of the PHM-induced decrease in the AT1R expression we observed in the RVLM astrocytes of SHRSPs (Fig. 1j).

### PHM generates low-amplitude pressure waves and induces interstitial fluid movement in rat RVLM

We then sought to determine the physical effects that PHM produced in the rat RVLM. To do so, we analyzed local pressure changes using a telemetry pressure sensor (Fig. 4a) as we described previously^24^. PHM generated pressure waves (changes) with ∼1.2 mm Hg peak amplitude (Fig. 4b−d). Because the frequency of these PHM-induced pressure changes was equivalent to that of PHM (2 Hz), they were likely due to the local cyclical microdeformation generated upon the PHM. We then postulated an analogy to bone, an organ that only yields to minimal deformation. Because the function of bone is known to be modulated by interstitial fluid flow-derived shear stress on osteocytes^23^, we speculated that the interstitial fluid movement generated by the microdeformation-induced stress distribution changes in the brain might result in the shear stress-mediated regulation of nervous cell functions^24^.

**Fig. 4.**
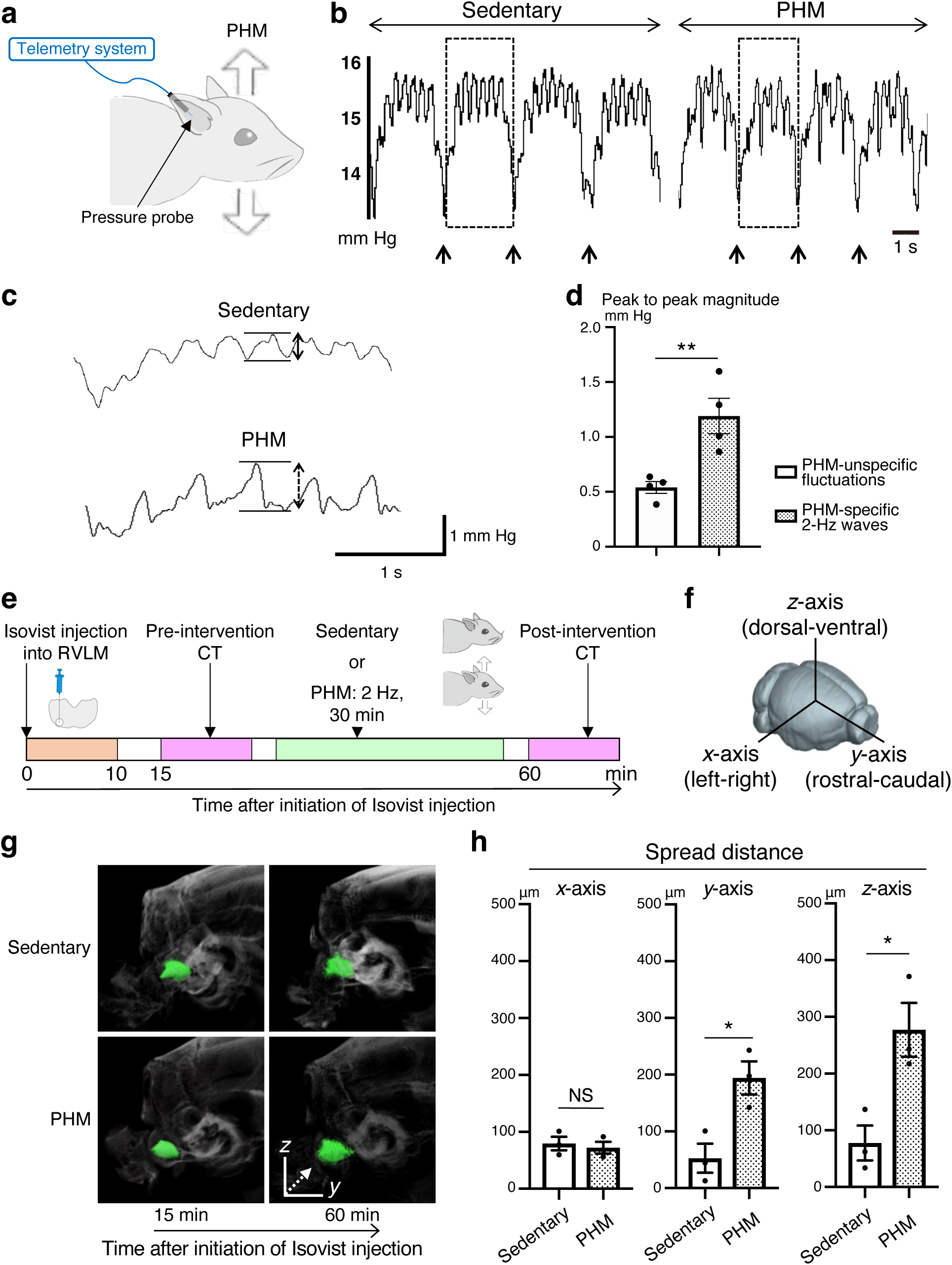
PHM generates pressure waves of low amplitude, and facilitates interstitial fluid movement (flow) in the rat RVLM. **a,** Schematic representation of the pressure measurement in the rat RVLM. **b,** Representative pressure waves recorded in the rat RVLM during the sedentary condition and PHM. Arrows point to the time of transition from inhalation to exhalation detected by simultaneous respiration monitoring. Scale bar, 1 s. Images are representative of four independent experiments with similar results. **c,** Respiration- unsynchronized pressure changes. Respiration-synchronized pressure waves indicated by dotted rectangles in **(b)** are presented with high magnification. Right-angled scale bar, 1 s / 1 mm Hg. Note that the 2-Hz pressure waves indicated by a two-headed dotted line arrow were specifically generated during PHM. **d,** Magnitude of PHM-specific and -unspecific pressure changes unsynchronized with respiration. The peak-to-peak magnitudes indicated by two-headed arrows in **(c)** were quantified (*P* = 0.0089. *n* = 4 rats for each group, 10 segments analyzed for each rat). **e,** Schematic representation of experimental protocol for the μCT analysis of Isovist injected into the rat RVLM. **f,** Definition of the *x*-(left-right), *y*-(rostral-caudal), and *z*-(dorsal-ventral) axes used in this study. **g,** Representative Isovist spread presented on X-ray images. Isovist clusters are indicated by green. Images are representative of three rats. A dotted line arrow indicates the main direction of spreading in this sample. **h,** Quantification of Isovist spread along each axis (left chart: *P* = 0.6666. middle chart: *P* = 0.0218. right chart: *P* = 0.0244. *n* = 3 rats for each group). Data are presented as mean ± s.e.m. \**P* < 0.05; \*\**P* < 0.01; NS, not significant, unpaired two-tailed Student’s *t-*test.

To analyze the PHM-induced interstitial fluid movement in the RVLM, we injected an iodine-based contrast agent (Isovist^®^) into the RVLM of anesthetized rats, and tracked its distribution with sequential micro-computed tomography (μCT) (Fig. 4e) as we previously did to analyze the movement of intramuscular interstitial fluid^34^. We found that PHM significantly promoted Isovist spreading in the rostral-caudal and dorsal-ventral (*y*- and *z*-axes, Fig. 4f) directions (Fig. 4g,h). In contrast, PHM did not significantly affect the left-right spreading (*x*- axis, Fig. 4f) of Isovist (Fig. 4g,h). From the extent of PHM-induced increase in Isovist spread, we estimated that the velocity of interstitial fluid movement in the rat RVLM was increased by approximately two- to three-times during PHM; however, precise evaluation was difficult because of differences in size and time scales between our μCT analysis (100-μm order, >30- min interval) and PHM effects at a cellular level (μm order, ∼0.5-s interval).

Our analyses using a multiphoton microscope (Extended Data Fig. 6a-c) and magnetic resonance (MR) imaging (MRI) (Extended Data Fig. 6d) indicated that the interstitial space of the rat RVLM is not randomly structured but oriented approximately along the centroidal line of this part of the brain as shown in Extended Data Fig. 6e. Furthermore, the cross-sectional area of the interstitial space was estimated to be 0.0083-0.18 μm^2^ (Extended Data Fig. 7a-d). We assume that PHM generates cyclical microdeformation in the rat RVLM (Extended Data Fig. 7e), thereby promoting interstitial fluid movement.

Integrating these findings with previous reports on the property^35^, flow velocity^36–38^, and occupancy^39^ of interstitial fluid in the brain, we calculated the average magnitude of interstitial fluid flow-derived shear stress exerted on rat RVLM cells during PHM (0.076−0.53 Pa; Supplementary Table 1).

### FSS on astrocytes decreases AT1R expression in vitro

We then sought to determine what type of mechanical force was responsible for the PHM-induced decrease in the AT1R expression in RVLM astrocytes (Fig. 1h,j) by in vitro experiments using cultured cells. Taking the approximate nature of our FSS calculation (Supplementary Table 1) into account, we extensively examined whether application of FSS or hydrostatic pressure change (HPC), another type of mechanical intervention, modulated AT1R expression. Based on our calculation (Supplementary Table 1), we applied pulsatile FSS with an average magnitude of 0.05−0.7 Pa to cultured primary astrocytes, which were prepared from the astrocyte-GFP mice^40^ (Extended Data Fig. 8a), using a system we previously reported^23, 24, 34^. Quantitative polymerase chain reaction (qPCR) analysis revealed that application of FSS with ≥0.3-Pa magnitude (0.5 Hz, 30 min) significantly decreased the AT1R expression in astrocytes for at least 24 h in an apparently magnitude-dependent manner (Fig. 5a and Extended Data Fig. 8b). In contrast, cyclical application of HPC, ranging from 1 to 40 mm Hg, nonsignificantly altered (≤10 mm Hg) or significantly increased (≥20 mm Hg) the AT1R expression in cultured astrocytes (Fig. 5b). Consistently, immunostaining (Fig. 5c,d) and fluorescently labeled ligand (Ang II) binding (Fig. 5e,f) analyses of cultured astrocytes indicated that the AT1R expression was significantly decreased by exposure to 30-min 0.7-Pa FSS. Collectively, FSS at magnitudes <1 Pa, but not HPC, on astrocytes decreases the AT1R expression in vitro. Notably, FSS application to Neuro2A cells, which exhibit neuronal phenotypes and morphology^41, 42^, did not decrease the AT1R expression (Extended Data Fig. 8c−e).

**Fig. 5.**
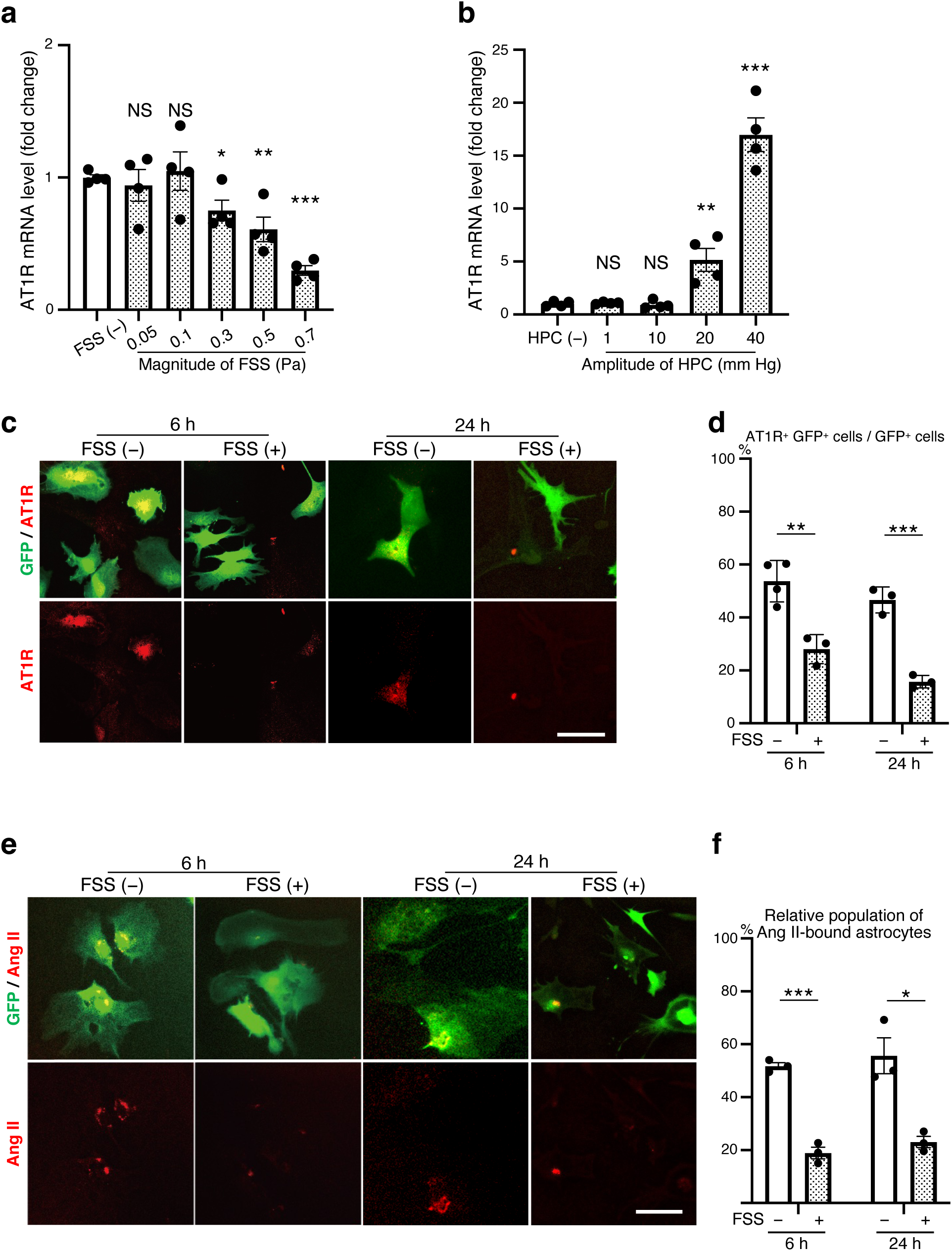
FSS on cultured astrocytes decreases their AT1R expression and Ang II-binding potential in vitro. **a–d,** AT1R expression in cultured astrocytes with or without exposure to FSS or HPC. Astrocytes prepared from the astrocyte-GFP mice, either left unexposed or exposed to pulsatile FSS (average 0.05−0.7 Pa, 0.5 Hz, 30 min) **(a)** or cyclical HPC (1−40 mm Hg, 0.5 Hz, 30 min) **(b)** were solubilized 24 h after the termination of intervention, and subjected to qPCR analysis. AT1R mRNA expression levels were normalized against GAPDH expression and scaled as the mean value from control samples (cells left unexposed to FSS or HPC) set as 1 (**a**: *P* = 0.6453 for 0.05 Pa, *P* = 0.7517 for 0.1 Pa, *P* = 0.0226 for 0.3 Pa, *P* = 0.0064 for 0.5 Pa, *P* < 0.0001 for 0.7 Pa. *n* = 4. **b**: *P* = 0.5592 for 1 mm Hg, *P* = 0.7113 for 10 mm Hg, *P* = 0.0088 for 20 mm Hg, *P* < 0.0001 for 40 mm Hg. *n* = 4). **(c)** Microscopic images of anti-AT1R (red) and anti-GFP (green) immunostaining of cultured astrocytes, either left unexposed or exposed to pulsatile FSS (0.7 Pa, 0.5 Hz, 30 min) and fixed 6 or 24 h after the intervention. Images are representative of three or four independent experiments with similar results. Scale bar, 50 μm. **(d)** Relative population of AT1R/GFP/ double positive (AT1R^+^ GFP^+^) cells were quantified as a ratio to total GFP-positive (GFP^+^) cells in each sample [*P* = 0.0049 for 6 h, *P* = 0. 0006 for 24 h. More than 100 GFP-positive cells were analyzed in each sample. *n* = 4 for 6 h FSS (−); *n* = 3 for the other groups]. **e,f,** Effect of FSS on astrocytes’ Ang II-binding potential. Cultured astrocytes were either left unexposed or exposed to FSS as in **(a,c−d)**. Six or twenty-four hours after the cessation of 30-min FSS application (0.7 Pa, 0.5 Hz), cells were subjected to fluorescent Ang II binding assay. **(e)** Microscopic images representative of three independent experiments with similar results. Scale bar, 50 μm. **(f)** GFP-positive cells with punctate red fluorescence (TAMRA-Ang II-bound astrocytes) were quantified as a ratio (%) to total GFP-positive cells in each sample (*P* = 0.0002 for 6 h, *P* = 0.0104 for 24 h. One-hundred GFP-positive cells were analyzed in each sample. *n* = 3 for each group). Data are presented as mean ± s.e.m. \**P* < 0.05, \*\**P* < 0.01, \*\*\**P* < 0.001; NS, not significant; unpaired two-tailed Student’s *t-*test.

The duration (>24 h) of FSS effects on the AT1R expression in astrocytes (Fig. 5) poses a possibility of cumulative effects of FSS applied repeatedly at 24-h intervals. Nevertheless, 2- day PHM (30 min/day) alleviated the pressor and depressor responses to Ang II and AT1R blocker, respectively, injected into the RVLM in SHRSPs (Extended Data Fig. 9a−e), supporting the relevance of the relatively quick decrease in AT1R expression in our FSS experiments to our in vivo observations. Taken together, our in vitro findings are consistent with the notion that the FSS-mediated persistent reduction of the AT1R expression is involved in the effects of daily PHM application on BP (Fig. 1b,c) and AT1R expression in the RVLM astrocytes (Fig. 1h,j) in SHRSPs. In contrast to PHM (Fig. 1b) and treadmill running^25^, both of which required >2 weeks to decrease the BP in hypertensive rats, daily AT1R blocker administration has been reported to decrease the BP of hypertensive rats in less than one week^43, 44^. The decrease in the AT1R signaling in RVLM astrocytes may take considerably longer time to elicit its consequences on cardiovascular variables as compared with the systemic RAS blockade^45^. Relatedly, 4-week PHM in SHRSPs initiated in the plateau phase of their hypertension development (age of 21 weeks) did not significantly alter BP (Extended Data Fig. 10a−c), although it did decrease the AT1R expression in RVLM astrocytes, but not neurons (Extended Data Fig. 10d−f). These findings imply the existence of a complex mechanism that links AT1R signaling in RVLM astrocytes with BP regulation, which can be irreversibly modified depending on various factors. For example, both vascular and renal functions in SHRSPs have been reported to be impaired in association with aging (≥16 weeks) and severity of hypertension (mean arterial pressure [MAP] ≥200 mm Hg)^46–49^.

### Hindrance of interstitial fluid movement by hydrogel introduction in the RVLM eliminates the ability of PHM to decrease the AT1R expression in RVLM astrocytes and the BP in SHRSPs

To examine whether the interstitial fluid movement in the RVLM mediated the effects of PHM on the BP and AT1R expression in the RVLM astrocytes of SHRSPs, we modulated the local interstitial fluid dynamics. Following the procedure we used to restrict the interstitial fluid movement in the mouse PFC^24^, we hindered the interstitial fluid movement in situ by microinjecting mutually reactive polyethylene glycol (PEG) gel-precursor (pre-gel) solutions into the rat RVLM (Fig. 6a). Injected pre-gel solution spread over the rat RVLM, leading to hydrogel formation in the interstitial space in situ (Extended Data Fig. 11a). We previously showed that hydrogel introduction only hinders the fluid movement but does not restrict the diffusion of small molecules inside the gel^24, 50^. Consistent with this, hydrogel introduction did not delay or attenuate the pressor and depressor responses to Ang II and Ang II antagonist, respectively, injected into the RVLM (Extended Data Fig. 11b−f), indicating rapid solute diffusivity through the hydrogel.

**Fig. 6.**
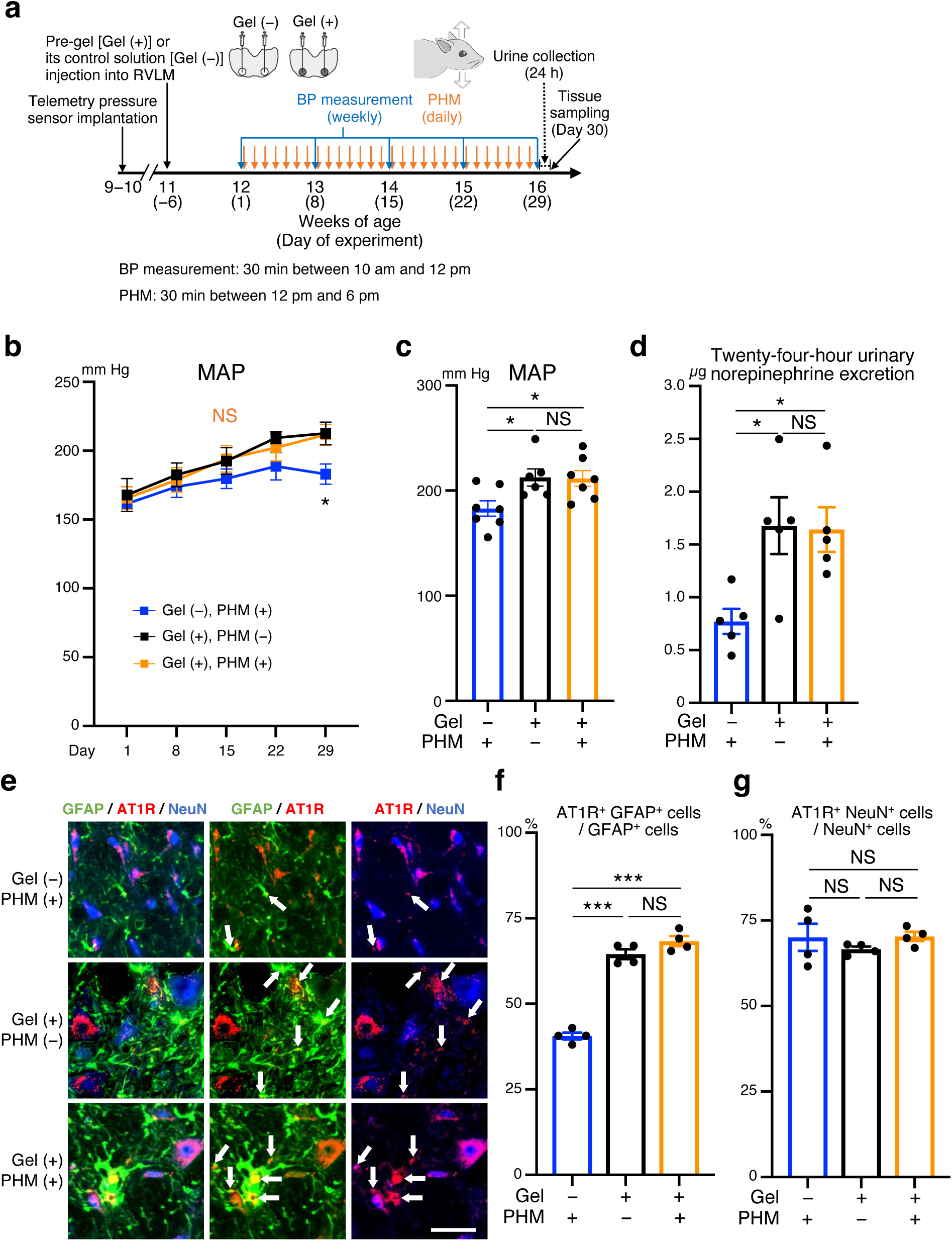
Hydrogel introduction eliminates the decreasing effects of PHM on BP and AT1R expression in the RVLM astrocytes in SHRSPs. **a,** Schematic representation of the experimental protocol to analyze the effects of PHM with and without PEG hydrogel introduction in the bilateral RVLMs of SHRSPs. PHM was applied daily for consecutive 28 days. **b−d,** Time courses **(b)** and values on Day 29 **(c)** of MAP, and 24-h urinary norepinephrine excretion **(d)** of SHRSPs, subjected to various combinations of daily PHM application and hydrogel introduction in the bilateral RVLMs. Note the absence of significant difference in the BP **(b,c)** and urinary norepinephrine excretion **(d)** of SHRSPs with hydrogel introduced RVLMs [Gel (+)] between with and without PHM [**b**, blue vs. orange: *P* = 0.4314 for Day 15, *P* = 0.4685 for Day 22, *P* = 0.0389 for Day 29; blue vs. black: *P* = 0.5372 for Day 15, *P* = 0.2000 for Day 22, *P* = 0.0406 for Day 29; black vs. orange: *P* = 0.9911 for Day 15, *P* = 0.8222 for Day 22, *P* = 0.9965 for Day 29. **c**: *P* = 0.0387 for column 1 vs. 2, *P* = 0.0372 for column 1 vs. 3, *P* = 0.9959 for column 2 vs. 3. *n* = 7 rats for Gel (−), PHM (+) and Gel (+), PHM (+); *n* = 6 rats for Gel (+), PHM (−); **d**: *P* = 0.0247 for column 1 vs. 2, *P* = 0.0307 for column 1 vs. 3, *P* = 0.9920 for column 2 vs. 3. *n* = 5 rats for each group]. **e**, Micrographic images of anti-GFAP (green), anti-AT1R (red), and anti-NeuN (blue) immunostaining of the RVLM in SHRSPs subjected to various combinations of hydrogel introduction in the bilateral RVLMs and 4-week PHM application. Arrows point to anti-AT1R immunosignals that overlap with anti-GFAP, but not anti-NeuN, immunosignals in merged images. Scale bar, 50 μm. Images are representative of four rats. **f,g,** Quantification of AT1R-positive astrocytes **(f)** and neurons **(g)** in the RVLM. Note the absence of significant difference in the ratio of AT1R-positive astrocytes of SHRSPs with hydrogel introduced RVLMs [Gel (+)] between with and without PHM (**f**, columns 2 and 3). Fifty NeuN^+^ cells and 100 GFAP^+^ cells were analyzed for each rat (**f**: *P* < 0.0001 for column 1 vs. 2 and column 1 vs. 3, *P* = 0.1597 for column 2 vs. 3. **g**: *P* = 0.6029 for column 1 vs. 2, *P* = 0.9963 for column 1 vs. 3, *P* = 0.5552 for column 2 vs. 3. *n* = 4 rats for each group). Data are presented as mean ± s.e.m. \**P* < 0.05; \*\*\**P* < 0.001; NS, not significant; two-way repeated measures ANOVA **(b)** or one-way ANOVA **(c,d,f,g)** with Tukey’s post hoc multiple comparisons test.

Hydrogel introduction in the bilateral RVLMs eliminated the ability of PHM to decrease the BP (Fig. 6b, black and orange lines; Fig. 6c, columns 2 and 3), urinary norepinephrine excretion (Fig. 6d, columns 2 and 3), and AT1R expression in the RVLM astrocytes (Fig. 6e, rows 2 and 3; Fig. 6f, columns 2 and 3) in SHRSPs. In contrast, hydrogel introduction increased the BP (Fig. 6b, blue and orange lines; Fig. 6c, columns 1 and 3), norepinephrine excretion (Fig. 6d, columns 1 and 3), and AT1R expression in the RVLM astrocytes (Fig. 6e, rows 1 and 3; Fig. 6f, columns 1 and 3) of SHRSPs subjected to PHM. The AT1R expression in the RVLM neurons of SHRSPs remained unaltered irrespective of the combination of PHM and hydrogel introduction (Fig. 6e,g). These results suggest that hydrogel introduction in the RVLM disrupts the mechanism mediating the PHM-induced decrease in BP, norepinephrine excretion, and AT1R expression in the RVLM astrocytes of SHRSPs. In line with this, the antihypertensive effect of daily treadmill running in SHRSPs was eliminated by hydrogel introduction in the RVLM (Extended Data Fig. 12a−c), supporting our hypothesis about the mechanism underlying the antihypertensive effects of exercise.

As was the case with the mouse PFC^24^, hydrogel introduction (Extended Data Fig. 13a) did not affect the overall cell number or apoptosis (Extended Data Fig. 13b,c), survival or apoptosis of the RVLM astrocytes (Extended Data Fig. 13d,e) and neurons (Extended Data Fig. 13f,g), and the expression of pro-inflammatory cytokines (TNF-α and IL-1β) (Extended Data Fig. 13h) in the RVLM. Furthermore, hydrogel introduction in the rat RVLM did not significantly alter the intramedullary pressure (Extended Data Fig. 13i). Collectively, the loss of the PHM effects by hydrogel introduction in the bilateral RVLMs of SHRSPs (Fig. 6b−f) is likely to result from the hydrogel-mediated alteration in interstitial fluid dynamics, rather than the decreased cell viability and/or enhanced inflammatory responses caused by the impaired nutrient supply, removal of metabolic wastes, or persistent PEG existence/contact.

Consistent with these results supporting the importance of interstitial fluid movement (flow) in the RVLM of SHRSPs, both their BP and HR remained unchanged during the transition from before to after the initiation of PHM (Extended Data Fig. 14), precluding the response of baroreceptors, either carotid or aortic, to PHM. Furthermore, the activity of the aortic depressor nerve, which transmits afferent signals from baroreceptors and chemoreceptors located in the aortic arch^51^, also remained unaltered from before to after PHM initiation (Extended Data Fig. 14). Therefore, the baroreceptor response does not appear to be responsible for the antihypertensive effects of PHM on SHRSPs.

### Vertically oscillating chair riding (VOCR) lowers BP in hypertensive adult humans

The results from our animal experiments reveal the antihypertensive effect of the mechanical accelerations generated in the head during treadmill running at a moderate velocity. This prompted us to test whether application of mechanical intervention to the head lowered BP in hypertensive humans. We observed light jogging or fast walking (locomotion at the velocity of 7 km/h) typically produce ∼2-Hz vertical acceleration waves with an amplitude of ∼1.0 × *g* in the person’s head (Extended Data Fig. 15a, top). It seemed difficult to apply vertical forces only to the heads of human subjects safely without leading to considerable discomfort and distress. Therefore, we constructed a chair that could vertically oscillate at a frequency of 2 Hz (Extended Data Fig. 15b) and produce ∼1.0 × *g* acceleration waves in the head of the occupant (Extended Data Fig. 15a, bottom), although the other body parts were also subjected to cyclical vertical movements in this system. The waveform of the vertical acceleration (Extended Data Fig. 15a, bottom) was also determined considering the subject’s comfort in addition to technical issues.

Given that previous reports regarding antihypertensive effects of aerobic exercise typically recommend ≥3−4 days per week (frequency) and ≥30 min per session or day (duration)^2^, we set our regimen of VOCR as 3 days/week (Monday, Wednesday, and Friday unless needed to assign otherwise for particular reasons such as public holidays) and 30 min/day. Our pilot study following protocol 1, in which we simply compared the subjects’ BP and HR before and after 4-week (12 times) VOCR (Extended Data Fig. 15c), showed that VOCR decreased the BP in hypertensive humans (Extended Data Fig. 15d).

We then conducted a human study of protocol 2, in which we followed the changes in subjects’ BP and HR more minutely (Extended Data Fig. 16a). Encouraged by the positive results from the study of protocol 1, we adopted the same VOCR regimen as to its frequency (3 days/week) and duration (30 min/day). To detect the trends of the BP and HR changes more reliably by reducing the influences from their interday variabilities, we followed and analyzed “value of the week”s (see Methods). Participants were subjected to serial blood sampling to measure the plasma catecholamines (epinephrine, norepinephrine, and dopamine) and renin activity, and serum aldosterone and C-reactive protein (CRP) before and after the intervention period (Extended Data Fig. 16a). To conduct the second blood sampling on the next day of the last bout of VOCR, the intervention period was extended from four weeks (total 12 times, typically 26 days) to 4.5 weeks (total 14 times, 30−31 days) because blood sampling could not be done during weekends at our hospital. Systolic BP (SBP), diastolic BP (DBP), and MAP (“value of the week”s) immediately after the intervention period significantly decreased as compared with those immediately before the intervention period (Fig. 7a). Notably, the post-intervention follow-up showed that the BP-lowering effect apparently persisted for four weeks, but not five weeks, after the last bout of VOCR (Fig. 7b). Similar to our animal study, we did not observe significant changes in the HR by the VOCR intervention (Fig. 7a,b and Extended Data Fig. 15d). Significant differences were not detected in the blood levels of catecholamines, aldosterone, renin activity, and CRP between before and after the VOCR intervention (Extended Data Fig. 16b).

**Fig. 7.**
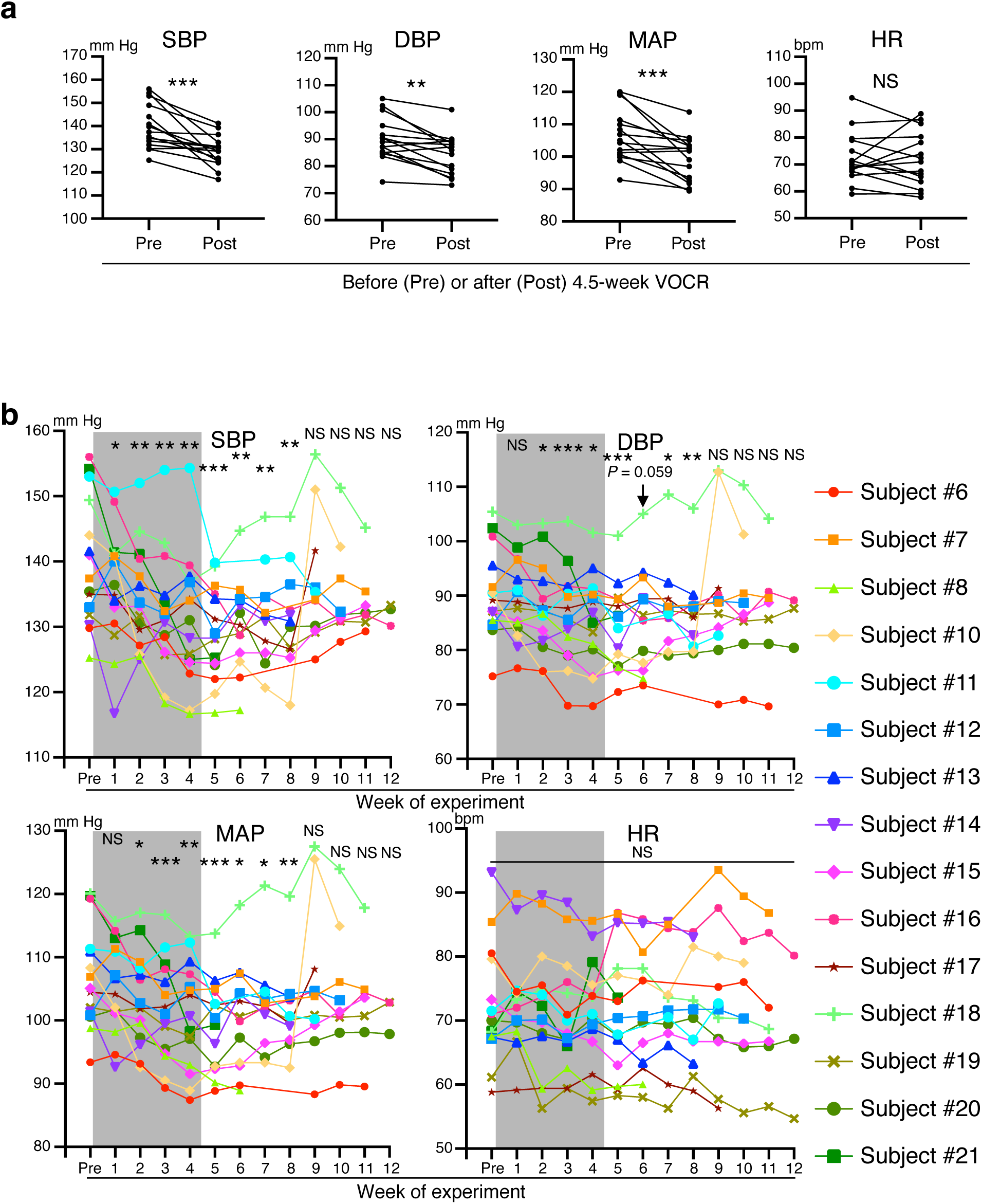
VOCR has an antihypertensive effect on hypertensive adult humans. **a,** BP and HR “value of the week”s immediately before and after 4.5-week VOCR in the study of protocol 2 (SBP: *P* = 0.0005. DBP: *P* = 0.0011. MAP: *P* = 0.0008. HR: *P* = 0.7845. *n* = 15). **b,** Subject number-corresponding trajectories and statistical analysis of BP and HR in the study of protocol Gray-shaded rectangles indicate the VOCR intervention periods (4.5 weeks). The colors and symbols of individual lines correspond to those of individual subject numbers listed on the right, excluding subject 9 (see Supplementary Table 2). Each “value of the week” was statistically compared with that of the week immediately before the initiation of VOCR intervention (SBP: *P* = 0.0293 for Pre vs. Week 1, *P* = 0.0028 for Pre vs. Week 2, *P* = 0.0013 for Pre vs. Week 3, *P* = 0.0035 for Pre vs. Week 4, *P* = 0.0002 for Pre vs. Week 5, *P* = 0.0078 for Pre vs. Week 6, *P* = 0.0035 for Pre vs. Week 7, *P* = 0.0075 for Pre vs. Week 8, *P* = 0.2132 for Pre vs. Week 9, *P* = 0.1314 for Pre vs. Week 10, *P* = 0.0973 for Pre vs. Week 11, *P* = 0.3993 for Pre vs. Week 12. DBP: *P* = 0.3022 for Pre vs. Week 1, *P* = 0.0436 for Pre vs. Week 2, *P* = 0.0010 for Pre vs. Week 3, *P* = 0.0100 for Pre vs. Week 4, *P* = 0.0006 for Pre vs. Week 5, *P* = 0.0599 for Pre vs. Week 6, *P* = 0.0488 for Pre vs. Week 7, *P* = 0.0096 for Pre vs. Week 8, *P* = 0.9346 for Pre vs. Week 9, *P* = 0.7850 for Pre vs. Week 10, *P* = 0.0769 for Pre vs. Week 11, and *P* = 0.3137 for Pre vs. Week 12. MAP: *P* = 0.1075 for Pre vs. Week 1, *P* = 0.0132 for Pre vs. Week 2, *P* = 0.0008 for Pre vs. Week 3, *P* = 0.0063 for Pre vs. Week 4, *P* = 0.0003 for Pre vs. Week 5, *P* = 0.0251 for Pre vs. Week 6, *P* = 0.0136 for Pre vs. Week 7, *P* = 0.0071 for Pre vs. Week 8, *P* = 0.6795 for Pre vs. Week 9, *P* = 0.4295 for Pre vs. Week 10, *P* = 0.0704 for Pre vs. Week 11, *P* = 0.3662 for Pre vs. Week 12. HR: *P* = 0.6287 for Pre vs. Week 1, *P* = 0.7840 for Pre vs. Week 2, *P* = 0.1573 for Pre vs. Week 3, *P* = 0.5380 for Pre vs. Week 4, *P* = 0.7331 for Pre vs. Week 5, *P* = 0.6995 for Pre vs. Week 6, *P* = 0.9110 for Pre vs. Week 7, *P* = 0.9875 for Pre vs. Week 8, *P* = 0.5866 for Pre vs. Week 9, *P* = 0.9566 for Pre vs. Week 10, *P* = 0.6487 for Pre vs. Week 11, *P* = 0.9905 for Pre vs. Week 12. *n* = 15 for Pre and Weeks 1 to 5; *n* = 13 for Week 6; *n* = 12 for Week 7; *n* = 11 for Weeks 8 and 9; *n* = 9 for Week 10; *n* = 7 for Week 11; *n* = 3 for Week 12). \**P* < 0.05; \*\**P* < 0.01; \*\*\**P* < 0.001; NS, not significant; paired two-tailed Student’s *t-*test.

The antihypertensive effect of VOCR observed in the participants of protocols 1 and 2 (Supplementary Table 2) prompted us to proceed to protocol 3 in which we examined whether non-oscillating chair riding (NOCR), a control for VOCR, affected the BP of hypertensive adult humans (Extended Data Fig. 17a). We asked the participants in protocol 3 for their agreement to the continuous beat-by-beat BP and inter-beat (R−R) interval (RRI) recording. In those who agreed to these measurements, we analyzed the SBP and RRI variabilities (see Methods).

Whereas 4.5-week NOCR did not significantly affect participants’ BP (Fig. 8a, top), VOCR significantly lowered it (Fig. 8a, bottom), as in the studies following protocols 1 and 2. NOCR did not significantly alter the low-frequency (LF) power in SBP variability (Fig. 8b) or the ratio of LF/high frequency (HF) power (LF/HF ratio) in RRI variability (Fig. 8c). In contrast, VOCR significantly decreased the former (Fig. 8b) and elicited a decreasing tendency in the latter (Fig. 8c). These findings indicate that VOCR decreases the vascular sympathetic nerve activity^52, 53^ and its dominance over the cardiac parasympathetic activity^52, 54^; nevertheless, there remains some controversy regarding the use of the LF/HF ratio in RRI variability as an appropriate relevant indicator^55, 56^.

**Fig. 8.**
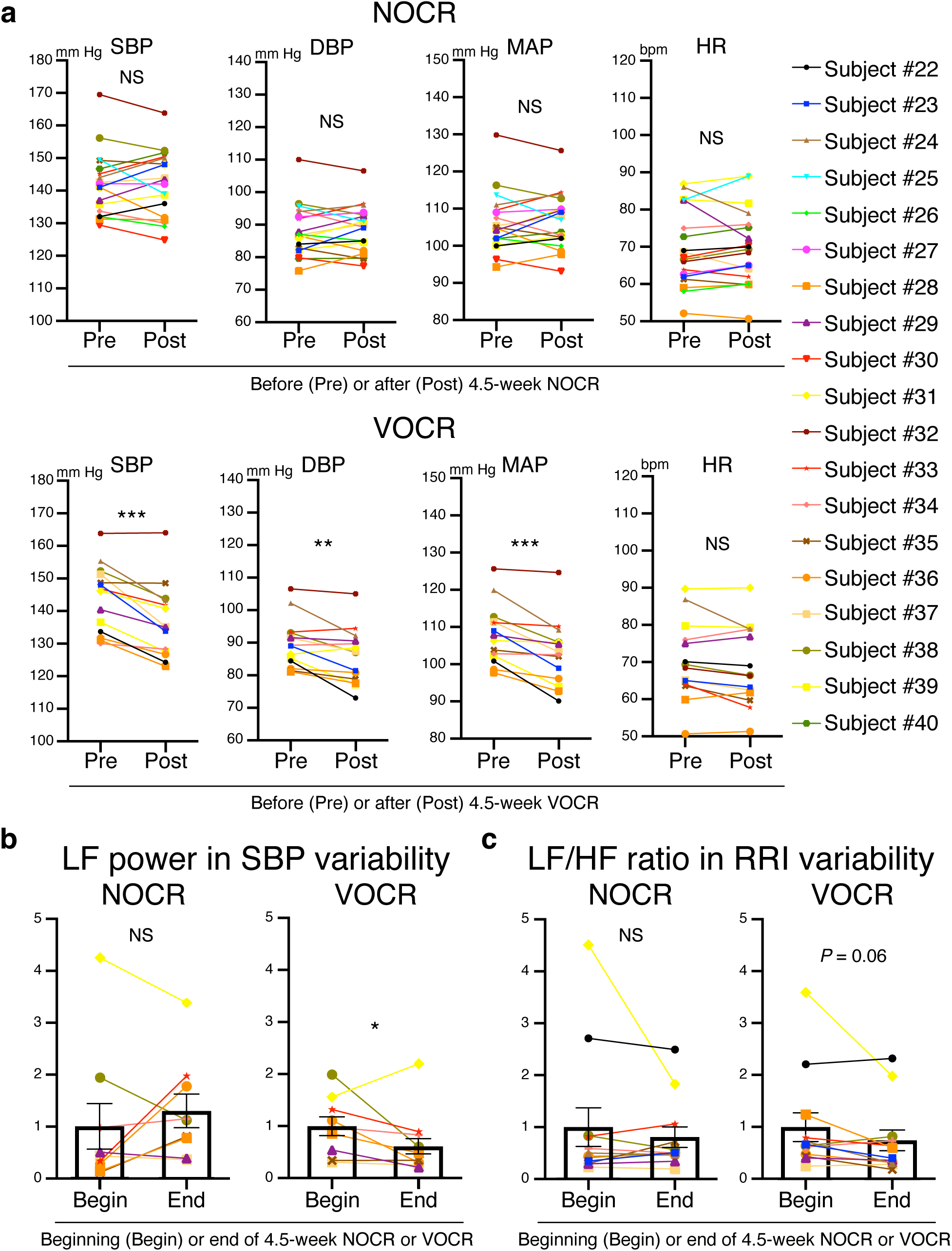
VOCR, but not NOCR, has an antihypertensive and sympathoinhibitory effect in hypertensive adult humans. **a,** BP and HR “value of the week”s immediately before and after 4.5-week NOCR (top) and VOCR (bottom) in the study of protocol 3 (NOCR, SBP: *P* = 0.9148. DBP: *P* = 0.6597. MAP: *P* = 0.7502. HR: *P* = 0.9002. *n* = 19; VOCR, SBP: *P* = 0.0001. DBP: *P* = 0.0051. MAP: *P* = 0.0006. HR: *P* = 0.0867. *n* = 14). **b,c,** LF power in SBP variability **(b)** and LF/HF ratio in RRI variability **(c)** at the beginning (Begin) and ending (End) periods of intervention (NOCR and VOCR) scaled as the mean value from “Begin” (left column in each graph) set as 1 (**b**, NOCR: *P* = 0.492, *n* = 10; VOCR: *P* = 0.016, *n* = 12. **c**, NOCR: *P* = 0.969, *n* = 12; VOCR: *P* = 0.063, *n* = 12). Data are presented as mean ± s.e.m. \**P* < 0.05; \*\**P* < 0.01; \*\*\**P* < 0.001; NS, not significant; paired two-tailed Student’s *t-*test (**a**) or Wilcoxon signed-rank test (**b,c**).

Collectively, our human studies suggest that VOCR, which reproduces mechanical accelerations in the head during light jogging or fast walking, has an antihypertensive and sympathoinhibitory effect in hypertensive humans. Although our animal studies were only conducted in male rats, VOCR had antihypertensive effects in both male and female humans (Extended Data Fig 17b). Importantly, in none of 33 VOCR subjects (Supplementary Table 2 and 3), apparent adverse events, including motion sickness and low back pain, were observed or manifested in relation to the VOCR intervention.

## Discussion

The antihypertensive effect of physical exercise has been shown to involve normalization of sympathetic hyperactivity in the brain^57^. However, it is still unclear whether exercise directly modulates the brain function. In this study, PHM, which reproduced mechanical accelerations generated in the head during treadmill running, allowed us to dissect bodily activity-derived physical effects.

Whereas AT1R signaling in both neurons and astrocytes in the RVLM have been reported to be involved in regulating BP^58, 59^, we observed that the AT1R expression in RVLM astrocytes was increased in SHRSPs as compared to that in WKY rats (Fig. 1j). In contrast, the AT1R expression in RVLM neurons was comparable between WKY rats and SHRSPs (Fig. 1i), although the AT1R expression in RVLM neurons has been shown to play an important role in other animal model(s) of hypertension^58^. Together with the decreases in the BP and urinary epinephrine excretion of SHRSPs in which RVLM astrocytes were transduced with the AGTRAP gene (Fig. 3d−f), the intensity of AT1R signaling in RVLM astrocytes appears to be critically involved in the pathogenesis of hypertension and sympathetic hyperactivity in SHRSPs.

Four-week PHM decreased the urinary norepinephrine excretion and AT1R expression in RVLM astrocytes of SHRSPs to the levels almost equivalent to those of WKY rats (Fig. 1f,j). However, PHM only partially alleviated the development of hypertension in SHRSPs (Fig. 1b,c) to the extent similar to the antihypertensive effects of treadmill running previously reported^25, 60^ or observed in this study (Extended Data Fig. 12b). Therefore, it is evident that factors other than the AT1R signaling in RVLM astrocytes also contribute to the pathogenesis of essential hypertension.

AT1R expression in the cultured astrocytes decreased upon FSS application (Fig. 5a,c−f). This was consistent with our findings that PHM and treadmill running decreased the AT1R expression in the RVLM astrocytes of SHRSPs (Fig. 1j and Extended Data Fig. 4c). However, the AT1R expression level in the RVLM astrocytes was low in WKY rats even without PHM (Fig. 1j), and this may raise a concern regarding the physiological relevance of our in vitro FSS experiments using cultured astrocytes. Still, it has been reported that cultured astrocytes typically exhibit increased “reactivity”, and do not fully recapitulate the physiological astrocytes in vivo^61^. We suggest that the FSS-induced decrease in the AT1R expression in cultured astrocytes we observed represents the physiological functions of astrocytes, despite that their increased basal AT1R expression may relate to the unphysiological aspects of two-dimensional (2-D) culture on stiff substrates (culture plastics). Cells in static culture are exposed to a complete absence of FSS, which may not be physiologically realized in vivo. Previous reports describe increased extracellular fluid in the brains of hypertensive humans^62^ and altered dynamics of the intracerebral interstitial fluid of SHRs^63^. Aberrant regulation of RVLM astrocytes’ function that relates to altered interstitial fluid movement-derived FSS may underlie the pathogenesis of essential hypertension.

PHM did not significantly alter the AT1R expression in SHRSPs’ RVLM neurons (Fig. 1i), and FSS did not decrease the AT1R expression in cultured Neuro2A cells (Extended Data Fig. 8c−e). However, we do not suspect that these results represent the absence of sensitivity of neurons to FSS or other type(s) of mechanical stimulation, particularly because we observed the PHM- and FSS-induced internalization of 5-HT_2A_ receptor expressed in mouse PFC neurons and Neuro2A cells, respectively^24^. Alternatively, we speculate that PHM and FSS may mitigate the hyperexpression of AT1R related to aforementioned pathological status of SHRSPs’ RVLM astrocytes or unphysiological nature of cultured astrocytes. In line with this notion, PHM did not significantly affect the AT1R expression in RVLM astrocytes in normotensive WKY rats (Fig. 1j).

Consistent with the lack of strict cell specificity in many of cellular responses to mechanical forces^64^, the FSS-induced decrease in AT1R expression, which was reported in vascular cells^19^, was also observed in cultured astrocytes. We speculate that there may be common homeostasis-regulatory mechanisms at the cellular level that involve fluid flow-derived shearing forces.

Based on our hypothesis concerning the similarity in the pathogenesis of high BP between human essential hypertension and SHRSPs, we conducted human studies in which we intended to reproduce the mechanical accelerations in the head that lowered the BP in SHRSPs. Although the mechanism behind the apparent antihypertensive effect of VOCR remains to be determined, the significant role of interstitial fluid dynamics in the RVLM, which we demonstrated by our animal experiments, might be shared between humans and rats or other animals (Extended Data Fig. 18). Whereas the plasma catecholamine levels were not significantly changed by the VOCR intervention in the human study of protocol 2 (Extended Data Fig. 16b), it is possible that the urinary epinephrine measures collected over 24 h in our rat PHM experiments (Fig. 1f, 3f, and 6d) enhanced our ability to capture the sympathetic nerve activity under ‘‘everyday-life’’ ambulatory conditions^65^. The findings from the study of protocol 3 suggest a sympathoinhibitory effect of VOCR (Fig. 8b,c).

Physical exercise is broadly useful to maintain the human health. Many of aerobic exercises, including walking and running, involve impact-generating bodily actions creating sharp accelerations in the head upon foot contacting with the ground. Furthermore, the antihypertensive effects of PHM in the rostral-caudal direction (Extended Data Fig. 2) or with a less peak magnitude (0.5 × *g*) or frequency (0.5 Hz) (Extended Data Fig. 3) may be relevant to the decrease in BP caused by other forms of exercise such as swimming and bicycle riding^66, 67^. We speculate that the beneficial effects of various types of exercises on a variety of brain function-related diseases and health disorders may rely at least partly on the modest changes in mechanical stress distribution in the brain, which may prompt optimal FSS on intracerebral nervous cells. To the contrary, alterations in interstitial fluid movement-derived shear stress may underlie the pathogenesis of various brain disorders, particularly those related to physical inactivity or aging.

### Limitation of study

There are several limitations of this study. We used SHRSP as an animal model of essential hypertension. There are several differences between SHRSP and human essential hypertension. SHRSPs develop hypertension in young adulthood, but not in middle age as in humans. Furthermore, SHRSPs cannot model environmental influences that trigger human hypertension, including increased salt intake, obesity, and physical inactivity. Nevertheless, SHRSP allows us to obtain a chronic stable hypertensive condition with minimal inter-individual variation, but without difficult or life-threatening technical interventions. We used only male rats for our animal studies, although sex is an important variable for nearly all diseases, including hypertension. This was because we intended to preclude or minimize the potential influence of estrogen and progesterone, both of which basically act protectively on cardiovascular systems, including the heart and endothelium. Perhaps for the same reason, in many or even most of animal experiments in previous studies investigating the pathogenesis of cardiovascular diseases, male animals have been used unless there is particular reason to analyze female animals. In particular, we intended to be consistent with previous studies in which the antihypertensive effects of treadmill running in male SHRs or SHRSPs were demonstrated^25, 60^. However, as we did not aim to investigate only male-specific matters, we included human participants of both sexes. Given that VOCR had antihypertensive effects in both male and female humans (Extended Data Fig. 17b), we anticipate that the mechanical regulation of RVLM astrocytes demonstrated in this study is not specific to males.

2-D cell culture experiments are non-physiologic particularly in light of 3-D microenvironments in vivo, and do not entirely recapitulate physiological conditions. Furthermore, our calculation of the FSS magnitude in vivo (Supplementary Table 1) is approximate. However, our extensive in vitro studies by testing FSS and HPC with various magnitudes/amplitudes (Fig. 5) supports the notion that FSS is responsible for the PHM-induced decrease in AT1R expression in rat RVLM astrocytes. Because of the easy detachment of mouse cerebral cortex- or hippocampus-derived primary neurons from the substrates by FSS, we tested Neuro2A cells, an alternative of cultured neuronal cells, which stably adhered to the substrates through FSS of magnitudes up to ∼1 Pa^24^.

We did not comprehensively analyze the effects of PHM on the brain functions, but focused on the study of RVLM. PHM may modulate the AT1R signaling in other brain regions that participate in the regulation of sympathetic nerve activity, including the anteroventral third ventricle, paraventricular nucleus of the hypothalamus, and nucleus tractus solitarii^6, 9^. It is technically difficult to specifically interfere solely with the interstitial flow in the brain or other tissues/organs in living animals. Nonetheless, hydrogel introduction in SHRSPs’ RVLM eliminated the decreasing effects of PHM on the BP and urinary norepinephrine excretion (Fig. 6b−d) as well as the antihypertensive effect of treadmill running (Extended Data Fig. 12), supporting the critical role for the RVLM. Because hydrogel may exert yet unknown effects, experiments of hydrogel introduction may not entirely prove the contribution of interstitial fluid movement. For example, hydrogel introduction may alter the stiffness and elasticity of extracellular matrix, which are known to affect the neurological physiology, pathology, and development^68^. Regardless, from the unaltered cell survival/apoptosis, pro-inflammatory protein expression, and pressure in the hydrogel-introduced RVLM of SHRSPs (Extended Data Fig. 13), massive detrimental or favorable processes are unlikely to be responsible. Given the involvement of reactive oxygen species (ROS) in hypertension in SHRSPs^69^ and the unaltered BP in hydrogel-introduced SHRSPs (Extended Data Fig. 12b), the immediate reduction of ROS by PEG itself^70^ also appears not to be responsible. Whereas further studies are required to strictly determine the specific role of interstitial fluid dynamics in the RVLM, our findings conform to the notion of its significance in BP control.

We conducted microinjection into the rat RVLM at a rate of 0.03−0.2 μL/min (see Methods), ∼4−20 times faster than that reportedly associated with excellent preservation of central nervous tissue^71^. Although microinjection of 0.1–0.2 μL/min has been successfully employed in animal (rat) brain research^72–75^, our approach may have compromised the brain region to some extent.

Unlike the case of PHM in rats, VOCR of humans generates vertical accelerations at various body parts in addition to the head. Therefore, we cannot preclude the possibility that the effects of VOCR also involves mechanical regulation of tissues and organs other than the brain. For example, the antihypertensive effect of human VOCR became significant in ∼2 weeks, approximately as quickly as or even a little more quickly than that of rat PHM (compare the MAP between SHRSPs in Fig. 1b and humans in Fig. 7b), despite the lower frequency of VOCR (PHM: 7 days/week; VOCR: 3 days/week). This could be attributed to some additional influence of cyclical vertical motion of body parts other than the head.

Our clinical studies are based on a small number of subjects (Supplementary Table 2 and 3) with a fixed condition (frequency of 2 Hz, peak acceleration of ∼1.0 × *g*, 30 min/day, 3 days/week, 12−14 rides), and need further analysis with a much larger sample size and varying conditions to determine the optimal VOCR.

The antihypertensive outcome of isometric exercise^76^ cannot be explained by direct mechanical effects on the brain; it is evidently mediated otherwise. Nevertheless, application of moderate exercise-mimicking mechanical intervention is expected to be highly safe with minimal possibility of adverse effects, providing a novel therapeutic/preventative strategy for physical disorders including those resistant to conventional treatments such as drug administration. Our approach utilizing mechanical interventions may bring considerable benefits to those who cannot receive benefits from exercise because of physical disabilities.

## Data availability

All data are included in this article and its supplementary information files. Raw data are available from the corresponding author upon reasonable request.

## Acknowledgements

We thank K. Nakanishi, K. Hamamoto, and N. Kume for their consistent support. This work was in part supported by Intramural Research Fund from the Japanese Ministry of Health, Labour and Welfare; Grants-in-Aid for Scientific Research from the Japan Society for the Promotion of Science (KAKENHI 15H01820, 15H04966, 18H04088, 20K21778, 21H04866 to Y.S.; 20K19367 to N.S.; 17H02127 and 18H03138 to T.O.; 19K06899 to A.K.); Brain Mapping by Integrated Neurotechnologies for Disease Studies (Brain/MINDS, JP20dm0207057 to H.H.) and Research and Development Grants for Comprehensive Research for Persons with Disabilities (21dk0310116j0201 to Y.S. and D.Y.) from the Japan Agency for Medical Research and Development (AMED); the JST MIRAI Project (to Y.S. and D.Y.); the Naito Science & Engineering Foundation (to Y.S.); the Uehara Memorial Foundation (to Y.S.).

## Author contributions

S.M. and N.S. conducted most of the animal/cell experiments. Y.S. and N.S. pursued the human studies. Y.S. conceived the research, designed the study, and led the project. K.S. and T.K. provided technical advice for the experiments involving measurement of cardiovascular variables. N.S., K.T., M.S. and Y.S. wrote the manuscript. T.M. and A.T. contributed to the design and construction of the machine for PHM. D.Y. helped the in vitro FSS experiments.

D.Y. and K.F. carried out calculation of in vivo FSS. T.S. and Y.Y. developed and provided the PEG hydrogel system. A.K. and H.H. prepared and provided the AAV vectors. S.Sat. and S.Sai. conducted the multiphoton microscopy and MRI experiments, respectively, and K.T. analyzed the images. K.Y. analyzed the data from human continuous BP and RRI measurements. K.T., T.K., M.A., H.I., Y.M., and T.O. supported Y.S. in the human studies. K.S., T.S., S.T. M.S., T.O., H.O., and M.N. provided technical, advisory and financial support.

---

**Extended Data Fig. 1.**
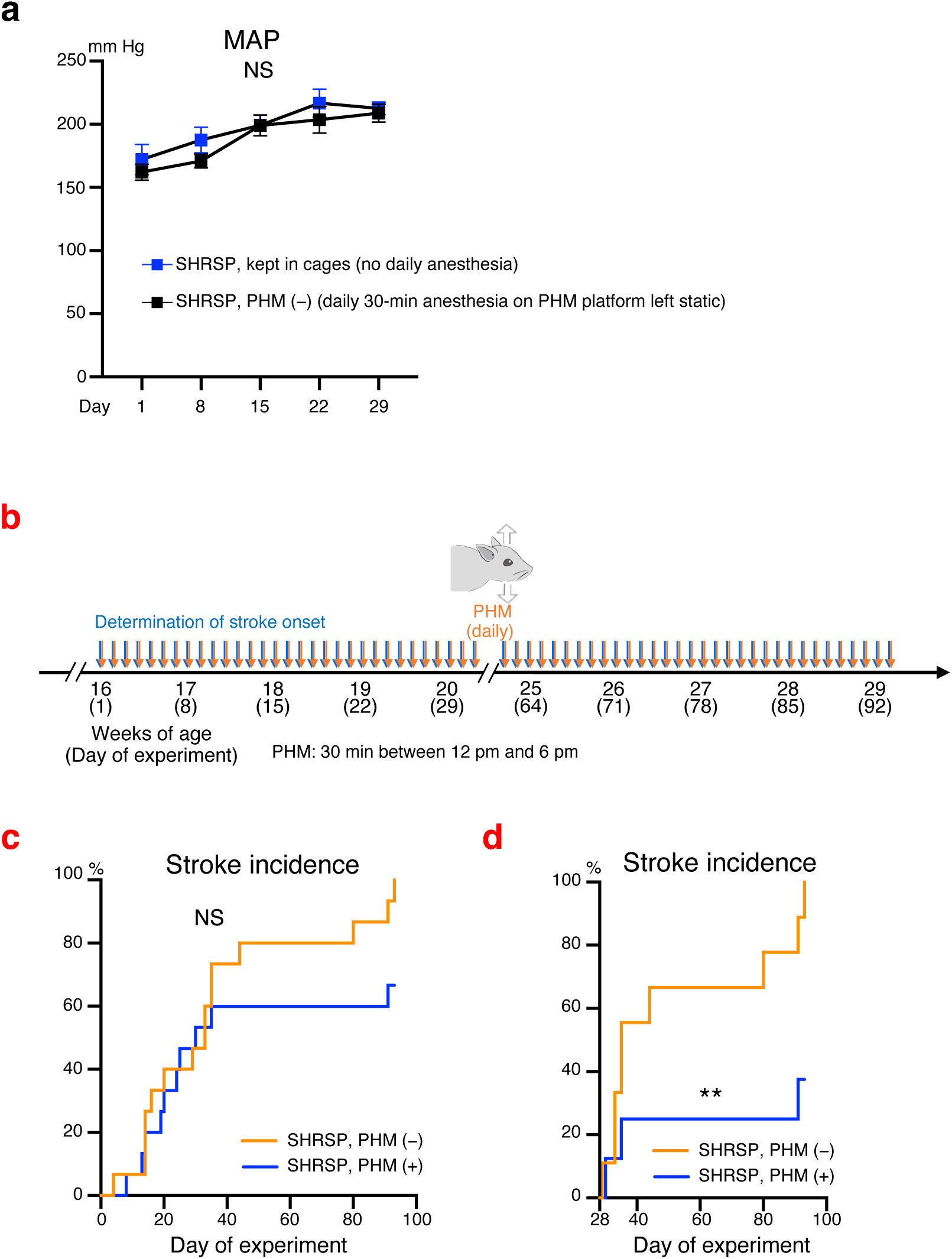
Daily anesthesia alone does not affect BP, and ≥4 weeks of daily PHM decreases or delays the stroke incidence in SHRSPs. **a,** Time courses of the MAP of SHRSPs either routinely kept in cages (no anesthesia) or subjected to daily anesthesia on the platform of PHM machine without turning on its switch [PHM (−)] [*P* > 0.9999 for Day 15, Day 22, and Day 29. *n* = 6 rats for no daily anesthesia; *n* = 8 rats for PHM (−)]. **b,** Schematic representation of the experimental protocol to analyze the effects of PHM on the stroke incidence in SHRSPs. **c,d,** Kaplan-Meier curve of the stroke incidence in SHRSPs with and without PHM represented as PHM (+) and PHM (−), respectively. Data from SHRSPs that did not have a stroke during the first four weeks after the initiation of intervention (daily PHM or anesthesia alone) is shown in **(d)** [**c**: *P* = 0.1179. *n* = 15 rats for PHM (−); *n* = 15 rats for PHM (+). **d**: *P* = 0.0093. *n* = 9 rats for PHM (−); *n* = 8 rats for PHM (+)]. \*\**P* < 0.01; NS, not significant; two-way repeated measures ANOVA with Bonferroni’s post hoc multiple comparisons test **(a)** or long-rank test **(c,d)**.

**Extended Data Fig. 2.**
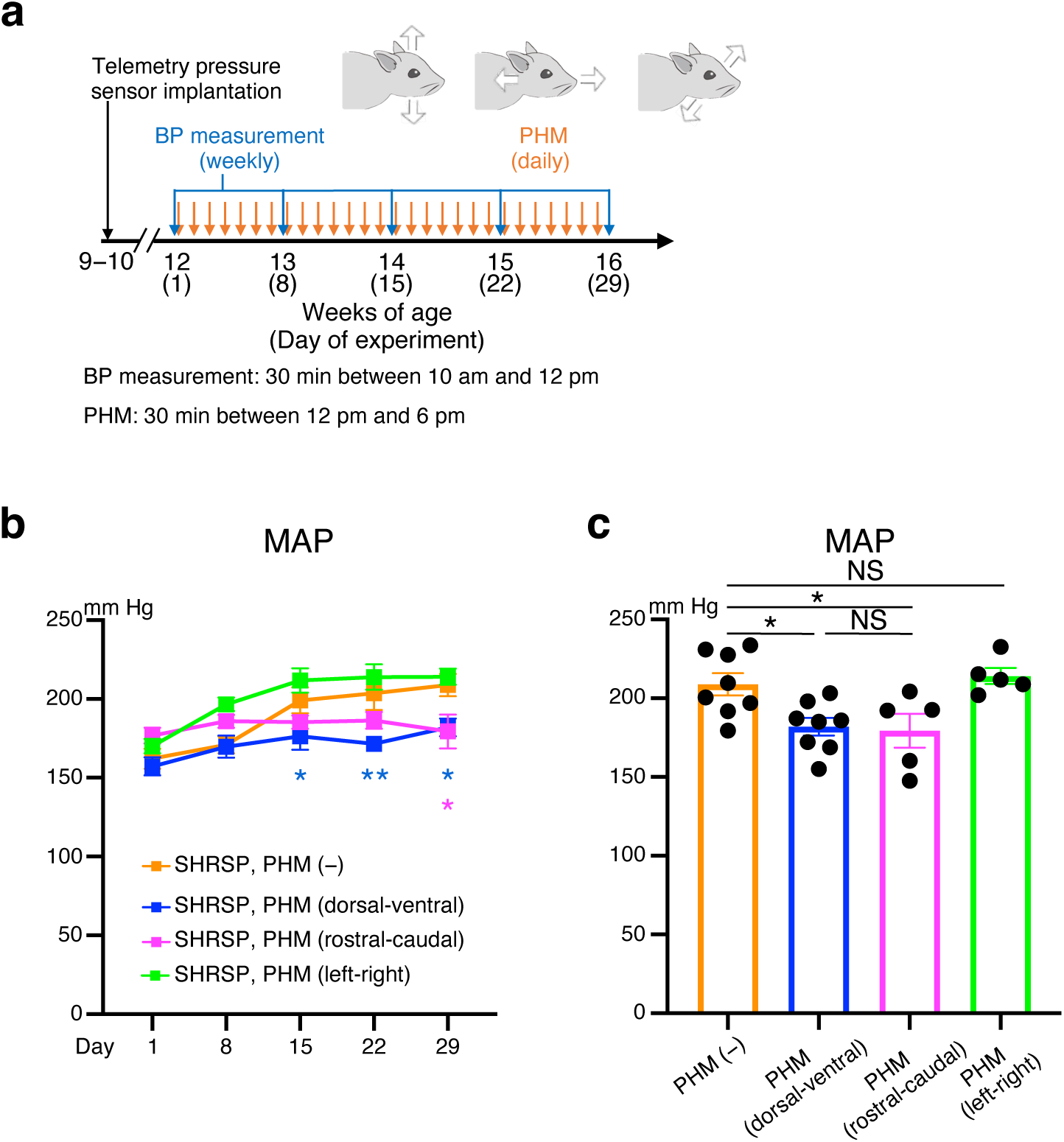
PHM in the rostral-caudal, but not left-right, direction lowers the BP in SHRSPs. **a,** Schematic representation of the experimental protocol to analyze the effects of PHM on the BP in rats. **b,c,** Time courses **(b)** and values on Day 29 **(c)** of MAP in SHRSPs, subjected to either daily PHM or anesthesia only [**b**, orange vs. blue: *P* = 0.0376 for Day 15, *P* = 0.0018 for Day 22, *P* = 0.0105 for Day 29; orange vs. magenta: *P* = 0.4131 for Day 15, *P* = 0.2448 for Day 22, *P* = 0.0148 for Day 29; orange vs. green: *P* = 0.4869 for Day 15, *P* = 0.6439 for Day 22, *P* = 0.9273 for Day 29. **c**: *P* = 0.0363 for column 1 vs. 2, *P* = 0.0465 for column 1 vs. 3, *P* = 0.9576 for column 1 vs. 4, *P* = 0.9948 for column 2 vs. 3. *n* = 8 rats for PHM (−) and PHM (dorsal-ventral); n = 5 rats for PHM (rostral-caudal) and PHM (left-right)]. The data for columns 1 and 2 in **(c)** are respectively identical to those for columns 3 and 4 in Fig. 1c. Data are presented as mean ± s.e.m. \**P* < 0.05; \*\**P* < 0.01; NS, not significant; two-way repeated measures ANOVA with Dunnett’s post hoc multiple comparisons test **(b)** or one-way ANOVA with Tukey’s post hoc multiple comparisons test **(c)**.

**Extended Data Fig. 3.**
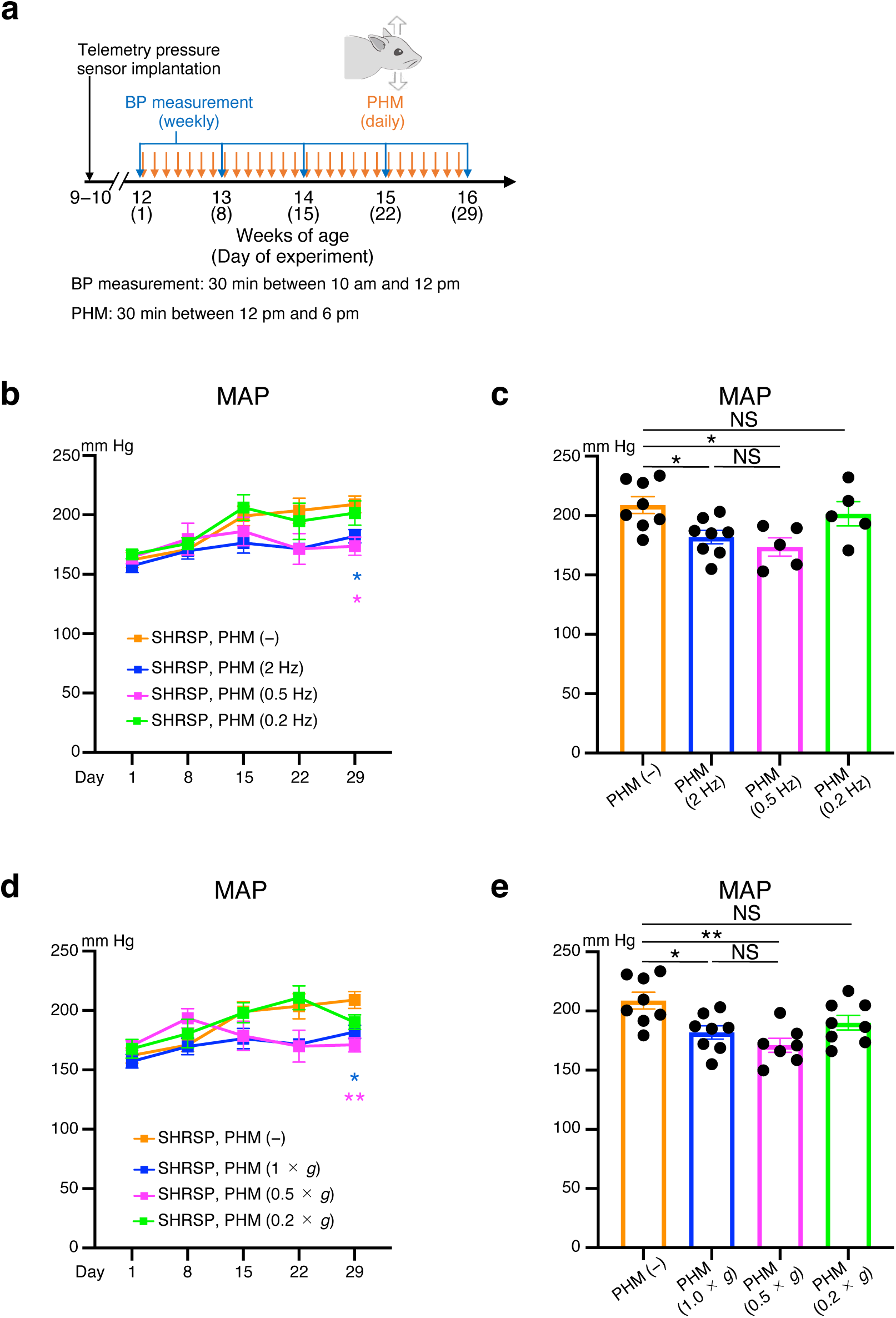
Antihypertensive effects of PHM with different frequencies or magnitudes. **a,** Schematic representation of the experimental protocol to analyze the effects of PHM on the BP in SHRSPs. **b**−**e,** Time courses **(b,d)** and values on Day 29 **(c,e)** of MAP of SHRSPs, subjected to daily PHM with either different frequencies **(b,c)** or peak accelerations **(d,e)** [**b**, orange vs. blue: *P* = 0.1871 for Day 15, *P* = 0.0519 for Day 22, *P* = 0.0282 for Day 29; orange vs. magenta: *P* = 0.7252 for Day 15, *P* = 0.2053 for Day 22, *P* = 0.0208 for Day 29; orange vs. green: *P* = 0.9212 for Day 15, *P* = 0.9356 for Day 22, *P* = 0.8950 for Day 29. **c**: *P* = 0.0435 for column 1 vs. 2, *P* = 0.0174 for column1 vs. 3, *P* = 0.9068 for column 1 vs. 4, *P* = 0.8682 for column 2 vs. 3. *n* = 8 rats for PHM (−) and PHM (2 Hz); n = 5 rats for PHM (0.5 Hz) and PHM (0.2 Hz). **d**, orange vs. blue: *P* = 0.1812 for Day 15, *P* = 0.0504 for Day 22, *P* = 0.0274 for Day 29; orange vs. magenta: *P* = 0.4146 for Day 15, *P* = 0.1740 for Day 22, *P* = 0.0036 for Day 29; orange vs. green: *P* = 0.9996 for Day 15, *P* = 0.9267 for Day 22, *P* = 0.1618 for Day 29. **e**: *P* = 0.0227 for column 1 vs. 2, *P* = 0.0014 for column 1 vs. 3, *P* = 0.1664 for column 1 vs. 4, *P* = 0.6283 for column 2 vs. 3. *n* = 8 rats for PHM (−), PHM (1 × *g*), and PHM (0.2 × *g*); n = 7 rats for PHM (0.5 × *g*)]. The data for columns 1 and 2 in **(c,e)** are respectively identical to those for columns 3 and 4 in Fig. 1c. Data are presented as mean ± s.e.m. \**P* < 0.05; \*\**P* < 0.01; NS, not significant; two-way repeated measures ANOVA with Dunnett’s post hoc multiple comparisons test **(b,d)** or one-way ANOVA with Tukey’s post hoc multiple comparisons test **(c,e)**.

**Extended Data Fig. 4.**
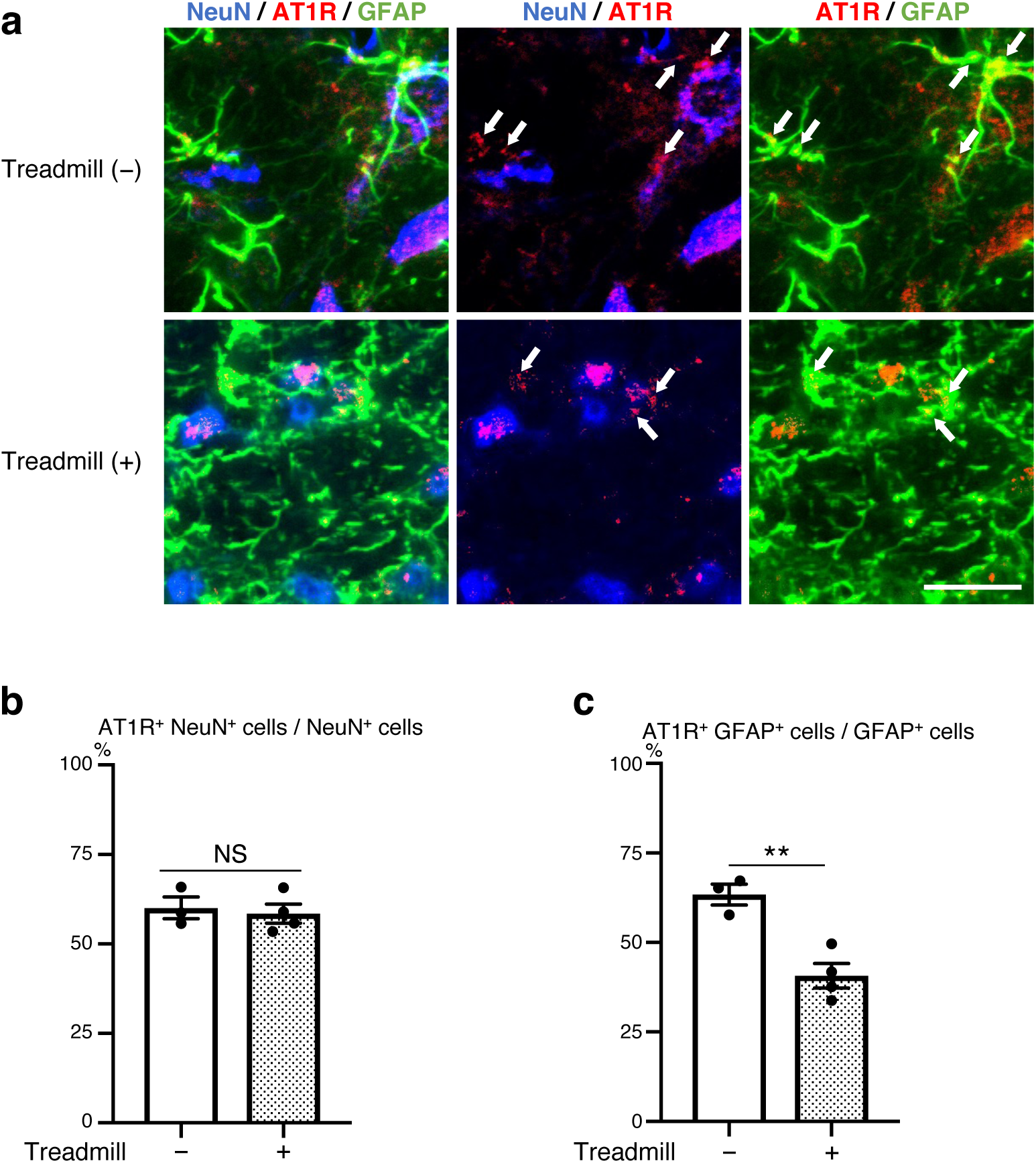
Treadmill running decreases AT1R expression in the RVLM astrocytes in SHRSPs. **a,** Micrographic images anti-NeuN (blue), anti-GFAP (green) and anti- AT1R (red) immunostaining of the RVLM of SHRSPs, either placed in the static treadmill machine or subjected to treadmill running at the velocity of 20 m/min (30 min/day, 28 days). Arrows point to anti-AT1R immunosignals that overlap with anti-GFAP, but not anti-NeuN, immunosignals in merged images. Scale bar, 50 μm. Images are representative of three or four rats. **b,c,** Quantification of AT1R-positive neurons **(b)** and astrocytes **(c)** in the RVLM of SHRSPs with or without 4-week treadmill running. Fifty NeuN^+^ cells and 100 GFAP^+^ cells were analyzed for each rat [**b**: *P* = 0.7056. **c**: *P* = 0.0048. *n* = 3 rats for treadmill (−); *n* = 4 rats for treadmill (+)]. Data are presented as mean ± s.e.m. \*\**P* < 0.01; NS, not significant, unpaired two-tailed Student’s *t-*test.

**Extended Data Fig. 5.**
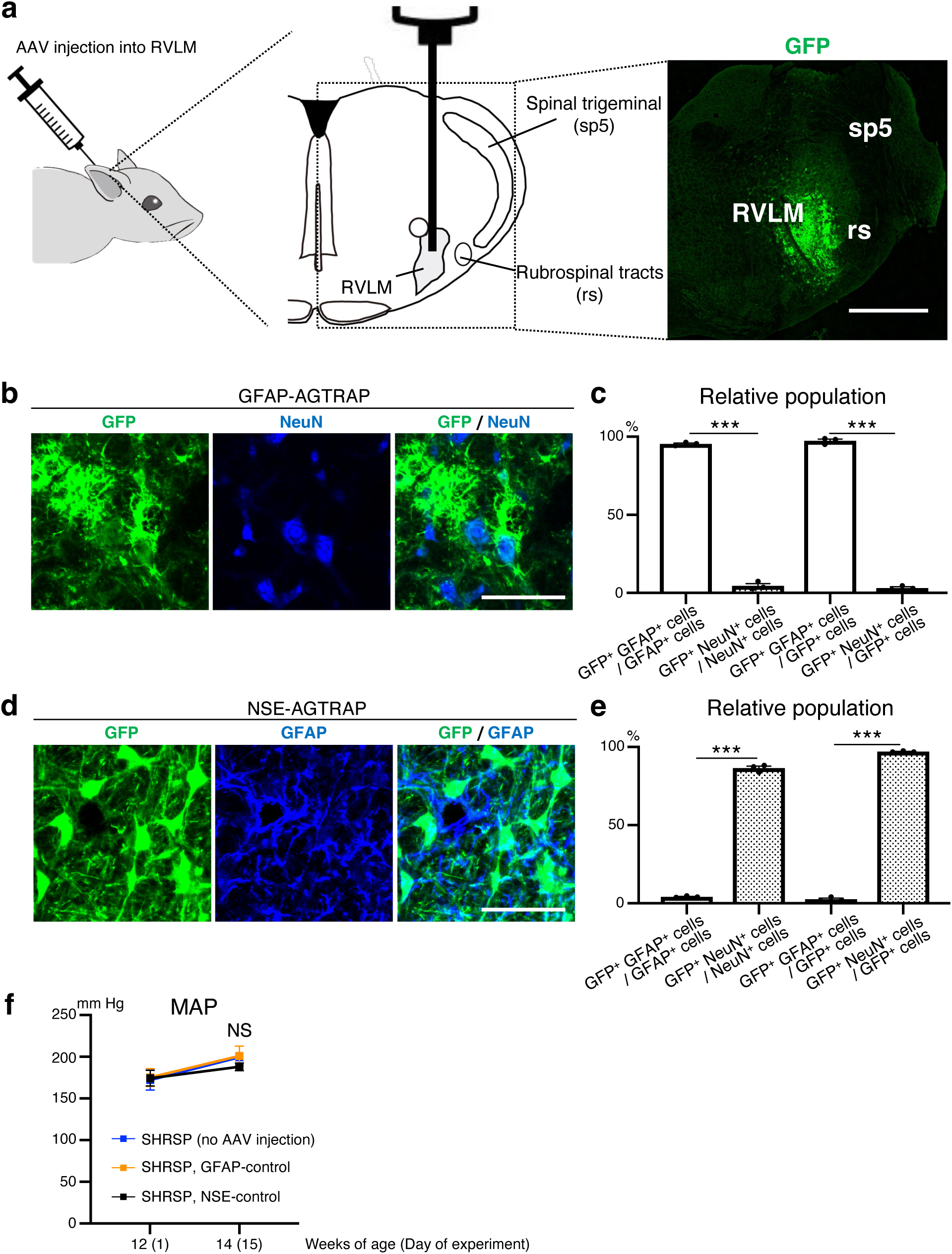
AAV-mediated transduction of RVLM astrocytes or neurons in SHRSPs. **a,** Schematic representation of the injection of AAV9 vectors to the RVLM. Micrographic image is representative of three rats analyzed in Fig. 3b (two weeks after the RVLM injection of GFAP-AGTRAP vector). GFP-derived fluorescence indicates cells expressing the transgene. Scale bar, 1 mm. **b,c,** Efficiency and specificity of astrocyte-specific expression of the transgene. **(b)** Micrographic images of GFP (green) and anti-NeuN immunostaining (blue) of the RVLM of SHRSPs analyzed in Fig. 3b. Scale bar, 50 μm. Images are representative of three rats. **(c)** Quantification of the efficiency and specificity of transgene expression. The relative populations (%) of GFP/ GFAP/ double positive (GFP^+^ GFAP^+^) or GFP/ NeuN/ double positive (GFP^+^ NeuN^+^) cells were calculated by referring their numbers to those of GFP^+^, GFAP^+^ or NeuN^+^ cells (*P* < 0.0001 for column 1 vs. 2 and *P* < 0.0001 for column 3 vs. 4. *n* = 3 rats for each group). **d,e,** Efficiency and specificity of neuron-specific expression of the transgene. **(d)** Micrographic images of GFP (green) and anti-GFAP immunostaining (blue) of the RVLM of SHRSPs analyzed in Fig. 3c. Scale bar, 50 μm. Images are representative of three rats. **(e)** Efficiency and specificity quantified as in **(c)** (*P* < 0.0001for column 1 vs. 2 and *P* < 0.0001 for column 3 vs. 4.). **f,** BP in SHRSPs injected with the control vectors. BP was measured and MAP was quantified as in Fig. 1b (*P* = 0.9918 for blue vs. orange, *P* = 0.6465 for blue vs. black, *P* = 0.5966 for orange vs. black. *n* = 6 rats for no AAV injection; *n* = 6 rats for GFAP-control; *n* = 7 rats for NSE-control). The data for blue line are identical to those demonstrated with blue line in Extended Data Fig. 1a. Data are presented as mean ± s.e.m. \*\*\**P* < 0.001; NS, not significant; unpaired two-tailed Student’s *t-*test **(c,e)** or two-way repeated measures ANOVA with Tukey’s post hoc multiple comparisons test **(f)**.

**Extended Data Fig. 6.**
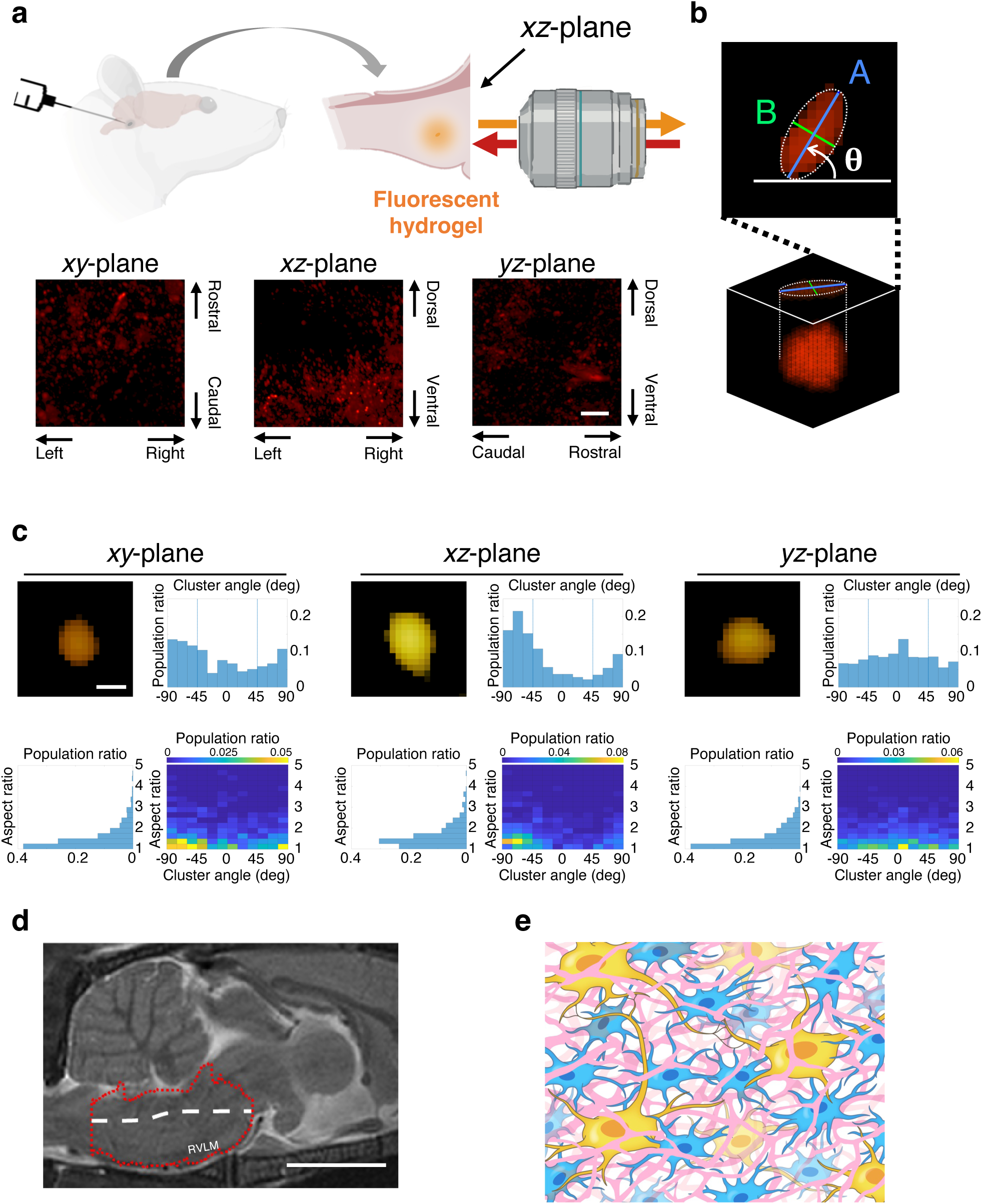
Structure and orientation of interstitial space in the rat RVLM. **a,** Multiphoton microscopic imaging of fluorescent hydrogel-introduced rat RVLM in three mutually orthogonal planes. Top: schematic illustration of hydrogel introduction and *xz*-plane imaging. Bottom: processed volume images (see Methods) of interstitial space-representing fluorescence clusters projected onto the *xy*- (bottom left), *xz*- (bottom center), and *yz*- (bottom right) planes. Scale bar, 20 μm. **b,** Determination of the cluster angle and calculation of the aspect ratio. Interstitial space-representing fluorescence clusters were extracted, and their projection and elliptic fitting on the stacked plane were conducted as described in the Methods. The cluster angle (θ) in degrees was defined and analyzed as positive (counterclockwise; see arc arrow) or negative (clockwise) from the horizontal line in each plane. The aspect ratio was calculated as A/B. **c,** Distribution of the cluster angle and aspect ratio. Volume image of a cluster of typical shape and orientation (top left), histograms of the angle distribution (top right) and aspect ratio (bottom left), and 2-D histogram of the cluster orientation/aspect ratio (bottom right) are shown for the *xy*-, *xz*-, and *yz*-planes. From the location of seemingly unimodal peaks in the 2-D histograms for the *xy*- and *xz*-planes, the *y*- and *z*-axes appear to be dominant over the *x*-axis concerning the cluster orientation. Furthermore, the *y*-axis appears slightly dominant over the *z*- axis (loose peak on the 0–15° block in the *yz*-plane). Collectively, the structure of the interstitial space-representing clusters seems loosely oriented in the direction close to the *y*-(rostral-caudal) axis with a slight counterclockwise (rostrally upward) inclination. Scale bar, 1 μm. **d,** Representative sagittal MR image of the rat brainstem (0.06 mm off the median plane). Red dotted line-indicated area and white broken line represent the lower brainstem and longest centroidal line of this part of the brain, respectively (see Methods). Note that the centroidal line is approximately along the *y*-axis with a slight rostrally upward inclination. Scale bar, 5 mm. Data shown in **(c,d)** represent at least three independent samples (rats) with similar results. **e,** Diagram of the mesoscale structure/orientation of interstitial space in the rat RVLM. The interstitial space (depicted in pink) is not randomly structured but oriented approximately along the direction of the centroidal line of this part of the brain with relatively minor lateral communications.

**Extended Data Fig. 7.**
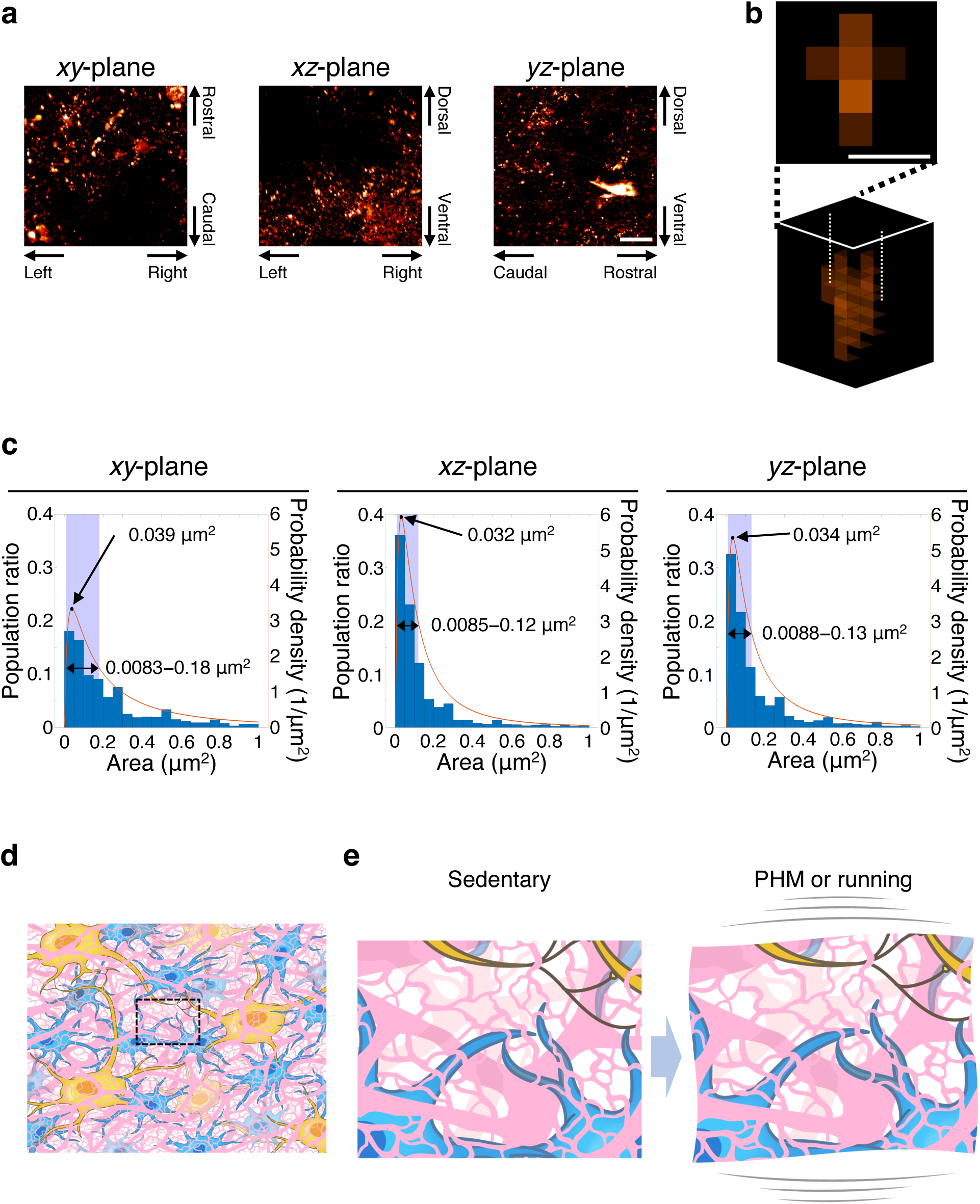
Dimension (cross-sectional area) of interstitial space in the rat RVLM. **a,** Multiphoton microscopic images of interstitial space-representing fluorescence clusters projected onto *xy*- (left), *xz*- (center), and *yz*- (right) planes. Scale bar, 20 μm. **b,** Projection of 4-μm stack images of an individual cluster. Scale bar, 0.5 μm. **c,** Distribution of the cross-sectional areas of individual clusters. Red curves represent the probability densities revealed by fitting to log-normal distribution^77, 78^. Black circles and two-way arrows indicate the mode and full width at half maximum (FWHM)^79^, respectively. **d,** Diagram of the interstitial space in the rat RVLM. Dotted rectangle indicates the area illustrated in **(e)**. Because we analyzed the interstitial space with imaging experiments based on the spread of the injected gelatable fluorescent PEG solutions, we assumed the interstitial space to have a continuous structure. Nevertheless, we do not preclude the possible existence of dead ends of interstitial space. **e,** Illustration of cyclic microdeformation during PHM or treadmill running that generates small pressure changes in the rat RVLM, facilitating or promoting interstitial fluid movement in situ.

**Extended Data Fig. 8.**
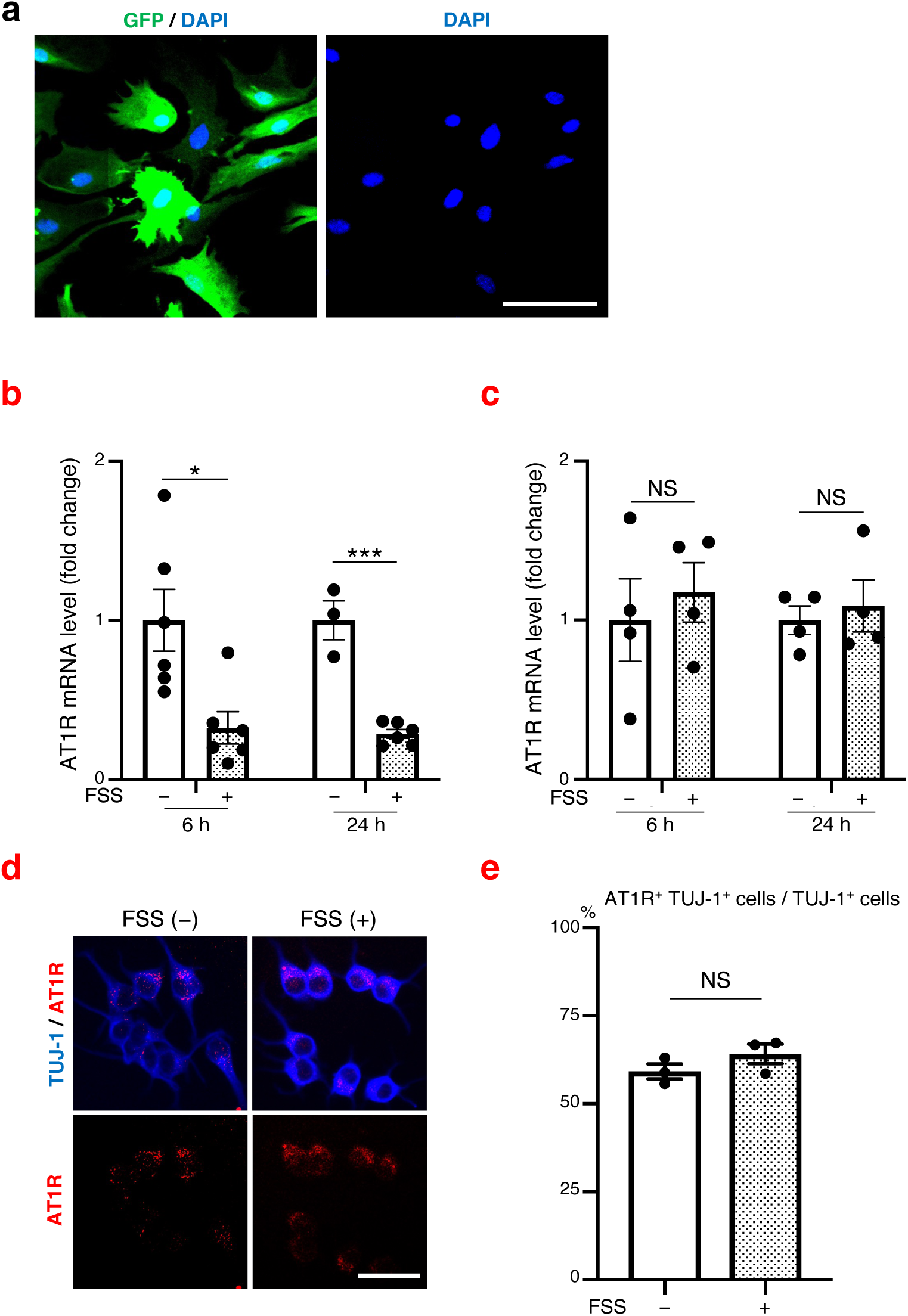
Preparation of astrocyte-enriched primary culture, and lack of decreasing effect of FSS on AT1R expression in cultured neuronal cells. **a,** Representative micrographic image of anti-GFP immunostaining (green) and DAPI staining (blue) of astrocyte-enriched culture prepared from the astrocyte-GFP mice. Scale bar, 50 μm. Note that most of the cells are GFP-positive. **b**,**c,** Effects of FSS on the AT1R mRNA expression in astrocytes and Neuro2A cells. AT1R mRNA expression in cultured astrocytes **(b)** and Neuro2A cells **(c)** 6 or 24 h after 30-min FSS application (0.7 Pa, 0.5 Hz) was analyzed as in Fig. 5a [**b**: *P* = 0.0116 for 6 h, *P* < 0.0001 for 24 h. *n* = 6 for 6 h; *n* = 3 for 24 h/FSS (−); *n* = 6 for 24 h/FSS (+). **c**: *P* = 0.6065 for 6 h, *P* = 0.6490 for 24 h. *n* = 4 for each group]. **d,** Microscopic images of anti-AT1R (red) and anti-TUJ-1 (blue) immunostaining of Neuro2A cells, either left unexposed or exposed to pulsatile FSS (0.7 Pa, 0.5 Hz, 30 min), and fixed 24 h after the intervention. Images are representative of three independent experiments with similar results. Scale bar, 50 μm. **e,** Relative population of AT1R/TUJ-1/ double positive (AT1R^+^ TUJ-1^+^) cells quantified as a ratio to total TUJ-1-positive (TUJ-1^+^) cells in each sample (*P* = 0.2308. More than 100 TUJ-1- positive cells were analyzed in each sample. *n* = 3). Data are presented as mean ± s.e.m. \**P* < 0.05; \*\*\**P* < 0.001; NS, not significant; unpaired two-tailed Student’s *t-*test **(b,c,e)**.

**Extended Data Fig. 9.**
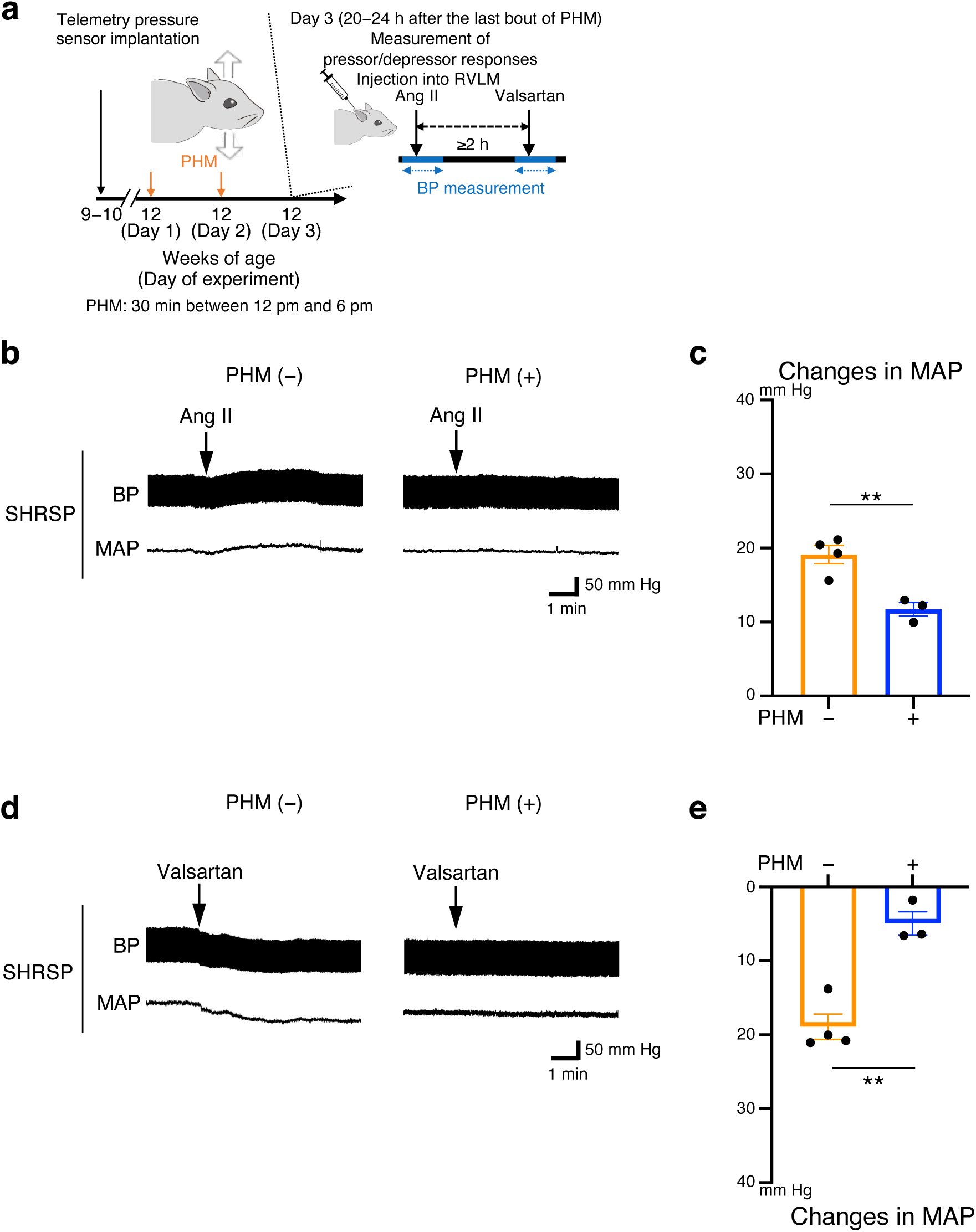
Two-day PHM alleviates the sensitivity of RVLM in SHRSPs to Ang II or valsartan. **a,** Schematic representation of the experimental protocol. Pressor and depressor responses were analyzed as in Fig. 2. **b**−**e,** Pressor **(b,c)** and depressor **(d,e)** responses in SHRSPs with and without 2-day PHM. **(b,d)** Representative trajectories of the BP (top in each panel) and MAP (bottom in each panel). Arrows point to the time of the initiation of RVLM injection of Ang II **(b)** or valsartan **(d)**. Right-angled scale bars, 1 min / 50 mm Hg. **(c,e)** MAP change caused by Ang II **(c)** or valsartan **(e)** injection [**c**: *P* = 0.0064. **e**: *P* = 0.0022. *n* = 4 rats for PHM (–); *n* = 3 rats for PHM (+)]. Data are presented as mean ± s.e.m. \*\**P* < 0.01; unpaired two-tailed Student’s *t-*test.

**Extended Data Fig. 10.**
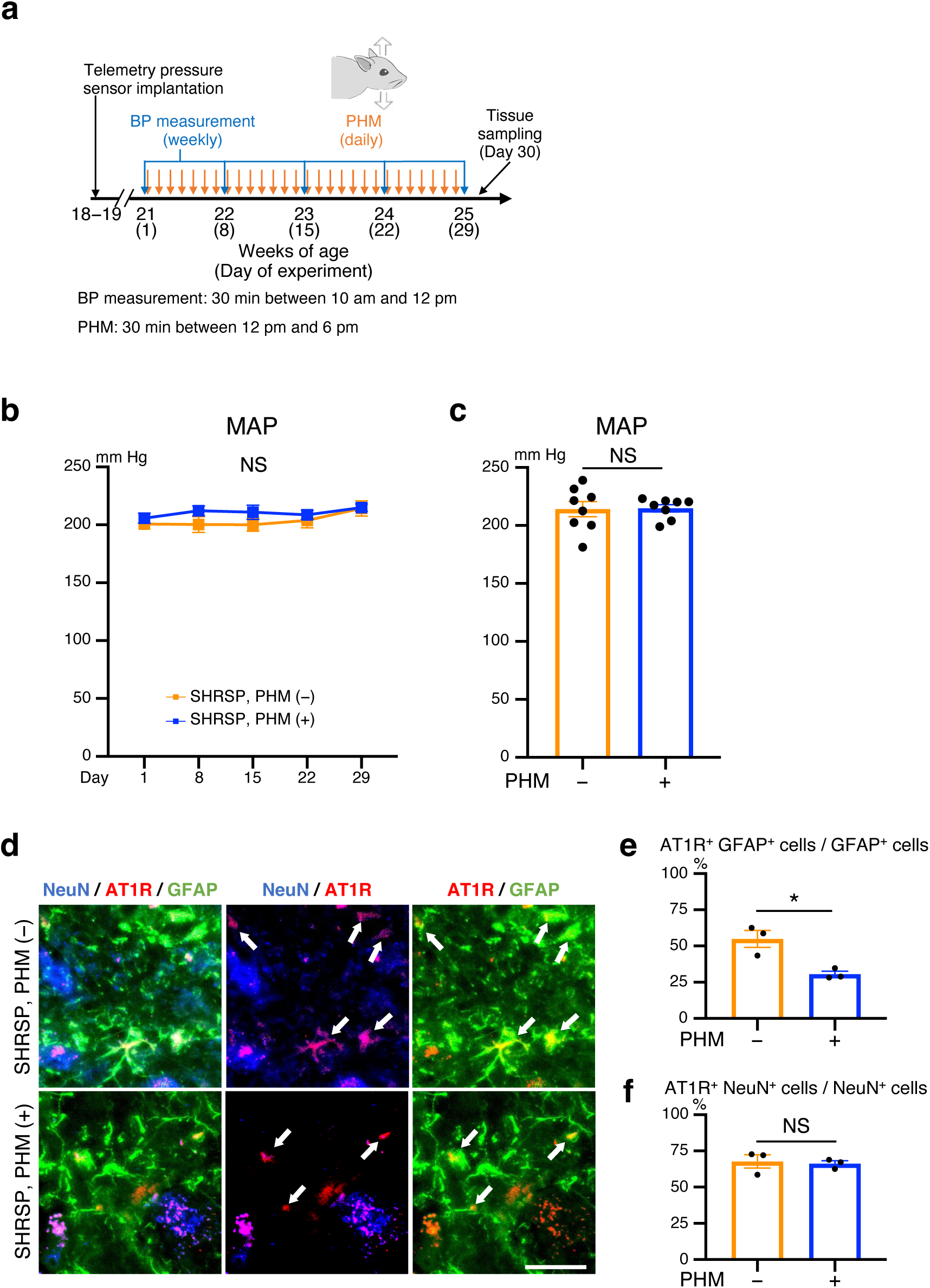
PHM does not lower the BP in SHRSPs, but decreases the AT1R expression in their RVLM astrocytes during the plateau phase of hypertension development. **a,** Schematic representation of the experimental protocol to analyze the effects of PHM. **b,c,** Time courses **(b)** and values on Day 29 **(c)** of the MAP of SHRSPs, subjected to either daily PHM or anesthesia only (**b**: *P* > 0.9999 for Day 15, Day 22, and Day 29. **c**: *P* = 0.9167. *n* = 8 rats for each group). **d,** Micrographic images of anti-GFAP (green), anti-AT1R (red), and anti-NeuN (blue) immunostaining of the RVLM of SHRSPs, either left sedentary (top) or subjected to PHM (bottom) under anesthesia (30 min/day, 28 days). Arrows point to anti-AT1R immunosignals that overlap with anti-GFAP, but not anti-NeuN, immunosignals in merged images. Scale bars, 50 µm. Images are representative of three rats. **e,f,** Quantification of AT1R-positive astrocytes **(e)** and neurons **(f)** in the RVLM of SHRSPs, either left sedentary or subjected to PHM. One-hundred GFAP-positive (GFAP^+^) cells **(e)** and 50 NeuN-positive (NeuN^+^) cells **(f)** were analyzed for each rat (**e**: *P* = 0.0173. **f**: *P* = 0.7812. *n* = 3 rats for each group). Data are presented as mean ± s.e.m. \**P* < 0.05; NS, not significant; two-way repeated measures ANOVA with Bonferroni’s post hoc multiple comparisons test **(b)** or unpaired two- tailed Student’s *t-*test **(c,e,f)**.

**Extended Data Fig. 11.**
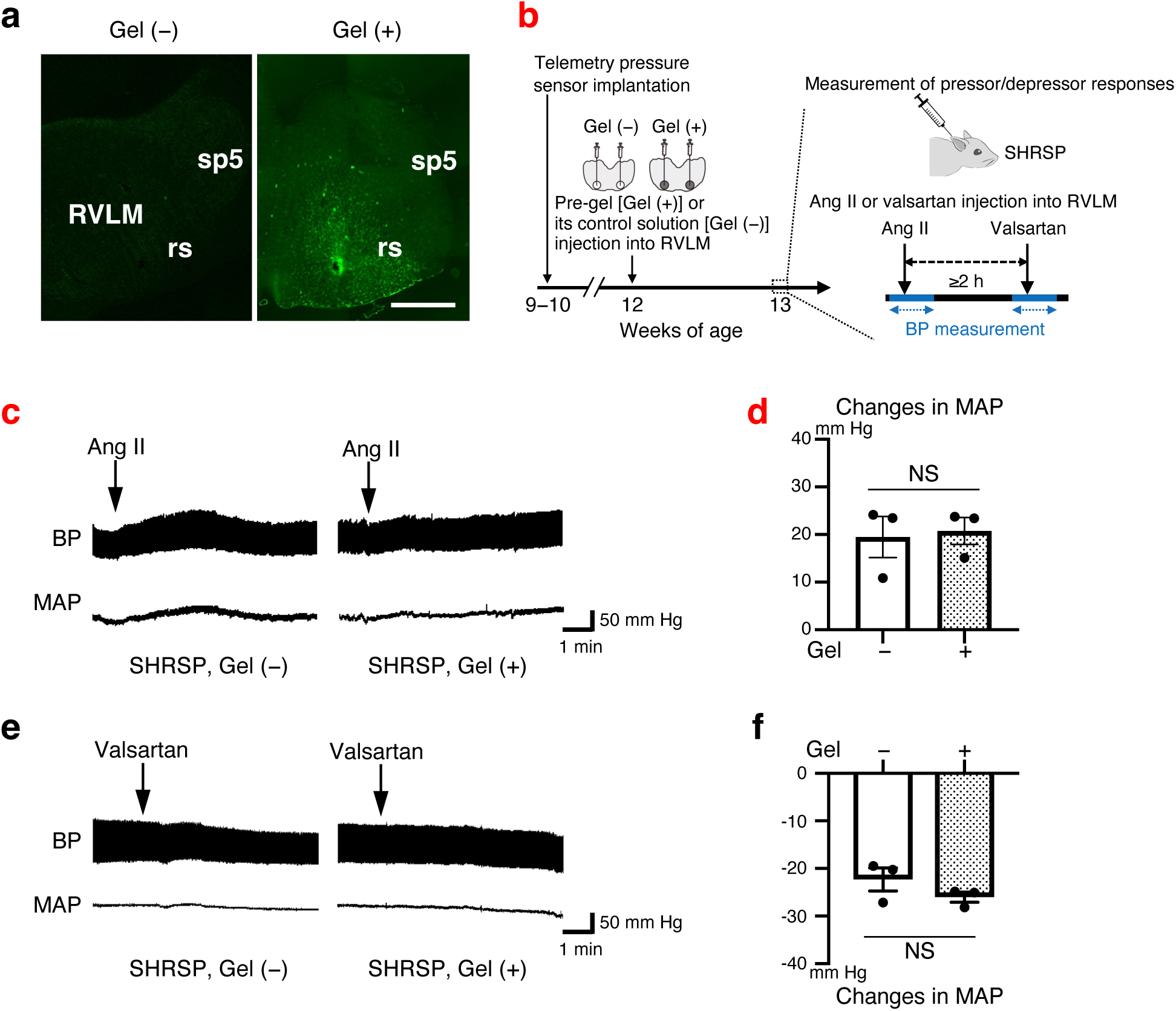
Hydrogel introduction in the RVLM does not affect the pressor and depressor responses to AngII and valsartan, respectively, in SHRSPs. **a,** Introduction of the PEG hydrogel in the rat RVLM. Twenty-four hours after the injection of control ungelatable fluorescent PEG solution (left) or one week after the injection of pre-gel fluorescent PEG solution (right), brainstem samples were prepared. Coronal-section images representative of three rats with similar results are shown. Scale bar, 1 mm. **b–f,** Pressor and depressor responses analyzed one week after the injection of pre-gel PEG solution or its ungelatable control. **(b)** Schematic representation of the experimental protocol. Pressor and depressor responses were analyzed as in Fig. 2. **(c–f)** Representative trajectories **(c,e)** and quantification **(d,f)** of the BP ascent upon AngII injection **(c,d)** and descent upon valsartan injection **(e,f)** to the RVLM of SHRSPs with or without the hydrogel introduction (**d**: *P* = 0.8226. **f**: *P* = 0.2342. *n* = 3 rats for each group). Right-angled scale bars, 1 min / 50 mm Hg. Data are presented as mean ± s.e.m. NS, not significant; unpaired two-tailed Student’s *t-*test.

**Extended Data Fig. 12.**
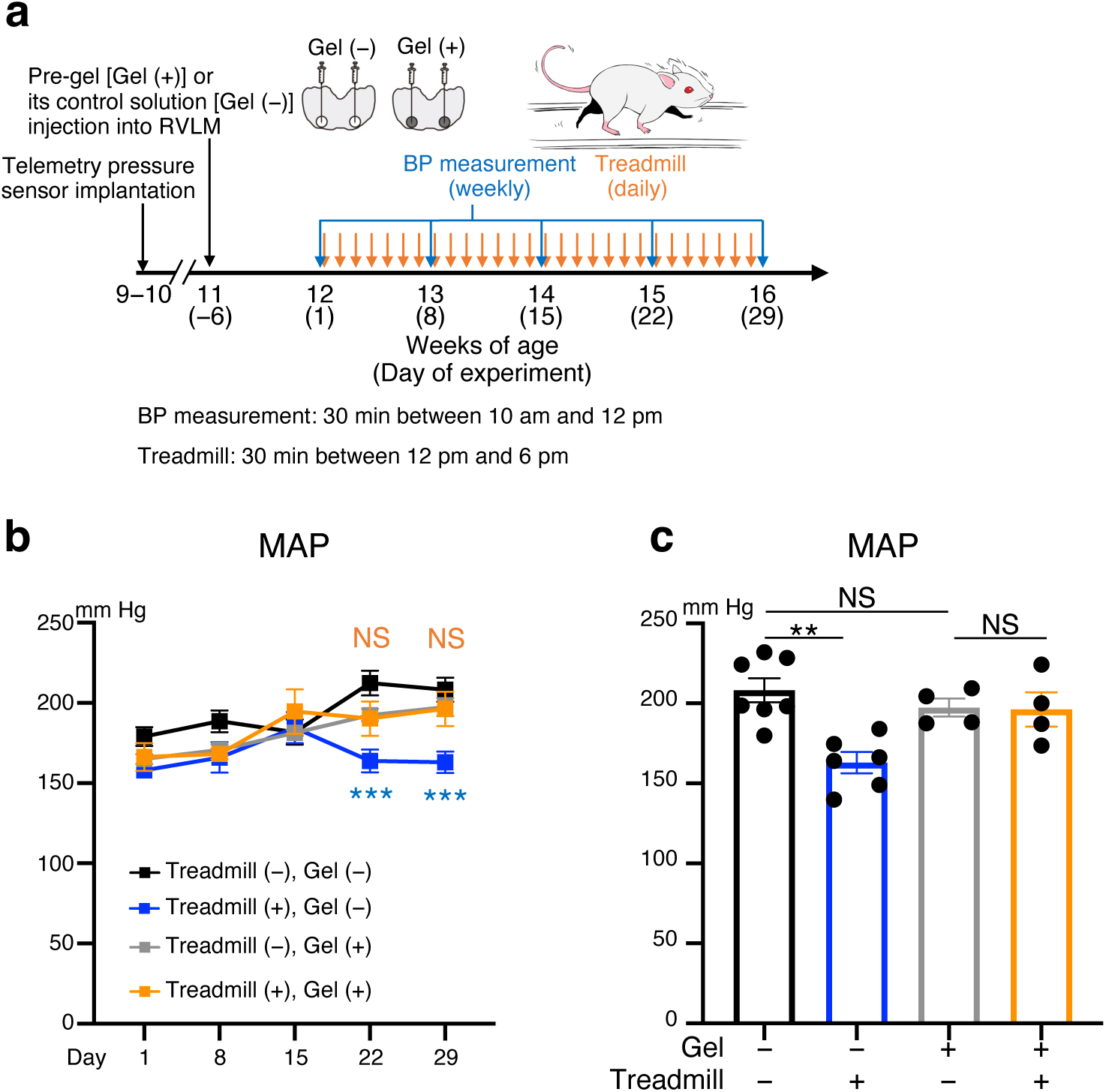
Hydrogel introduction to the RVLM eliminates the antihypertensive effect of treadmill running in SHRSPs. **a,** Schematic representation of the experimental protocol to analyze the effects of treadmill running on the BP in SHRSPs. **b,c,** Time courses **(b)** and values on Day 29 **(c)** of the MAP of SHRSPs with (**b**: gray and orange lines, **c**: columns 3 and 4) and without (**b**: black and blue lines, **c**: columns 1 and 2) the hydrogel introduction, subjected to either daily treadmill running (30 min/day, 28 days; **b**: blue and orange lines, **c**: columns 2 and 4) or its control (placing on the belt without turning on the treadmilling; **b**: black and gray lines, **c**: columns 1 and 3) [**b**, black vs. blue: *P* > 0.9999 for Day 15, *P* = 0.0001 for Day 22, *P* = 0.0004 for Day 29; gray vs. orange: *P* > 0.9999 for Day 15, Day 22, and Day 29. **c**: *P* = 0.0015 for column 1 vs. 2, *P* = 0.9997 for column 3 vs. 4, *P* = 0.7751 for column 1 vs. 3. *n* = 7 rats for treadmill (–)/Gel (–); *n* = 6 rats for treadmill (+)/Gel (–); *n* = 4 rats for each group of Gel (+)]. Data are presented as mean ± s.e.m. \*\**P* < 0.01, \*\*\**P* < 0.001; NS, not significant; two-way repeated measures ANOVA with Bonferroni’s post hoc comparisons test **(b)** or one-way ANOVA with Tukey’s post hoc multiple comparisons test **(c)**.

**Extended Data Fig. 13.**
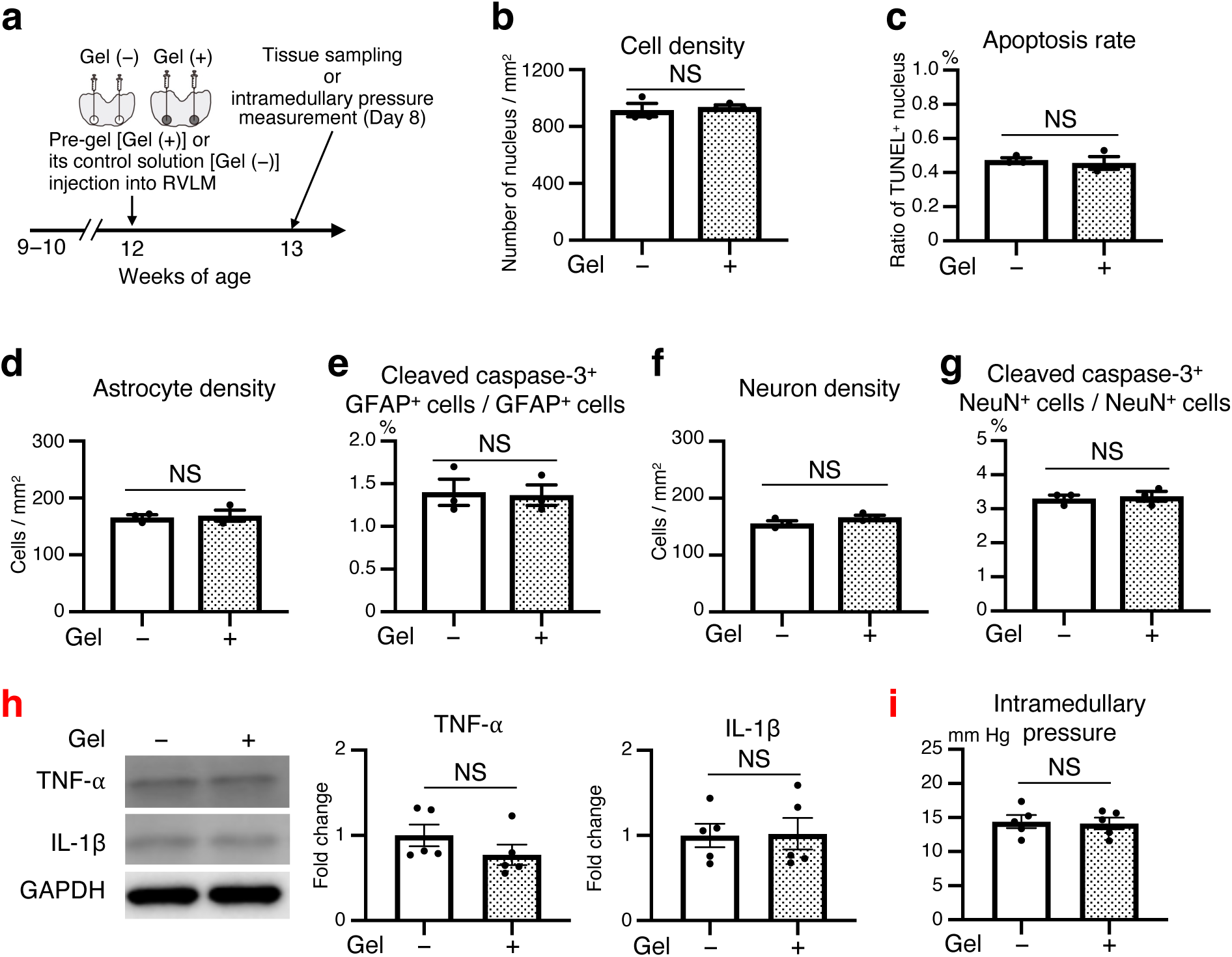
Hydrogel introduction does not affect the cell number/apoptosis, the expression of pro-inflammatory cytokines, and the pressure in the RVLM of SHRSPs. **a,** Schematic representation of the experimental protocol to analyze the hydrogel-introduced RVLM in SHRSPs. **b−g,** Effects of hydrogel introduction on the survival/apoptosis of RVLM cells in SHRSPs. Fixed rat RVLM sections were subjected to TUNEL assay **(b,c)**, or combinations of anti-GFAP, anti-NeuN and anti-cleaved caspase-3 immunostaining **(e−g)**. DAPI-positive nuclei **(b)**, GFAP- **(d)** or NeuN- **(f)** positive cells were counted, and the relative populations of cells doubly positive for indicated combinations of TUNEL **(c)** and cleaved caspase-3 **(e,g)** were quantified. Each value in **(b−g)** represents an average from five images of 1 x 1-mm area analyzed for each rat. **h,** Expression of TNF-α and IL-1β in SHRSPs’ RVLM with and without the hydrogel introduction. Anti-TNF-α, anti-IL-1β, and anti-GAPDH immunoblots (left). Expressions of TNF-α and IL-1β were normalized against GAPDH expression, and scaled with the mean values of the control samples [Gel (−)] set as 1. **i,** Intramedullary pressure of SHRSPs with and without the hydrogel introduction in their RVLM. The intramedullary pressure was measured as in Fig. 4a−d, and the mean value was obtained from 30-s steady-state continuous measurement for each rat (**b**: *P* = 0.6518. **c**: *P* = 0.6943. **d**: *P* = 0.7938. **e**: *P* = 0.8722. **f**: *P* = 0.1679. **g**: *P* = 0.7247. **h**: *P* = 0.2312 for TNF-α, *P* = 0.9342 for IL-1β. **i**: *P* = 0.8433. **b-g**: *n* = 3 rats for each group. **h,i**: *n* = 5 rats for each group). Data are presented as mean ± s.e.m. NS, not significant; unpaired two-tailed Student’s *t-*test **(b-i)**.

**Extended Data Fig. 14.**
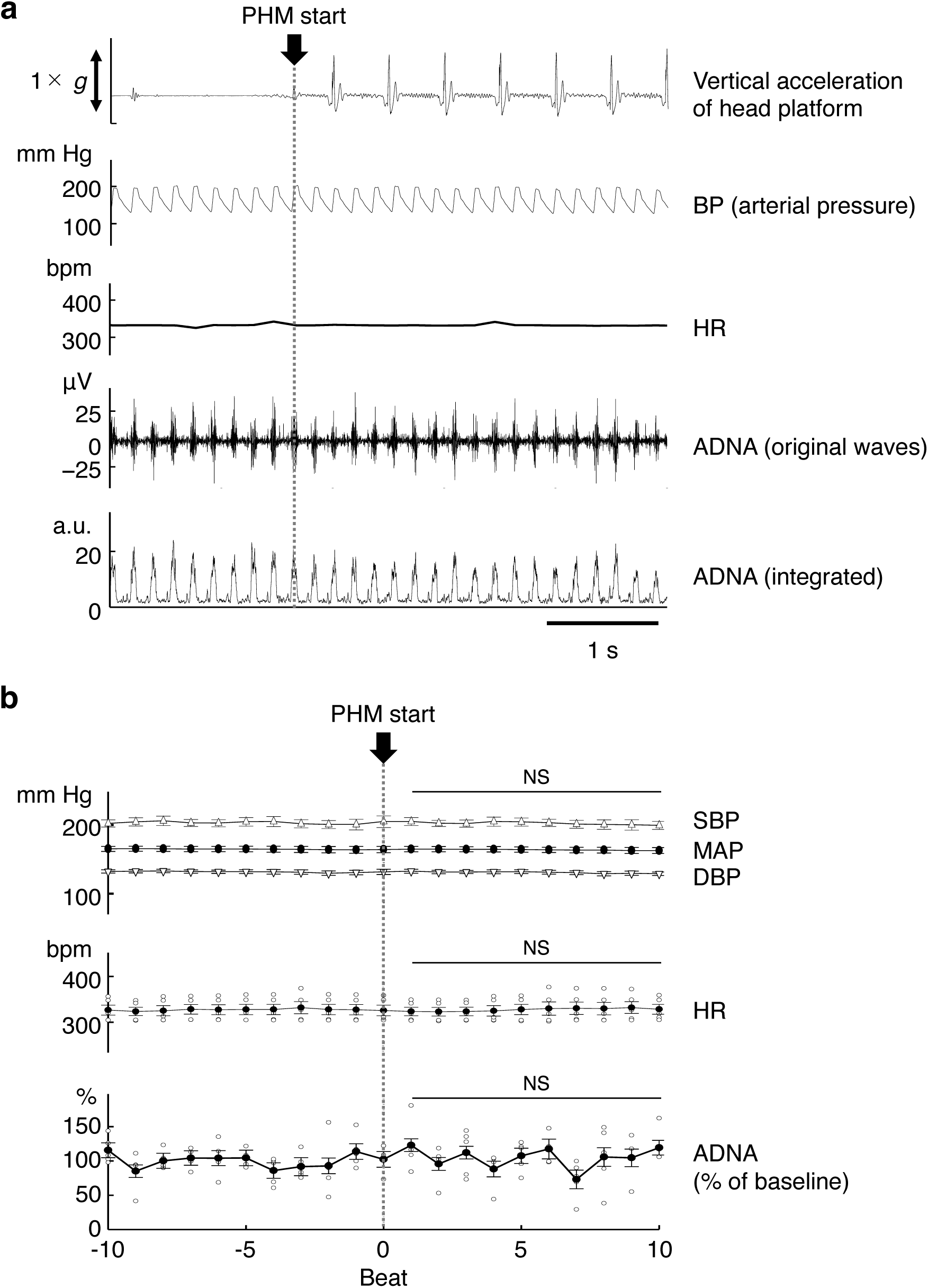
BP, HR, and aortic depressor nerve activity (ADNA) remain unchanged during the transition from before to after the initiation of PHM in SHRSPs. **a,** BP, HR, and ADNA in a 16-week-old male SHRSP monitored and recorded simultaneously with vertical accelerations of the oscillating PHM platform. Integrated ADNA was normalized in each rat and is presented in arbitrary units (a.u.) (see Methods). Data shown represent five independent experiments using five different SHRSPs with similar results. **b,** Beat-by-beat BP (SBP, MAP, and DBP), HR, and ADNA. The beat that elicited SBP immediately before the PHM initiation was defined as “beat 0”. Statistical analysis was conducted against the mean values of beat –9 to beat 0 (SBP; *P* = 0.5829 for beat 1, *P* = 0.9994 for beat 2, *P* = 0.9627 for beat 3, *P* = 0.4169 for beat 4, *P* = 0.6948 for beat 5, *P* = 0.9999 for beat 6, *P* = 0.8152 for beat 7, *P* = 0.1381 for beat 8, *P* = 0.3191 for beat 9, and *P* = 0.5286 for beat 10. MAP; *P* = 0.9906 for beat 1, *P* > 0.9999 for beat 2, *P* = 0.9997 for beat 3, *P* = 0.9997 for beat 4, *P* = 0.9349 for beat 5, *P* = 0.3562 for beat 6, *P* = 0.2239 for beat 7, *P* = 0.1599 for beat 8, *P* = 0.3001 for beat 9, and *P* = 0.2596 for beat 10. DBP; *P* = 0.9810 for beat 1, *P* > 0.9999 for beat 2, *P* > 0.9999 for beat 3, *P* > 0.9999 for beat 4, *P* = 0.9996 for beat 5, *P* = 0.9997 for beat 6, *P* = 0.9421 for beat 7, *P* = 0.5327 for beat 8, *P* = 0.4949 for beat 9, and *P* = 0.5215 for beat 10. HR; *P* = 0.8547 for beat 1, *P* = 0.7825 for beat 2, *P* =0.8267 for beat 3, *P* = 0.6800 for beat 4, *P* = 0.9996 for beat 5, *P* = 0.9834 for beat 6, *P* = 0.8658 for beat 7, *P* = 0.9460 for beat 8, *P* = 0.4602 for beat 9, and *P* =0.8992 for beat 10. ADNA; *P* = 0.6663 for beat 1*, P* = 0.9996 for beat 2, *P* = 0.9258 for beat 3, *P* = 0.8960 for beat 4, *P* = 0.9211 for beat 5, *P* = 0.4492 for beat 6, *P* = 0.3520 for beat 7, *P* =0.9995 for beat 8, *P* =0.9979 for beat 9, and *P* = 0.5181 for beat 10. *n* = 5 rats). Data are presented as mean ± s.e.m. NS, not significant; one-way repeated measures ANOVA with Dunnett’s multiple comparisons test. Arrows and dotted lines in (**a,b**) indicate the time point of PHM initiation.

**Extended Data Fig. 15.**
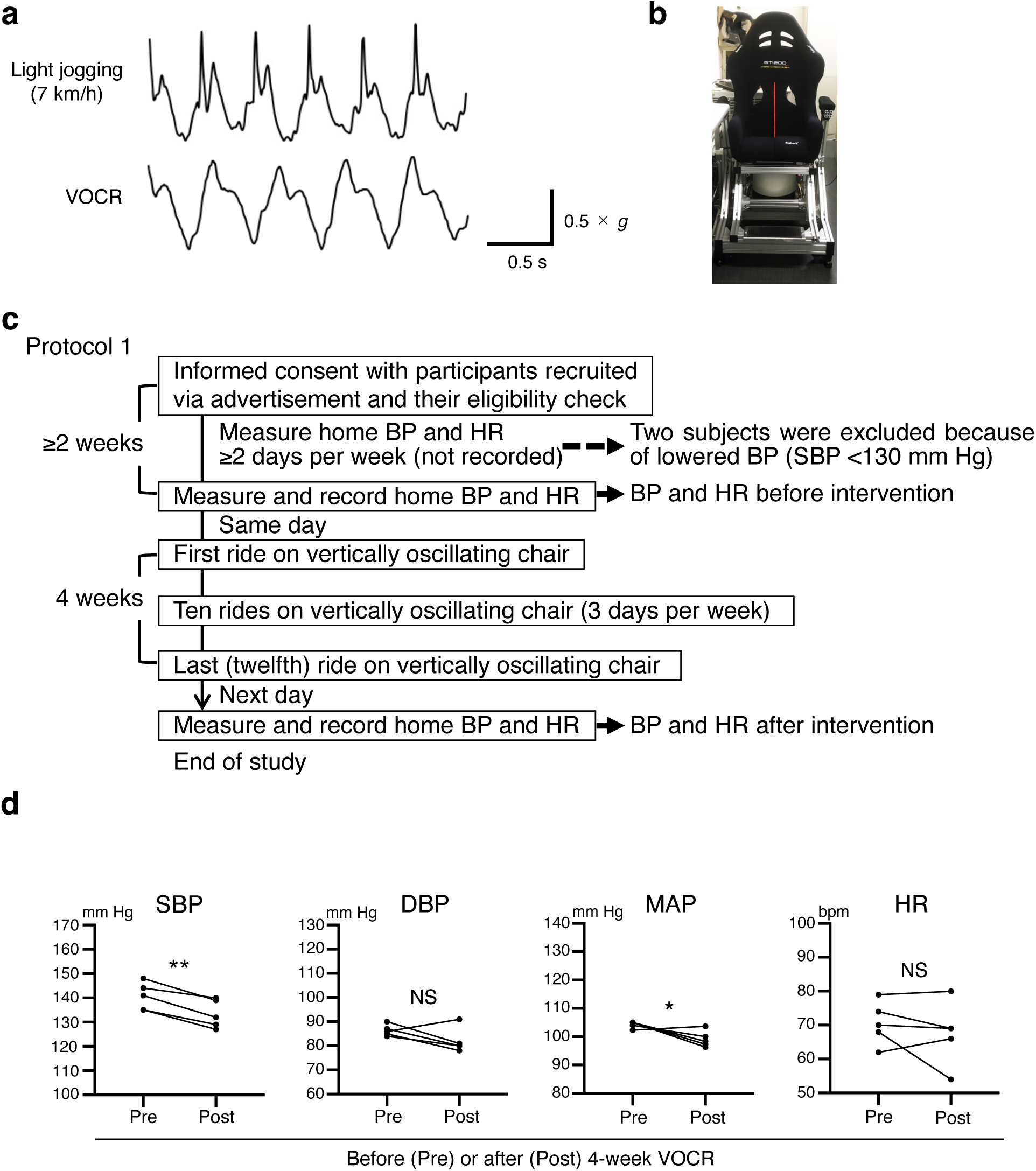
Vertically oscillating chair that reproduces the mechanical accelerations in the head during light jogging, and the design and results of protocol 1 as a pilot study to examine antihypertensive effect of VOCR in hypertensive adult humans. **a,** Vertical accelerations generated at adult human head during light jogging on a treadmill machine (velocity: 7 km/h) and VOCR (frequency: 2 Hz). The VOCR system was adjusted to produce ∼1.0 × *g* vertical acceleration peaks. Right-angled scale bar, 0.5 s / 0.5 × *g*. Images are representative of three independent experiments with similar results. **b**, Photograph of the chair. **c,** Schematic representation of protocol 1. **d,** BP and HR “value of the day”s immediately before and after 4-week VOCR in the study of protocol 1 (SBP: *P* = 0.0018. DBP: *P* = 0.1509. MAP: *P* = 0.0459. HR: *P* = 0.3900. *n* = 5). \**P* < 0.05; \*\**P* < 0.01; NS, not significant; paired two-tailed Student’s *t-*test.

**Extended Data Fig. 16.**
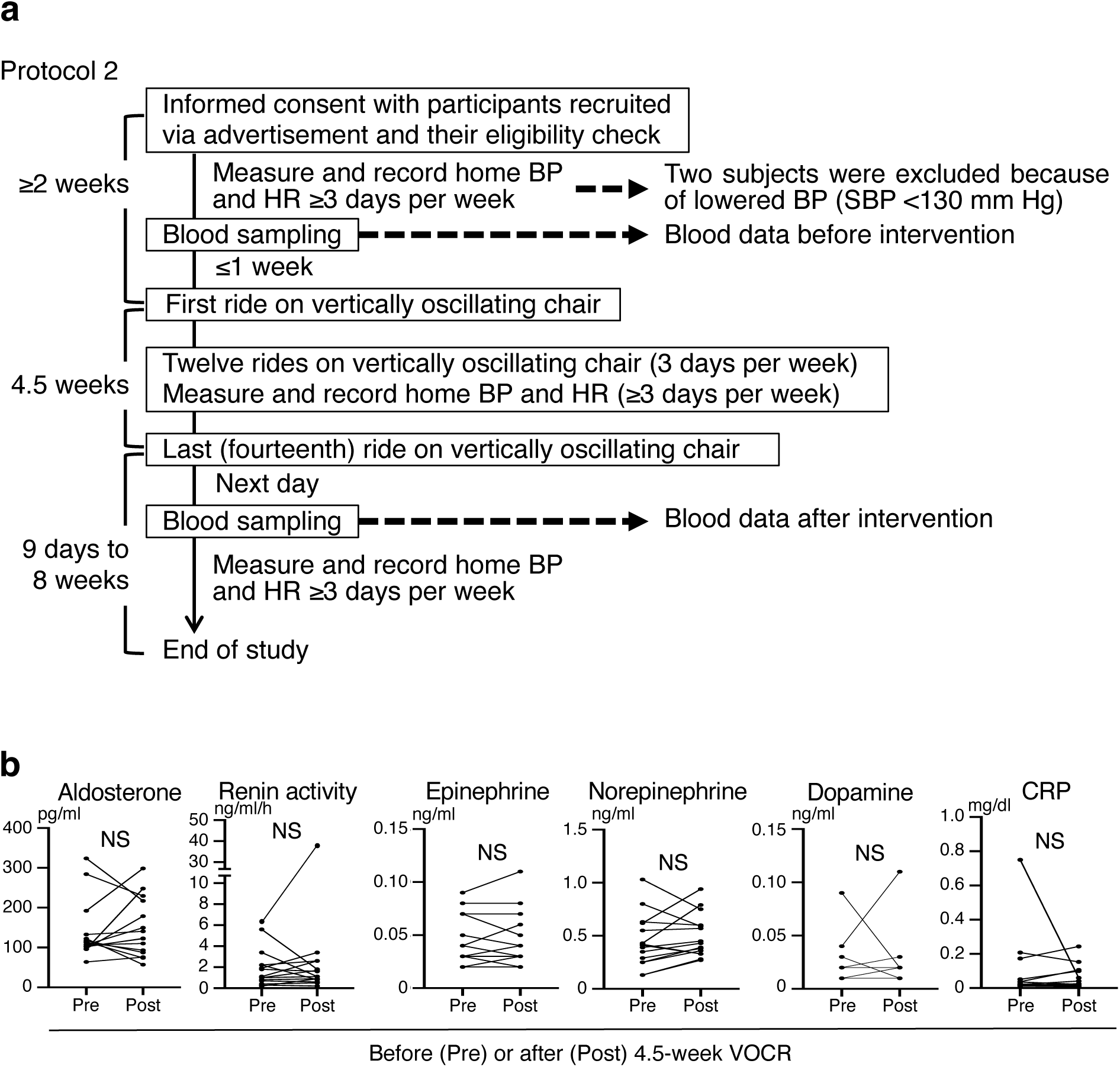
Design and blood test results of the human study of protocol 2. **a,** Schematic representation of protocol 2. **b,** Blood test values before and after the intervention period. Significant change was not observed in any of tested parameters. NS, not significant; paired two-tailed Student’s *t-*test. (Aldosterone: *P* = 0.6265. Renin activity: *P* = 0.3794. Epinephrine: *P* = 0.5103. Norepinephrine: *P* = 0.2653. Dopamine: *P* > 0.9999. CRP: *P* = 0.4412. *n* = 15. A participant (subject #18) showed a large increase in plasma renin activity after VOCR. We advised him to consult his primary care physician, who ruled out the disqualifying conditions for this study (e.g., severe renal disease; see Methods) based on comprehensive evaluation. Therefore, we did not exclude subject #18 from our statistical analysis of BP and HR.

**Extended Data Fig. 17.**
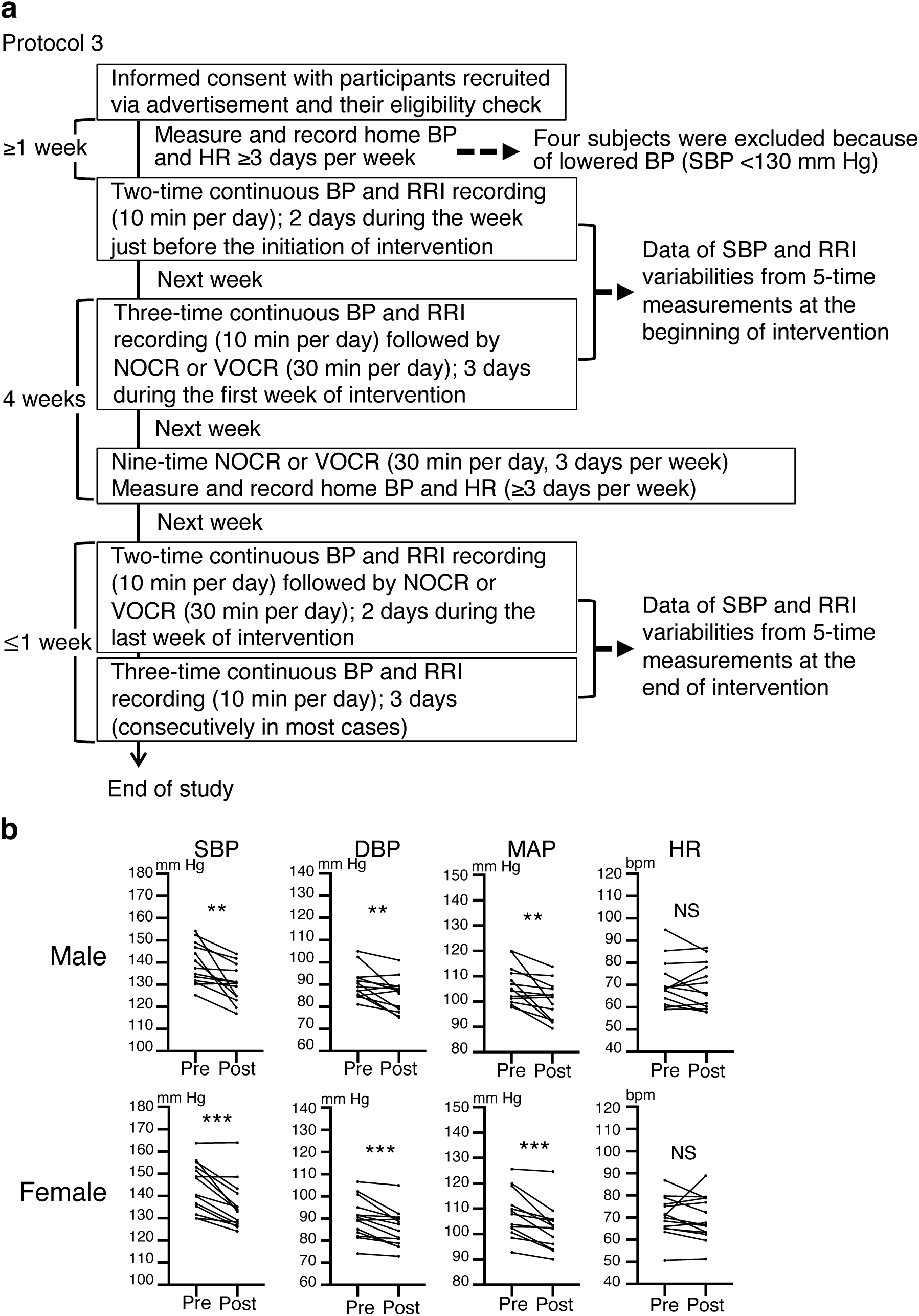
Design of the human study of protocol 3, and sex-segregated analysis of BP and HR in protocols 2 and 3. **a,** Schematic representation of protocol 3. **b,** BP and HR “value of the week”s immediately before and after 4.5-week VOCR in the study of protocols 2 and 3. To avoid duplicate inputs, the VOCR data from subjects #22 and #31 in protocol 3, who were subjects #15 and #7 in protocol 2 (see Supplementary Table 3), were excluded from this analysis (male, SBP: *P* = 0.0033. DBP: *P* = 0.0094. MAP: *P* = 0.0046. HR: *P* = 0.4373. *n* = 13; female, SBP: *P* = 0.0002. DBP: *P* = 0.0005. MAP: *P* = 0.0003. HR: *P* = 0.6257. *n* = 14). \*\**P* < 0.01; \*\*\**P* < 0.001; NS, not significant; paired two-tailed Student’s *t-*test.

**Extended Data Fig. 18.**
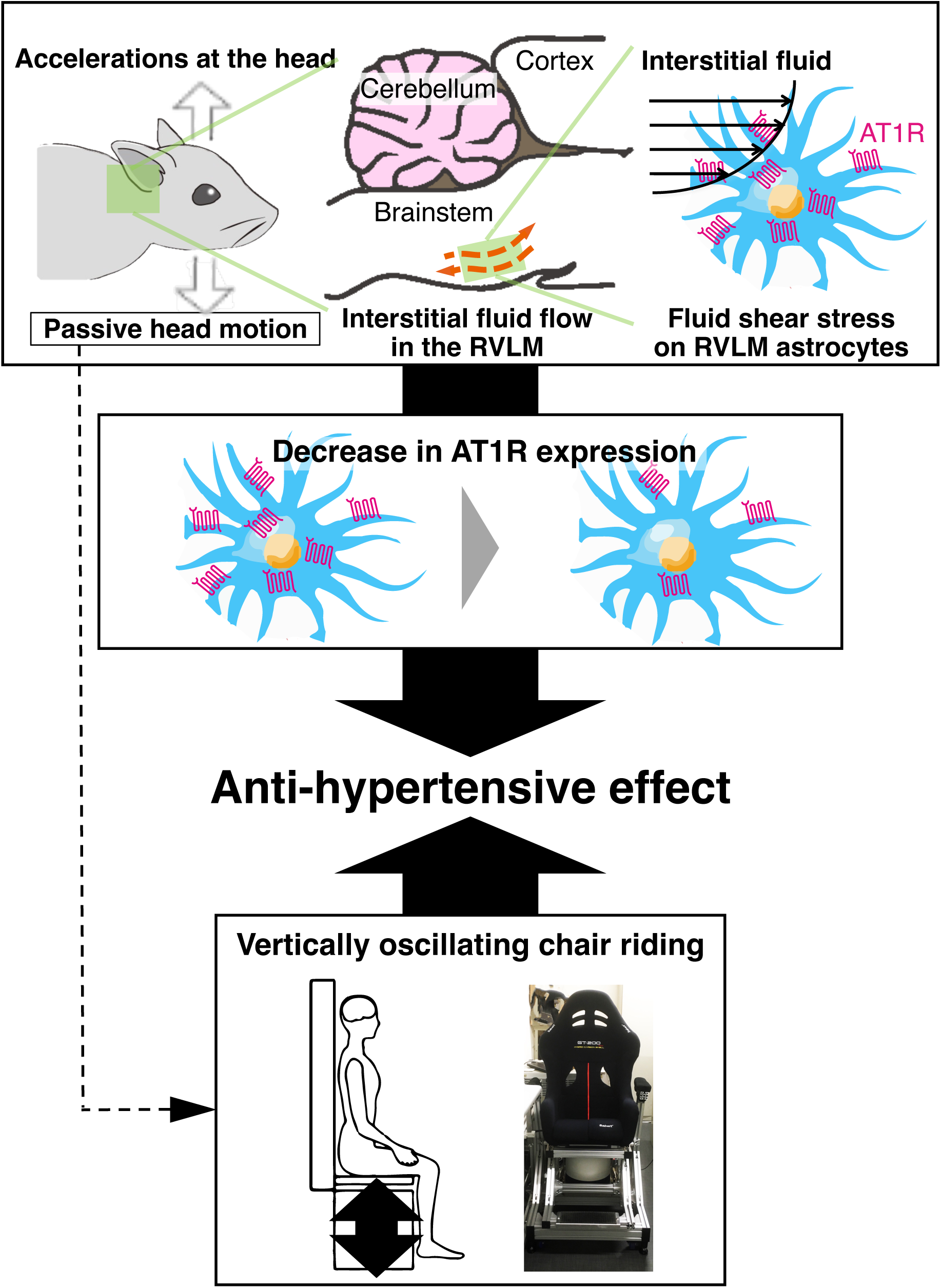
Schematic representation of the antihypertensive effects observed in PHM of hypertensive rats and VOCR of hypertensive humans. The results from our animal experiments indicate that cyclical application of mechanical intervention to the head generates interstitial fluid movement in the RVLM, leading to FSS-induced decrease in the AT1R expression in astrocytes in situ, and thereby ameliorates hypertension. Our studies also show that the VOCR of hypertensive adult humans, which produces vertical accelerations at their heads, lowers their BP.

**Supplementary Table 1.** **Calculation of the magnitude of PHM-generated FSS on rat RVLM cells. a,** Values referenced for calculation of the magnitude of FSS that PHM generated in the rat RVLM. Based on the estimated structure of the interstitial space in the rat RVLM (Extended Data Fig. 6e and Extended Data Fig. 7d), we referenced 4.5−5.5 as the Kozeny constant^80–82^ to calculate the Darcy permeability (see Methods). Because the velocity of interstitial fluid movement has been reported to be ∼0.2 μm/s in sedentary rats^37^ and mice^38^, we entered 0.4–0.6 μm/s as its value during PHM, based on our interpretation of the μCT study (i.e., two- to three-fold increase in velocity; Fig. 4g,h). The property of the interstitial fluid in the brain, whose composition is similar to that of the cerebrospinal fluid (CSF)^35^, was referenced from previous studies describing the viscosity of human CSF^83, 84^. The diameter of the individual interstitial space was referenced from the values of the cross-sectional area of the interstitial space as shown in Extended Data Fig. 7c. **b,** Calculation of the magnitude of FSS generated by PHM. The Darcy permeability (*K*_*p*_) is calculated as 6.63 × 10^-19^ − 1.45 × 10^-17^ m^2^. Following the calculation reported previously^85^, the magnitude of FSS (τ) at the cell surface is estimated as 0.076−0.53 Pa.

| Property | Value |
| --- | --- |
| Porosity ($\varepsilon$) | 0.2 (20%) |
| Diameter of interstitial space ($D_p$; $\mu\text{m}$) | 0.10–0.48 |
| Kozeny constant ($k$) | 4.5–5.5 |
| Viscosity of interstitial fluid ($\mu$; $\text{mPa}\cdot\text{s}$) | 0.72 |
| Velocity of interstitial fluid movement (flow) ($\mu\text{m/s}$) | 0.4–0.6 |

FSS ( $\tau_x$ ) at the cell surface:
, where $K_p$ is the Darcy permeability in the brain (RVLM).
When the values listed in **a** are introduced in these equations, the Darcy permeability is calculated as $6.63 \times 10^{-19} - 1.45 \times 10^{-17} \text{ m}^2$ , and the magnitude of FSS is estimated as **0.076–0.53 Pa**.

**Supplementary Table 2.**
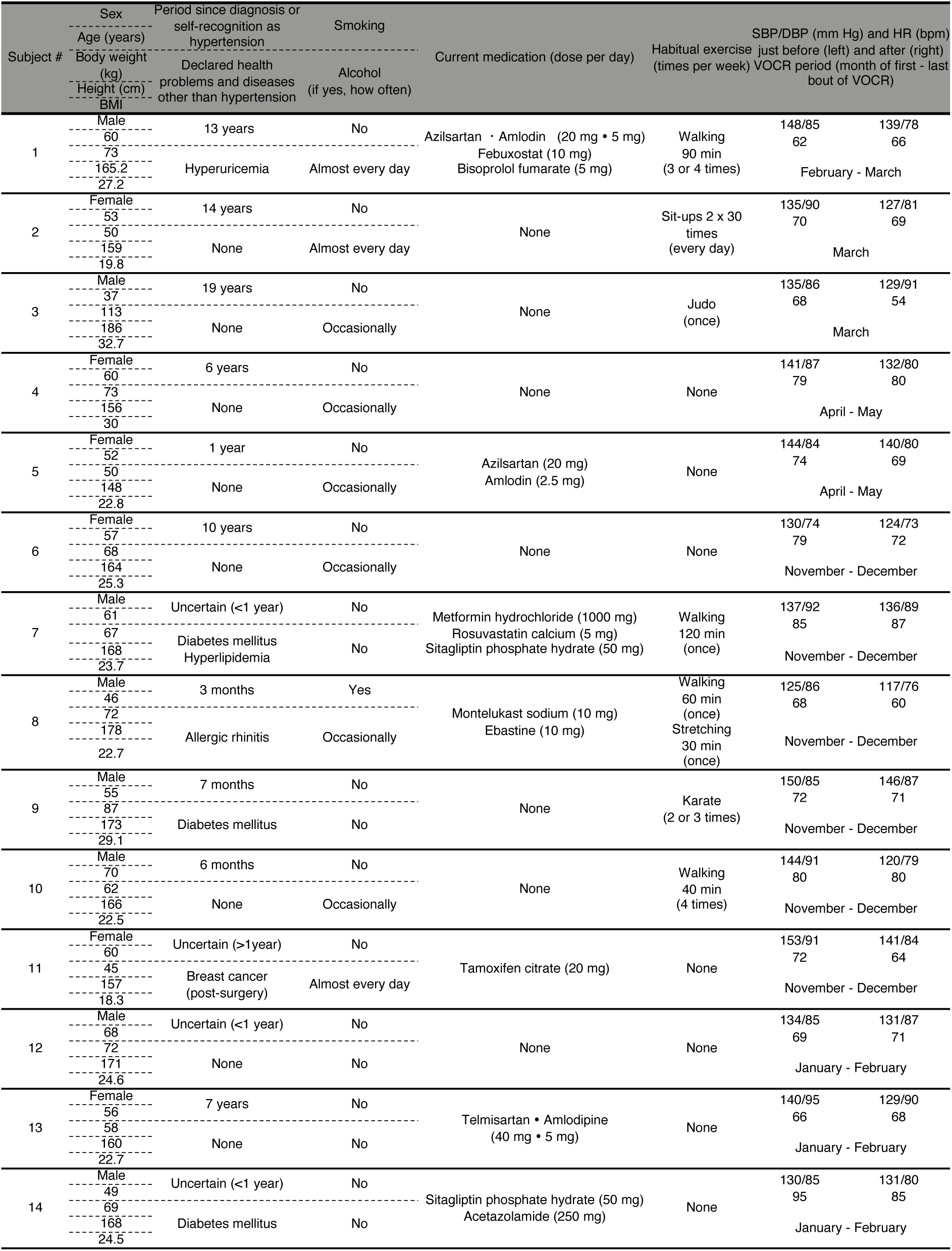
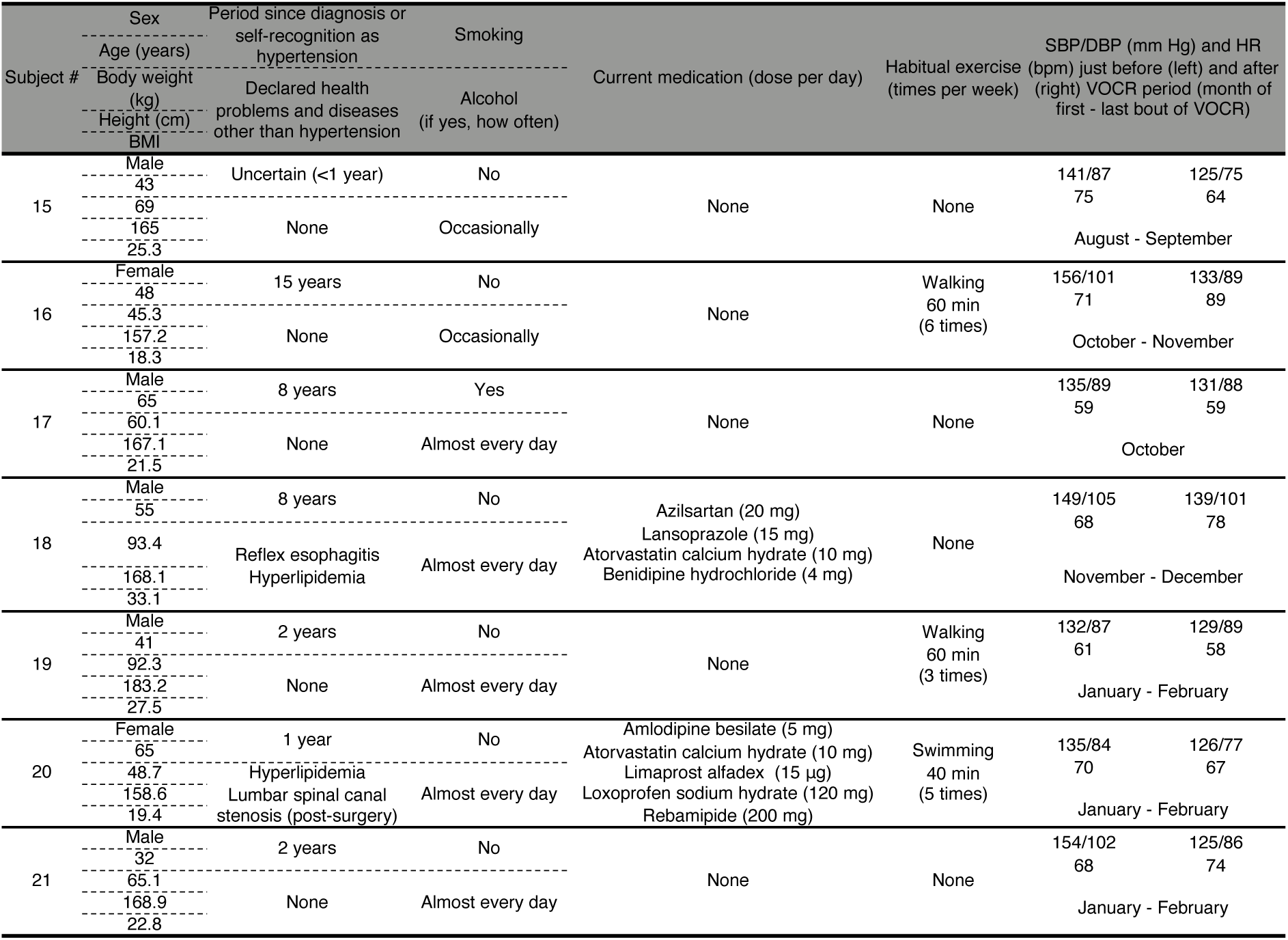
Information of subjects who participated in the human studies of protocol 1 and 2. SBP/DBP (mm Hg) and HR (bpm) just before and after VOCR are “value of the day”s in subjects #1−#5 (protocol 1) and “value of the week”s in subjects #6−#21 (protocol 2). Subject 9 was excluded from our statistical analysis because of the high serum CRP value before VOCR (2.85 mg/dL), which made it difficult to rule out acute infection, a possible disqualifier, at the time of the initiation of VOCR, albeit the lack of specific complaint or local symptom related to acute physical problem(s). His serum CRP after the VOCR period was within the normal range (0.12 mg/dL).

**Supplementary Table 3.**
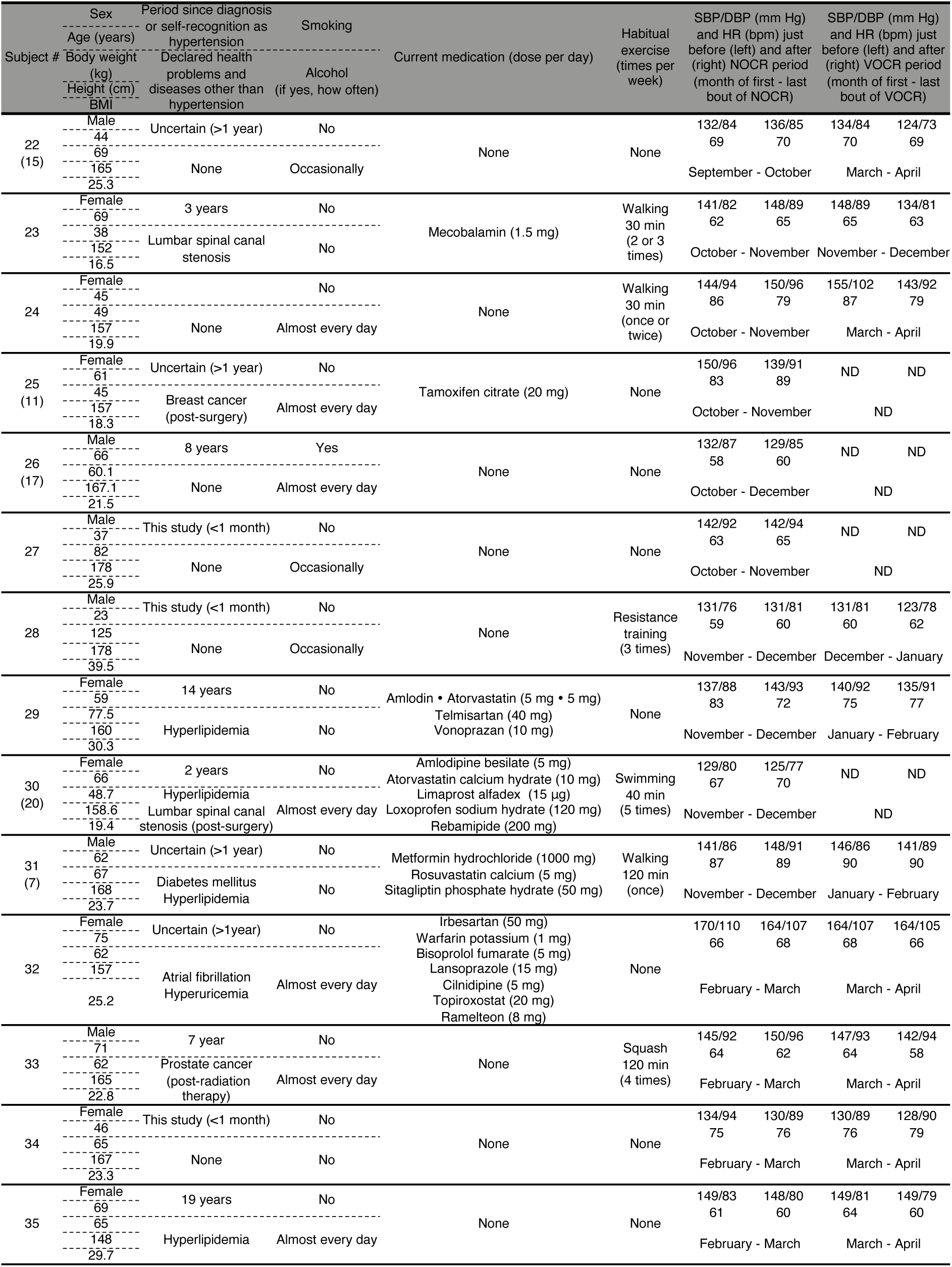
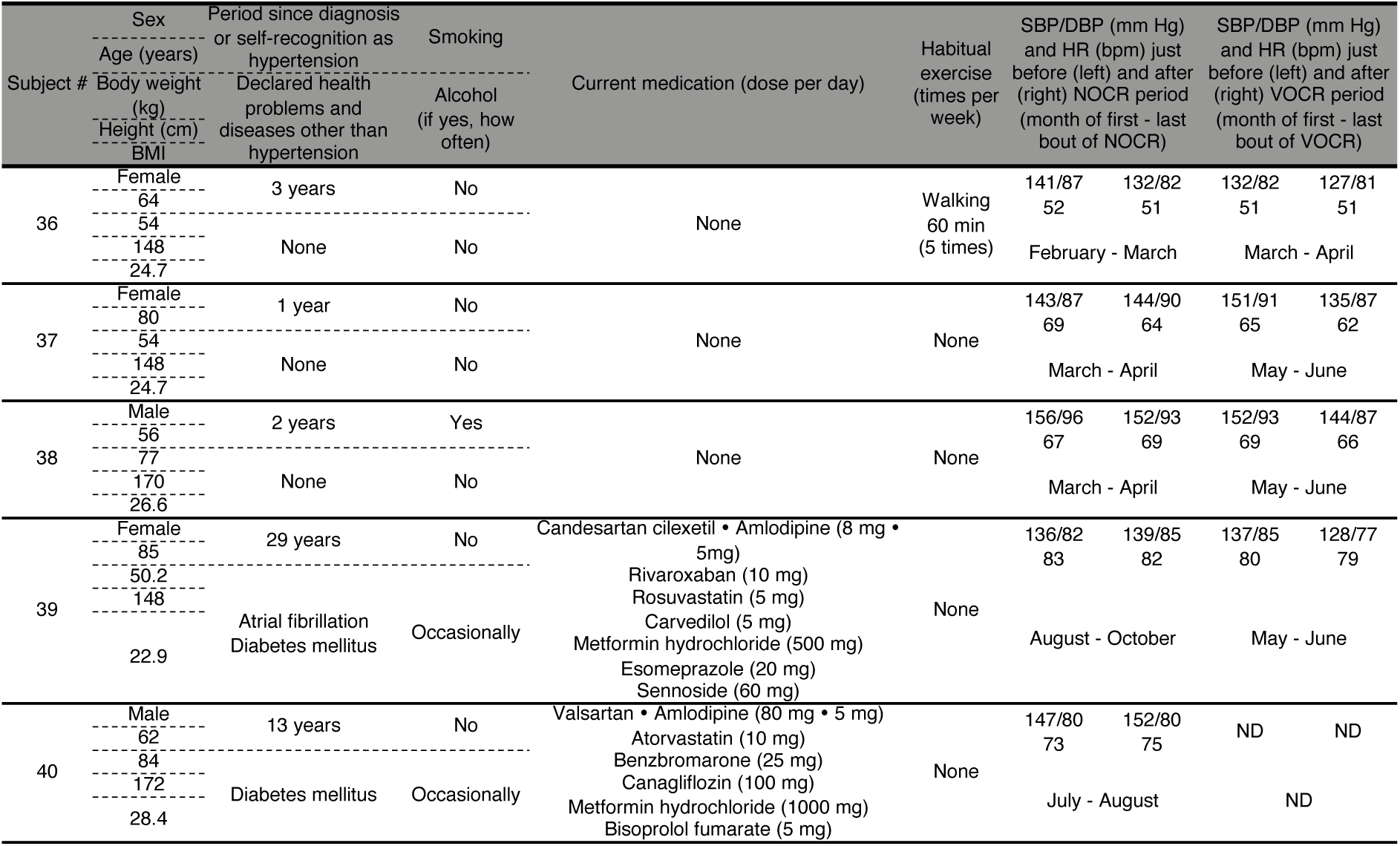
Information of subjects who participated in the study of protocol 2. SBP/DBP (mm Hg) and HR (bpm) just before and after NOCR or VOCR are “value of the week”s. Five of 19 subjects in protocol 3 were participants in protocol 2. The parenthesized subject numbers indicate their subject numbers in protocol 2 (see Supplementary Table 2). They participated in protocol 3 upon their eligibility check at least nine months after the last bout of VOCR in protocol 2.

## Methods

### Animal experiments and human studies

Animals were housed under a 12/12 h light-dark cycle with controlled temperature (23−25°C), and treated with humane care under approval from the Animal Care and Use Committee of National Rehabilitation Center for Persons with Disabilities (approval number: 30-07). Male SHRSP/Izm and WKY/Izm rats were provided from the Disease Model Cooperative Association (Kyoto, Japan) and astrocyte-GFP mice (Aldh1L1-GFP mice)^40^ were obtained from GENSAT (New York, NY), acclimated to the laboratory environments for at least 1 week, randomly divided into experimental groups, and used for experiments.

All participants in our human studies provided written informed consent. The studies were approved by the Ethics Committees of the Iwai Medical Foundation and the National Rehabilitation Center for Persons with Disabilities (approval number: 30-01).

### Chemicals and antibodies

All the chemicals were purchased from Sigma-Aldrich unless noted otherwise. Primary antibodies and their dilution rates used for immunostaining in this study are as follows: mouse monoclonal anti-GFAP (MAB360; Millipore, Billerica, MA) at 1:1,000; rabbit polyclonal anti-GFAP (Z0334; Dako, Glostrup, Denmark) at 1:1,000; chicken polyclonal anti-GFAP (ab4674; Abcam, Cambridge, UK) at 1:2,000; rabbit polyclonal anti-cleaved caspase- 3 (9661; Cell Signaling Technology, Danvers, MA) at 1:1,000; mouse monoclonal anti-NeuN (MAB377; Millipore) at 1:200; rabbit polyclonal anti-NeuN (ABN78; Millipore) at 1:1,000, rabbit polyclonal anti-AT1R (HPA003596; Sigma-Aldrich, St. Louis, MO) at 1:200; rabbit polyclonal anti-AGTRAP (HPA044120; Sigma-Aldrich) at 1:1,000; rabbit polyclonal anti-GFP (598; MBL, Nagoya, Japan) at 1:2,000; chicken polyclonal anti-GFP (ab13970; Abcam) at 1:2,000; mouse monoclonal anti-TUJ-1 (ab78078; Abcam) at 1:1,000. Secondary antibodies conjugated with Alexa Fluor 350, 488, 568, 633, and 647 (Thermo Fisher Scientific, Waltham, MA) were used at a dilution rate of 1:400. Cell nuclei were stained with DAPI (D9542; Sigma- Aldrich). Primary antibodies and their dilution rates used for immunoblot analysis are as follows: rabbit polyclonal anti-TNF-α (ab66579; Abcam) at 1:250; rabbit polyclonal anti-IL-1β (ab9722; Abcam) at 1:250; rabbit polyclonal anti-GAPDH (5174; Cell Signaling Technology) at 1:10,000. Horseradish peroxidase-conjugated anti-rabbit IgG (H + L) secondary antibody (W401B; Promega, Madison, WI) was used at a dilution rate of 1:5,000 for anti-TNF-α or -IL- 1β blotting, and 1:10,000 for anti-GAPDH blotting.

### PHM application to rats

Rats were subjected to PHM in a prone position using a platform that we developed to move their heads up and down^24^ (schematically represented in Fig. 1a). During PHM, animals were kept anesthetized with 1.5% isoflurane except for the µCT study, in which we used intraperitoneal injection of 2 mg/kg of midazolam (Sandoz, Basel, Switzerland), 2.5 mg/kg of butorphanol (Meiji Seika, Tokyo, Japan) and 0.15 mg/kg of medetomidine (Kyoritsu Seiyaku, Tokyo, Japan) for anesthesia. Body temperature of tested animals was maintained using a light heater. The PHM system was set up to reproduce the head motion (5 mm, 2 Hz) of treadmill running (20 m/min) which made 1.0 × *g* vertical acceleration peaks in the head of rats examined^24^. The control rats in PHM experiments were anesthetized likewise, and placed in a prone position with their heads on the platform that was left static.

### Treadmill running of rats

Rats were subjected to compulsive running using a belt drive treadmill equipped with an electrical shock system (MK-680S; Muromachi, Tokyo, Japan). We habituated the rats to the treadmill system by placing them in the machine several times without turning on the treadmill belt during the acclimation period. The electrical stimulation was turned on only once or twice during the first 5 min of the 30-min treadmill running on the first day of the 4-week treadmill running period. Thereafter, we did not need to turn on the electrical shock system to have the animals keep running, perhaps because the velocities we employed (20 m/min) were moderate. The control rats in treadmill running experiments were placed on the belt daily for 30 min without turning on the treadmilling.

### Measurement of BP and HR of rats by radio-telemetry

A telemetry pressure probe equipped with a micro electro mechanical systems (MEMS)-based 2.0-mm-long 0.47-mm-wide 0.60-mm thick sensor (Millar, Houston, TX) was implanted in the abdominal aorta of a rat at 9−10 weeks of age, following the surgical procedure described previously^86^. Rats were allowed to recover for at least 14 days before the initiation of experimental interventions or analyses. During the periods of experimentation that involved repeated BP and HR measurements over multiple weeks, BP and HR were monitored and recorded for continuous 30 min every 7 days between 10:00 AM and 12:00 PM (noon) by a multichannel amplifier and signal converter (LabChart 8; ADInstruments, Bella Vista, Australia).

### Measurement of urinary norepinephrine excretion of rats

Urine excreted during the indicated 24-h period was collected under an acidic condition using a metabolic cage (KN-646; Natsume Seisakusho, Tokyo, Japan) connected to a glass flask containing 10 mL of 6 N HCl, and stored at −80°C until assayed. Excretion of urinary norepinephrine was calculated by multiplying its concentration measured using an enzyme-linked immunosorbent assay (ELISA) kit (KA1891; Abnova, Taipei, Taiwan) with the urine volume.

### Tissue preparation and immunostaining (immunohistochemical or immunocytochemical analysis)

Rats were anesthetized with intraperitoneal injection of midazolam, butorphanol, and medetomidine, and perfused transcardially with 4% paraformaldehyde (PFA; TAAB Laboratories Equipment, Aldermaston, UK) in phosphate-buffered saline [PBS (137 mM NaCl, 10 mM Na_2_HPO_4_, 2.7 mM KCl, 1.5 mM KH_2_PO_4_)]. The brainstems were excised and post-fixed with 4% PFA in PBS overnight at 4°C. The tissues were cryoprotected by soaking in 20% sucrose/PBS for 24 h and in 30% sucrose/PBS for additional 24 h at 4°C. Fixed brainstems were frozen in optimal cutting temperature compound (Sakura Finetek, Tokyo, Japan) and cut into 16 μm-thick coronal sections using a cryostat (CM3050S; Leica Microsystems, Wetzlar, Germany). Sliced sections were permeabilized and blocked with 0.1% Tween-20 and 4% donkey serum (Merck Millipore, Burlington, MA) in Tris-buffered saline, blocked with, and stained by incubating with primary antibodies at appropriate dilutions followed by their species-matched secondary antibodies. Cell nuclei were stained with DAPI (Sigma-Aldrich). The slides were mounted with ProLong Gold Antifade Reagent (Thermo Fisher Scientific) and images were captured with a BZ-9000 digital microscope system (Keyence, Osaka, Japan). Quantitative immunohistochemical analysis (i.e., cell counting) of the RVLM was performed in the approximately same relative section for each rat (11.3–12.3 mm caudal to the bregma, 1.5–2.5 mm lateral to the midline, 0–1 mm dorsal to the ventral surface of the medulla).

For immunocytochemistry, cultured cells were fixed with 4% PFA in PBS for 20 min at room temperature (RT) and permeabilized and blocked with 0.1% Triton X-100 and 10% fetal bovine (FBS; GE Healthcare Life Science, Marlborough, MA) in PBS for 30 min at RT. The cells were then incubated with primary antibodies for 2 h and then with secondary antibodies for 1 h at RT.

### Microinjection into the rat RVLM

Rats were anesthetized with intraperitoneal injection of midazolam, butorphanol, and medetomidine except for Ang II or valsartan injection studies, in which we used 1.2−1.4 g/kg of urethane (Sigma-Aldrich), and subjected to microinjection as described previously^87^. In brief, a 25s-G microsyringe (Hamilton, Bonaduz, Switzerland) was stereotaxically positioned on anesthetized rats after exposure of dorsal surface of medulla. The needle placement was defined according to an atlas of the rat with stereotaxic coordinates^88^ ^63^; anteroposterior angle: 18°, 1.8 mm lateral to the calamus scriptorius, 3.5 mm ventral to the dorsal surface of the medulla. The placement of the needle tip in RVLM was confirmed by ensuring the pressure response to a test-dose injection of L-glutamate^12, 89^ (100 nL of 1 mmol/L in PBS). Microinjection of various compounds or mediums was made through a needle reinserted at the same coordinates with fixed infusion rates using a microsyringe pump instrument (KD scientific, Holliston, MA). Except for the experiments to analyze the pressor or depressor responses, we held the syringe for 5 min after the injection to avoid reflux, pulled out the needle carefully, and sutured the skin. The volumes and rates of microinjection were as follows; Ang II (Auspep, Tullamarine, Australia) and valsartan (Tocris Bioscience, Bristol, UK): 100 nL of 1 mmol/L in PBS at 0.1 µL/min, AAV solutions: 300 nL at 0.03 µL/min, PEG solutions: 1 µL at 0.1 µL/min, Isovist: 1 µL at 0.2 µL/min.

### Analysis of pressor/depressor responses

Rats implanted with a telemetry pressure probe were anesthetized with urethane, and subjected to the analysis of pressor/depressor responses. Monitoring the BP, we injected Ang II or valsartan (100 pmol) stereotaxically into the unilateral RVLM following the microinjection procedures described above. The injection side (right or left) was chosen randomly. When both pressor and depressor responses were analyzed, at least 2 h elapsed between the injections of Ang II and valsartan (Fig. 2a). The maximal MAP change elicited from the baseline was statistically analyzed^60^.

### Production of AAV vectors

To obtain astrocyte- and neuron-specific transduction, we used AAV9 vectors that expressed a transgene under the control of mouse GFAP and rat NSE promoters, respectively. The astrocyte-specific GFAP promoter consists of 0.6-kb hybrid fragments containing ABC1D genomic regions upstream of the mouse *GFAP* gene^90^. The neuron-specific NSE promoter is composed of the 1.2-kb genomic region upstream of the rat *NSE* gene^91^. Full-length rat AGTRAP cDNA was synthesized (Eurofins Genomics, Tokyo, Japan) and inserted into plasmid pAAV-GFAP-GFP-P2A-Cre-woodchuck hepatitis virus post- transcriptional regulatory element (WPRE)-simian virus 40 polyadenylation signal (SV40pA) and pAAV-NSE-GFP-P2A-Cre-WPRE-SV40pA to generate pAAV-GFAP-GFP-P2A-AGTRAP- WPRE-SV40pA and pAAV-NSE-GFP-P2A-AGTRAP-WPRE-SV40pA. pAAV-GFAP-GFP-WPRE-SV40pA and pAAV-NSE-GFP-WPRE-SV40pA were used for experimental controls. Recombinant single-strand AAV2/9 vectors were produced by transfection of HEK293T cells (Thermo Fisher Scientific) with the respective pAAV expression plasmid, pAAV2/9 and a helper plasmid (Stratagene, La Jolla, CA) as previously described^92^. After harvesting conditioned medium, the viral particles were precipitated using polyethylene glycol 8000 and iodixanol continuous gradient centrifugation as previously described^93^. The genomic titer of purified AAV9 vectors was determined by qPCR that targeted the WPRE sequence.

### Measurement of pressure in the rat medulla

Intramedullary pressure was measured using a blood pressure telemeter following the procedure we previously reported^24, 94^. To place the pressure sensor in the rat medulla, we made a ∼1-mm sized hole in the occipital bone (2 mm lateral to the midline, 2 mm rostral to the caudal margin of occipital bone) using a dental handpiece (Osada Electric, Tokyo, Japan), stereotaxically inserted a 20-G needle (Terumo, Tokyo, Japan) as a guide sheath (4 mm in depth from the occipital bone surface) at an anteroposterior angle of 20°, and sealed it airtightly using dental cement (GC, Tokyo, Japan) to avoid pressure escape. We macroscopically observed that the needle tip was placed in the rat RVLM after these procedures in our pilot experiments. However, for this pressure measurement, we did not strictly confirm the placement of the needle tip in the RVLM using the aforementioned test-dose injection of L-glutamate. Therefore, we designate this experiment as an “intramedullary” pressure measurement. During the pressure measurement, respiration was monitored using a pulse transducer (TN1012/ST; ADInstruments) attached to the tested rats. Low-pass (50 Hz) filtered intramedullary pressure waves were analyzed using LabChart 8 software.

### In vivo analysis of the distribution dynamics of interstitial fluid in the rat RVLM using μCT

Isovist (Bayer, Berlin, Germany) was stereotaxically microinjected into the RVLM of anesthetized 12-week-old male WKY rats following the procedure described above, and visualized using µCT (inspeXio SMX-100CT; Shimadzu, Kyoto, Japan). After Isovist injection, rats were subjected to two serial brain µCT scans between which PHM was either applied or left unapplied (kept sedentary) for 30 min (Fig. 4e). µCT images were analyzed using software for 3-D morphometry (TRI/3D-BON-FCS64; RATOC System, Tokyo, Japan). Voxels with ≥1.02 times signal intensity as compared with that of air was defined as Isovist cluster in the rat RVLM.

### Hydrogel introduction in the rat RVLM

Just before use, a pre-mixture of PEG with functional groups (25 g/L in PBS) was prepared from tetra-armed thiol-terminal (TetraPEG-SH) (Yuka-Sangyo, Tokyo, Japan) and acrylate-terminal (Tetra-PEG-ACR) (JenKem Technology, TX, USA) PEG solutions as we previously described^24^. Tetra-armed polyethylene glycol without functional groups (25 g/L in PBS) was used as an ungelatable control. For the analysis of hydrogel distribution in rat RVLM, we used Tetra-PEG-SH fluorescently labeled with a thiol-reactive probe (Thermo Fisher Scientific or Merck KGaA, Darmstadt, Germany). Microinjection of PEG solutions into rat RVLM was conducted as described above.

To specifically analyze the consequences of PHM and hydrogel introduction by minimizing possible invasive influences of microinjection itself, we gave 1-week recovery time before the first BP measurement, and then applied PHM to the rats (daily 30 min, 14 or 28 days). Immediately subsequent to post-PHM 24-h urine collection (Fig. 6a), rats were sacrificed by transcardial infusion of PFA and subjected to histological analysis.

### Multiphoton microscopic imaging and analysis of the interstitial space structure/orientation and dimension in the rat RVLM

We prepared PFA-fixed lower brainstem tissue samples from 12-week-old WKY rats 24 h after introducing fluorescently labeled hydrogel conjugated with Alexa Fluor 594 C5 maleimide (A10256, Thermo Fisher Scientific) to their RVLM as described above. We used a multiphoton microscope (FVMPE-RS- UPSP2; Olympus, Tokyo, Japan), which is suitable for viewing a deep part of a relatively large sample. This was because we wanted to minimize the possible influence of plastic deformation/distortion or disruption of hydrogels (and tissues) that might occur during the experimental procedures (e.g., sample sectioning) particularly in the vicinity of the cut section. The sample was partially embedded in 2% (w/v) agarose (A9414; Sigma-Aldrich) and immersed in PBS. An excitation laser (Mai Tai DeepSee; Spectra-Physics, Milpitas, CA) was tuned at 800 nm, and the beam was focused with an objective lens (XLPLN25XWMP2; Olympus). The fluorescence was collected with the same objective lens, and detected using a GaAsP detector after passing through a red band pass filter (BA575-645; Olympus). Considering the relative extension of multiphoton excitation in the incident direction of excitation laser beam^95^, which is parallel to the optical axis of objective lens, we acquired and analyzed images of the three mutually orthogonal planes (*xy*-, *xz*-, and *yz*-planes; Extended Data Fig. 6a-c and Extended Data Fig. 7a-c).

Image analysis to determine the structure/orientation of the interstitial space was conducted as follows. Approximately 20-μm stacked long-rectangular voxel images (20 voxels of 1-μm length) were resampled to 100 cubic voxels (0.2072 μm on each side), and smoothed to a width of 5 voxels (∼1 μm). Following the previous reports on the occupancy of interstitial space in the brain^74, 96^, we extracted the top 20% high-fluorescence voxel population as clusters representing the interstitial space (Extended Data Fig. 6a, bottom). For huge clusters (>5,000 voxels), the bottom 50% population of low-fluorescence voxels in them were removed to reduce their size. These procedures, together with a clearance of clusters of <50 voxels enabled us to apply elliptic fitting to determine the major and minor axes of each processed cluster on the stacked plane (Extended Data Fig. 6b). For each sample, we analyzed 100 slices, focusing on the plane with the highest mean fluorescence intensity along the stack direction.

To quantitatively analyze the cross-sectional area of the interstitial space, we resampled ∼4-μm stacked long-rectangular voxel images to 20 cubic voxels (Extended Data Fig. 7a). Based on the voxel number of each individual cluster (Extended Data Fig. 7b), we drew the distribution of its cross-sectional area (Extended Data Fig. 7c). Analyzing the probability density by fitting to the log-normal distribution^77, 78^, we determined the mode and full width at half maximum (FWHM)^79^ of the cross-sectional areas of the individual interstitial spaces (Extended Data Fig. 7c). We used MATLAB software (version 2021a; Math Works, Natick, MA) for these image analyses.

### MRI scanning and orientation analysis of the rat lower brainstem

Twelve-week-old male WKY rats were scanned in a 7.0-Tesla MRI system (Biospec 70/30-USR; Bruker BioSpin, Ettlingen, Germany) using a 2-D-rapid acquisition with relaxation enhancement (RARE).

Parameters for the sagittal T_2_-RARE sequence were as follows: echo time (TE), 33 ms; repetition time (TR), 3600 ms; slice thickness (SL), 0.75 mm; field of view (FOV), 38.4 × 38.4 mm; voxel, 0.15 × 0.15 × 0.75 mm; RARE factor, 8; number of averages, 4. We analyzed the MR images using SPM12 on MATLAB (https://www.fil.ion.ucl.ac.uk/spm/software/spm12/). Each image was smoothed with 0.24-mm FWHM, and registered to the template atlas of the Fischer 344 rat brain^97^ (An MRI-Derived Neuroanatomical Atlas of the Fischer 344 Rat Brain (Version v4) [Data set]. Zenodo. https://doi.org/10.5281/zenodo.3900544). Assuming an equivalent weight for each voxel, we determined the center of mass of the lower brainstem in each image in a particular plane (*xy*-, *xz*- or *yz*-plane) and drew its centroidal line by combining the least squares and sigmoid fitting. Because the *y*-axis was actually the direction of the maximum diameter of this part of the brain, the longest centroidal line was drawn by fitting to the mass centers determined in the *xz*-plane images (Extended Data Fig. 6d).

### Calculation of the magnitude of FSS on the cells in the rat RVLM during PHM

We calculated the interstitial fluid flow-derived FSS imposed on cells in the rat RVLM during PHM by referring to the findings from our μCT analysis (Fig. 4g,h) and multiphoton microscopy analyses (Extended Data Fig. 6 and 7) according to Henry Darcy’s law, which defines the flux density of penetrating fluid per unit time^98^. Based on the PHM-induced enhancement of Isovist spread (Fig. 4g,h), we estimated the velocity of the interstitial fluid flow (*u*) during PHM as 0.4−0.6 μm/s, two- to three-times that in the brain of sedentary rodents, which is reported to be 0.2 μm/s^37, 38^. We then calculated the Darcy permeability (*K*_*p*_), which relates to the flow or pressure drop of liquid across a porous structure^99^, in the rat RVLM tissue. The Kozeny–Carman equation allows us to calculate permeability given pore-size/sphericity data^80, 81^, where a value of 4.5−5.5 was considered the Kozeny constant^80–82^ (Supplementary Table 1a) based on the apparent structure/orientation of the interstitial space in the rat RVLM (Extended Data Fig. 6e and Extended Data Fig. 7d). We referenced the interstitial space dimension (cross-sectional area or diameter) estimated by analyzing the microscopic images (Extended Data Fig. 7c). FSS (τ) at the interstitial cell surface was calculated as described in Supplementary Table 1^85^.

### Cell culture

Primary cultures of astrocytes were prepared from the cortex of neonatal (one day old) mouse pups as described previously^100, 101^ with some modifications. The brains were excised from astrocyte-GFP mice^40^ immediately after euthanasia by CO_2_ inhalation (typically 4−5 pups at one time), and the cortices were dissected to small pieces in DMEM (FUJIFILM Wako Pure Chemical, Osaka, Japan) using forceps under a stereomicroscope (SV-11; Zeiss, Oberkochen, Germany). The dissected pieces of cortex tissue were collected by 190-×-*g* centrifugation for 3 min, suspended, and incubated in an isotonic solution (124 mM NaCl, 5 mM KCl, 3.2 mM MgCl_2_, 0.1 mM CaCl_2_, 26 mM NaHCO_3_, 10 mM D-glucose, 100 I.U./mL penicillin, 100 µg/ml streptomycin) containing 0.1% trypsin, 0.67 mg/ml hyaluronidase, and 0.1 mg/ml deoxyribonuclease I at 37°C for 10 min to dissociate cells. Subsequently, an equal volume of DMEM/F12 (Thermo Fisher Scientific) containing 10% FBS, 100 I.U./mL penicillin, and 100 μg/mL streptomycin was added to neutralize trypsin. After multiple pipetting for further homogenization with a 10-mL serological pipet, the cell suspension was cleared of tissue debris using a 70-µm nylon-mesh strainer (Corning Life Sciences, Corning, NY). Cells were then washed twice with DMEM/F12 containing 10% FBS, 100 I.U./mL penicillin, and 100 μg/mL streptomycin, plated on a poly-D-lysine-coated T75 flask (Corning Life Sciences, Corning, NY), and kept in a humidified incubator (5% CO_2_, 95% air, 37°C). The culture medium was replaced with fresh one every 3 days, and it typically took 7−10 days for the cells to become confluent in the flask. Then, astrocyte-enriched cell population was obtained by physically detaching the other types of cells, such as microglia and oligodendrocytes. After shaking the flask overnight using an orbital shaker (200 rpm with BR-40LF; TAITEC, Koshigaya, Japan), detached cells were removed together with the medium. Cells remained attached to the flask were replated on poly-D-lysine-coated 10-cm dishes (Corning Life Sciences) at a ratio of 1:3 regarding the density (typically 3−4 dishes). The medium was replaced with fresh one every 3 days until the cells reached 80−90% confluence and were used for experiments. We confirmed that >95% of the cells were GFP-positive (Extended Data Fig. 8a), indicating a high degree of astrocyte purity.

Mouse neuroblastoma-derived Neuro2A cells (provided from Dr. Yokota, Tokyo Medical and Dental University, Japan), which exhibit neuronal phenotypes and morphology^41, 42^, were cultured in DMEM supplemented with 10% FBS, 100 I.U./mL penicillin and 100 μg/mL streptomycin in a humidified incubator (5% CO_2_, 95% air, 37°C).

### Application of FSS to astrocytes or Neuro2A cells in culture

Astrocytes or Neuro2A cells grown in a poly-D-lysine-coated 35-mm culture dish (Corning Life Sciences, Corning, NY) were exposed to pulsatile FSS (average 0.7 Pa) for 30 min. As we previously reported^23, 24, 34^, a parallel plate flow-chamber and a roller pump (Masterflex; Cole-Parmer, Vernon Hills, IL) were used to apply FSS. The flow-chamber, which was composed of a cell culture dish, a polycarbonate I/O unit, and a silicone gasket, generated a 23-mm-long 10-mm-wide 0.5-mm-high flow channel. Astrocytes or Neuro2A cells were seeded at a density of 8 × 10^5^ cells/8.0 cm^2^. To maintain pH and temperature of culture medium, we used a 5% CO_2_-containing reservoir and a temperature-controlled bath.

### Application of HPC to astrocytes in culture

Astrocytes seeded at a density of 8 × 10^5^ cells/8.0 cm^2^ were exposed to cyclical HPC with various amplitudes at a frequency of 0.5 Hz for 30 min using a custom-made pressure system described previously^34^. The system consists of a cell culture dish, a polycarbonate pressure chamber, a silicone gasket, an O-ring, a quartz glass, two holding jigs, a thermostatic chamber, and a syringe pump. The system was completely airtight, enabling precise and strict pressure control with the syringe pump. To maintain the temperature of culture medium, the entire system was placed in a 37°C incubator during HPC application. In both FSS and HPC experiments, cell adhesion and morphology were observed using a light microscope (DM IRE2; Leica Microsystems, Wetzlar, Germany).

### Quantitative PCR Analysis (reverse transcription and real-time PCR)

500 ng of total RNA extracted from cell culture were subjected to reverse transcription, using ISOGEN II (NIPPON GENE, Tokyo, Japan) and PrimeScript RT reagent Kit (TaKaRa, Kusatsu, Japan). The resulting cDNA was subjected to real-time PCR analysis using glyceraldehyde-3-phosphate dehydrogenase (GAPDH) as an internal control in Applied Biosystems 7500 Real Time PCR System with Power SYBR Green PCR Master Mix (Thermo Fisher Scientific).

The primers (sense and antisense, respectively) were as follows: mouse *Agtr1a* (AT1R-encoding gene); 5’-AAAGGCCAAGTCGCACTCAAG-3’ and 5’-TCCACCTCAGAACAAGACGCA-3’, mouse *Gapdh* (GAPDH-encoding gene); 5’-GCAAAGTGGAGATTGTTGCCAT-3’ and 5’-CCTTGACTGTGCCGTTGAATTT-3’, WPRE (for genomic titration of purified AAV9 vectors); 5’-CTGTTGGGCACTGACAATTC-3’ and 5’-GAAGGGACGTAGCAGAAGGA-3’.

### Fluorescent Ang II binding assay

Six or twenty-four hours after the termination of FSS application, cultured astrocytes were incubated with Ang II type 2 receptor inhibitor, PD123319 (10^-6^ mol/L in PBS; ab144564, Abcam), for 20 min, and then with tetramethylrhodamine (TAMRA)-labeled Ang II (10^-8^ mol/L in PBS; AS-61181, AnaSpec, Fremont, CA) for 30 min. After 3 times wash with PBS, samples were fixed and subjected to anti-GFP immunostaining to strengthen the GFP-derived green fluorescence signals and corroborate our analysis on astrocytes prepared from astrocyte-GFP mice as well as to secure the binding of fluorescent Ang II. Green and red fluorescence was viewed with a fluorescence microscope (BZ-9000 HS; Keyence). Samples from astrocytes left unexposed to FSS were prepared and viewed likewise, and provided an experimental control.

### Terminal deoxynucleotidyl transferase-mediated dUTP nick-end labeling (TUNEL) assay

Rat RVLM sections were stained using a TUNEL kit (Biotium, Fremont, CA) according to the manufacturer’s protocols, counterstained with DAPI, and then viewed using a 20× objective with a fluorescence microscope (BZ-9000 HS; Keyence). The nuclei of apoptotic cells were determined by the existence of green fluorescent patches, and cell apoptosis was quantified by referring their counts to total numbers of nuclei defined by DAPI staining.

### Immunoblot analysis

Rat RVLM tissue was excised immediately after cervical dislocation, mechanically homogenized, solubilized with RIPA buffer (25 mM Tris•HCl pH 7.6, 150 mM NaCl, 1% NP-40, 1% sodium deoxycholate, 0.1% SDS) with protease and phosphatase inhibitors (78440; Thermo Fisher Scientific), and subjected to SDS-PAGE followed by anti-TNF-α, anti-IL-1β, and anti-GADPH immunoblotting. Specific signals were visualized and quantified using Odyssey infrared imaging system (LI-COR Biosciences, Lincoln, NE) and ImageJ software (NIH, Bethesda, MD).

### Analysis of rat ADNA and BP during the transition from before to after the initiation of PHM

ADNA recording was conducted in isoflurane (2%)-anesthetized 12−16-week-old male SHRSPs as described previously^102^ with some modifications. Briefly, using a stereomicroscope (M80; Leica, Heerbrugg, Switzerland), we surgically exposed and isolated the left aortic depressor nerve through an anterior neck approach. We then placed a bipolar cuff electrode below the nerve bundle and embedded the nerve-electrode contacting complex in a two- component silicone gel (932; Wacker Chemie, Munich, Germany). Multifiber ADNA was 50−10,000-Hz band-pass filtered and amplified with a differential preamplifier (MEG-5200; Nihon Kohden, Tokyo, Japan). The ADNA was identified using its unique spontaneous activity synchronous with the arterial systolic cycle as an indicator. BP was monitored with a pressure transducer (DX-360; Nihon Kohden, Tokyo, Japan)/polygraph system (UBS-100-6; Unique Medical, Tokyo, Japan) connected to a catheter inserted in the right femoral artery. The PHM cycle was also monitored using a piezoelectric pulse transducer (MLT1010; ADInstruments). ADNA, BP, and PHM cycle were digitized using Power1401 mkII and Spike2 software (Cambridge Electronic Design, Cambridge, UK) at a 3-kHz sampling frequency. The recorded ADNA was then subjected to DC offset removal, rectified, integrated and smoothed with a time constant of 3 ms by Spike2. The integrated value of ADNA measured after euthanasia by isoflurane overdose was considered background noise and subtracted from the entire data collected. ADNA was then normalized in each rat (presented in a.u. in Extended Data Fig. 14a). The relative ADNA value (%) averaged for each heartbeat was scaled with the mean of 10 heartbeats immediately before the PHM initiation set as a baseline (100%) (Extended Data Fig. 14b). The SBP, DBP, MAP, and HR were determined using Spike2.

### Measurement of accelerations in the human head

To measure the accelerations at the human head during treadmill running or VOCR, we fixed an accelerometer (NinjaScan-Light; Switchscience, Tokyo, Japan) on foreheads with a surgical tape. Vertical acceleration was evaluated using the software application provided from the manufacture.

### Design and participants of clinical study on antihypertensive effects of vertically oscillating chair riding

We conducted single-arm (protocol 1 and 2) and non-randomized controlled (protocol 3) clinical studies. The study of protocol 1 (Extended Data Fig. 15c) was carried out at the affiliated health services facility of Iwai Medical Foundation (Iwai Keiaien, Tokyo, Japan).

The studies of protocol 2 (Extended Data Fig. 16a) and 3 (Extended Data Fig. 17a) were carried out at the National Rehabilitation Center for Persons with Disabilities Hospital. In protocol 3, participants were assigned to either NOCR or VOCR, depending on their intention and choice, whereas 14 individuals underwent both NOCR and VOCR after we obtained their consent (Supplementary Table 3). In principle for the “NOCR and VOCR” participants, NOCR intervention was administered prior to VOCR considering the antihypertensive effect of VOCR remaining for a while after its termination (Fig. 7b). In only one participant (#39), VOCR intervention was performed prior to NOCR. She first participated in the VOCR study upon her agreement and choice, and then agreed to the NOCR study after the completion of the VOCR intervention. We confirmed that her BP had stably returned to the pre-VOCR level before starting the NOCR intervention; her NOCR was initiated 10 weeks after the last bout of VOCR. Continuous beat-by-beat recording of BP and RRI was conducted for the participants who agreed to these measurements in protocol 3. Because the system for continuous BP recording became available after we started the protocol 3 study, two participants in the NOCR group (#22 and #23) were subjected to continuous RRI recording alone (i.e., void of continuous BP recording). Subjects #32 and #39 were excluded from continuous BP and RRI recording, despite agreeing to these measurements, because arrhythmia was revealed on electrocardiography (the first RRI recording) and confirmed to be atrial fibrillation by their primary care physicians (Supplementary Table 3).

Subjects were considered eligible if they were 20−75 (protocols 1 and 2) or ≥20 (protocol 3) years old of age and confirmed to have 130−160 mm Hg of SBP at the time of interview for informed consent and eligibility check. Subjects with mental or psychological illnesses, history or presence of cardiovascular events, history or presence of severe dysfunction/disorder of liver, kidney, lung, gastrointestinal tract, and spine, or presence of acute injuries/diseases (e.g., recent traumas and infectious diseases) were excluded with the exception of those who were given permission for participating in this study from their primary care physicians. Whereas antihypertensive medication did not disqualify the subjects (Supplementary Table 2 and 3), they were advised not to change their medication from at least one month prior to the first bout of NOCR or VOCR through the study period (i.e., up to 8 weeks after the last bout of NOCR or VOCR). At a certain (approximately fixed) time point in the morning (typically just before breakfast), they conducted 3 consecutive measurements of BP (mm Hg) and HR (bpm) using an automated upper arm-cuff sphygmomanometer, and recorded the values from all those measurements. These procedures of BP measurement and recording accord with the Japanese Society of Hypertension Guidelines for the management of hypertension (JSH2019)^103^. Subjects were directed to start periodical (≥3 days per week) BP/HR measurements at least 2 weeks (protocols 1 and 2) or 1 week (protocol 3) before the initiation of the intervention (i.e., NOCR or VOCR) and continue to measure BP/HR throughout the study period using the same sphygmomanometers. Particularly at the studies of protocol 2 and 3 (i.e., the studies at the National Rehabilitation Center for Persons with Disabilities), participants were advised to record all the data of those measurements. Those whose BP lowered below the eligibility requirement of the study (≥130 mm Hg of SBP) prior to the initiation of intervention (NOCR or VOCR) were eliminated from the study. Participants were directed to be rested and keep calm for at least 1 min before starting to measure BP/HR. The mean BP/HR value from 3- time measurements was defined as “value of the day”, and used for statistical analysis. When BP and HR were measured and recorded on ≥3 days during a particular week in the studies of protocol 2 and 3, the mean of all the “value of the day”s through the week was defined as “value of the week”. For the participants who agreed, periodical BP/HR measurement and recording (≥3 days per week) was extended up to 8 weeks after the last bout of VOCR (protocol 2 and 3). MAP was calculated with a standard formula as follows: MAP = DBP + 1/3 (SBP – DBP).

### Blood sampling and measurement of parameters in plasma and serum of humans

Blood sampling in the human study of protocol 2 was conducted between 12 PM and 3 PM. Participants were rested in a sitting position for at least 10 min before starting the sampling procedures. After plasma (for catecholamines and renin activity) and serum (for aldosterone and CRP) separation by centrifugation, we outsourced the measurement of parameters be tested (BML, Kawagoe, Japan).

### Continuous beat-by-beat BP and RRI recording to evaluate vascular sympathetic nerve activity and balance of cardiac sympathetic/parasympathetic nerve activity, respectively

In the study of protocol 3, we evaluated the vascular sympathetic nerve activity and the balance between the cardiac sympathetic and parasympathetic nerve activities (i.e., sympatho-vagal balance) in the participants who agreed to these measurements. We analyzed the LF (0.04–0.15 Hz) power of the power spectrum density (PSD) of SBP variability^52, 53^ and the LF/HF (0.15–0.4 Hz) ratio of the PSD of RRI variability^52, 54^, respectively, at the beginning and ending periods of the intervention (NOCR and VOCR) (Extended Data Fig. 17a). The PSDs of the SBP and RRI variability signals were calculated by applying a Fourier transform with Hanning window^104^. Continuous BP recording was performed using a continuous non-invasive arterial pressure system that measures BP via an automated inflatable cuff wrapped around the proximal phalanx of index or middle finger (LiDCOrapid V3; Merit Medical Japan, Tokyo, Japan)^105^. Continuous RRI recording was conducted using a wearable electrocardiography system (myBeat; Union Tool, Tokyo, Japan)^106^. Whilst we monitored BP and RRI continuously for 10 min at each recording, we analyzed the data from the central 256 s (4.27 min) of the last 5 min. It took us ∼5 min to prepare for continuous BP and RRI monitoring and confirm the proper system operation before starting BP and RRI recording. Therefore, the analyzed data were from the subjects who had rested on the chair for at least 10 min. The average values from five-time (i.e., 5-day) measurements at the beginning and ending periods of intervention (NOCR or VOCR) were statistically analyzed. In case the BP was not determined or SBP was recorded as <50 mm Hg, the measurement was considered a failure. For RRI analysis, we first removed RRIs of <0.5 s and >1.5 s as erroneous values. RRI values outside the range of 0.6- to 1.4-times the average of the 10 immediately preceding ones were also removed because of a lack of reliability. In case the sum of time for BP recording failures exceeded 4% or that for RRI value removal exceeded 10% during the last 5 min of recording, the continuous BP or RRI measurement on that particular day was deemed unsuccessful and excluded from the statistical analysis. Nevertheless, all statistically analyzed data were the mean values from at least four measurements except for the “End” RRI variability for subject #31’s NOCR, which was from three measurements because of two failures in recording.

### Statistical analysis

All the quantitative data are presented as mean ± s.e.m. Parametric statistical analyses were conducted by paired or unpaired two-tailed Student’s *t-*test for two- group comparison except for SBP and RRI variabilities, using Prism software (Version 8; GraphPad Software, San Diego, CA). SBP and RRI variabilities were analyzed by Wilcoxon signed-rank test, using R software (https://www.r-project.org/) in the human study of protocol 3. One-way ANOVA and one- or two-way repeated measures ANOVA with Tukey’s, Dunnett’s, or Bonferroni’s post hoc test were conducted for multiple (≥3) group comparison, using Prism. Differences were considered as significant at *P* values below 0.05.

## References

1. Hagberg, J.M., Park, J.J. & Brown, M.D. The role of exercise training in the treatment of hypertension: an update. Sports Med 30, 193–206 (2000).

2. Pescatello, L.S., MacDonald, H.V., Lamberti, L. & Johnson, B.T. Exercise for hypertension: A prescription update integrating existing recommendations with emerging research. Curr Hypertens Rep 17, 87 (2015).

3. Lim, S.S., et al. A comparative risk assessment of burden of disease and injury attributable to 67 risk factors and risk factor clusters in 21 regions, 1990-2010: a systematic analysis for the Global Burden of Disease Study 2010. Lancet 380, 2224–2260 (2012).

4. Carretero, O.A. & Oparil, S. Essential hypertension. Part I: definition and etiology. Circulation 101, 329–335 (2000).

5. Guyton, A.C. Abnormal renal function and autoregulation in essential hypertension. Hypertension 18, III49-53 (1991).

6. Guyenet, P.G. The sympathetic control of blood pressure. Nat Rev Neurosci 7, 335–346 (2006).

7. Grassi, G., Mark, A. & Esler, M. The sympathetic nervous system alterations in human hypertension. Circ Res 116, 976–990 (2015).

8. Malpas, S.C. Sympathetic nervous system overactivity and its role in the development of cardiovascular disease. Physiol Rev 90, 513–557 (2010).

9. Dampney, R.A. Functional organization of central pathways regulating the cardiovascular system. Physiol Rev 74, 323–364 (1994).

10. Fyhrquist, F. & Saijonmaa, O. Renin-angiotensin system revisited. J Intern Med 264, 224–236 (2008).

11. Forrester, S.J., et al. Angiotensin II signal transduction: An update on mechanisms of physiology and pathophysiology. Physiol Rev 98, 1627–1738 (2018).

12. Ito, S., Komatsu, K., Tsukamoto, K., Kanmatsuse, K. & Sved, A.F. Ventrolateral medulla AT1 receptors support blood pressure in hypertensive rats. Hypertension 40, 552–559 (2002).

13. Muratani, H., Ferrario, C.M. & Averill, D.B. Ventrolateral medulla in spontaneously hypertensive rats: role of angiotensin II. Am J Physiol 264, R388–395 (1993).

14. Kishi, T. & Sunagawa, K. Exercise training plus calorie restriction causes synergistic protection against cognitive decline via up-regulation of BDNF in hippocampus of stroke-prone hypertensive rats. Conf Proc IEEE Eng Med Biol Soc 2012, 6764–6767 (2012).

15. Nabika, T., Ohara, H., Kato, N. & Isomura, M. The stroke-prone spontaneously hypertensive rat: still a useful model for post-GWAS genetic studies? Hypertens Res 35, 477–484 (2012).

16. Lu, X., et al. Effects of local mechanical stimulation on coronary artery endothelial function and angiotensin II type 1 receptor in pressure or flow-overload. J Hypertens 31, 720–729 (2013).

17. Galie, P.A., Russell, M.W., Westfall, M.V. & Stegemann, J.P. Interstitial fluid flow and cyclic strain differentially regulate cardiac fibroblast activation via AT1R and TGF-β1. Exp Cell Res 318, 75–84 (2012).

18. Zou, Y., et al. Mechanical stress activates angiotensin II type 1 receptor without the involvement of angiotensin II. Nat Cell Biol 6, 499–506 (2004).

19. Ramkhelawon, B., et al. Shear stress regulates angiotensin type 1 receptor expression in endothelial cells. Circ Res 105, 869–875 (2009).

20. Li, E.C., Heran, B.S. & Wright, J.M. Angiotensin converting enzyme (ACE) inhibitors versus angiotensin receptor blockers for primary hypertension. *Cochrane Database Syst Rev*, CD009096 (2014).

21. Weinbaum, S., Cowin, S.C. & Zeng, Y. A model for the excitation of osteocytes by mechanical loading-induced bone fluid shear stresses. J Biomech 27, 339–360 (1994).

22. Tatsumi, S., et al. Targeted ablation of osteocytes induces osteoporosis with defective mechanotransduction. Cell Metab 5, 464–475 (2007).

23. Miyazaki, T., et al. Mechanical regulation of bone homeostasis through p130Cas- mediated alleviation of NF-κB activity. Sci Adv 5, eaau7802 (2019).

24. Ryu, Y., et al. Mechanical regulation underlies effects of exercise on serotonin-induced signaling in the prefrontal cortex neurons. iScience 23, 100874 (2020).

25. Minami, N., et al. Effects of exercise and β-blocker on blood pressure and baroreflexes in spontaneously hypertensive rats. Am J Hypertens 16, 966–972 (2003).

26. Kim, S.E., et al. Treadmill exercise prevents aging-induced failure of memory through an increase in neurogenesis and suppression of apoptosis in rat hippocampus. Exp Gerontol 45, 357–365 (2010).

27. Bertagnolli, M., et al. Exercise training reduces sympathetic modulation on cardiovascular system and cardiac oxidative stress in spontaneously hypertensive rats. Am J Hypertens 21, 1188–1193 (2008).

28. Chidsey, C.A., Braunwald, E. & Morrow, A.G. Catecholamine excretion and cardiac stores of norepinephrine in congestive heart failure. Am J Med 39, 442–451 (1965).

29. Mullen, R.J., Buck, C.R. & Smith, A.M. NeuN, a neuronal specific nuclear protein in vertebrates. Development 116, 201–211 (1992).

30. Freeman, M.R. Specification and morphogenesis of astrocytes. Science 330, 774–778 (2010).

31. Tamura, K., et al. The physiology and pathophysiology of a novel angiotensin receptor- binding protein ATRAP/Agtrap. Curr Pharm Des 19, 3043–3048 (2013).

32. Huda, F., et al. Distinct transduction profiles in the CNS via three injection routes of AAV9 and the application to generation of a neurodegenerative mouse model. Mol Ther Methods Clin Dev 1, 14032 (2014).

33. Tang, X., et al. “Self-cleaving” 2A peptide from porcine teschovirus-1 mediates cleavage of dual fluorescent proteins in transgenic Eimeria tenella. Vet Res 47, 68 (2016).

34. Saitou, K., et al. Local cyclical compression modulates macrophage function in situ and alleviates immobilization-induced muscle atrophy. Clin Sci (Lond) 132, 2147–2161 (2018).

35. Cserr, H.F. & Patlak, C.S. Secretion and bulk flow of interstitial fluid. in Physiology and Pharmacology of the Blood-Brain Barrier. ed. Bradbury, M.W.B. 245–261. Springer Berlin Heidelberg, Berlin, Heidelberg (1992).

36. Chatterjee, K., Carman-Esparza, C.M. & Munson, J.M. Methods to measure, model and manipulate fluid flow in brain. J Neurosci Methods 333, 108541 (2020).

37. Geer, C.P. & Grossman, S.A. Interstitial fluid flow along white matter tracts: a potentially important mechanism for the dissemination of primary brain tumors. J Neurooncol 32, 193–201 (1997).

38. Kingsmore, K.M., et al. MRI analysis to map interstitial flow in the brain tumor microenvironment. APL Bioeng 2 (2018).

39. Hrabetova, S. & Nicholson, C. Biophysical Properties of Brain Extracellular Space Explored with Ion-Selective Microelectrodes, Integrative Optical Imaging and Related Techniques. in Electrochemical Methods for Neuroscience. eds. Michael, A.C. & Borland, L.M. (2007).

40. Gong, S., et al. A gene expression atlas of the central nervous system based on bacterial artificial chromosomes. Nature 425, 917–925 (2003).

41. Goshima, Y., Ohsako, S. & Yamauchi, T. Overexpression of Ca^2+^/calmodulin-dependent protein kinase II in Neuro2A and NG108-15 neuroblastoma cell lines promotes neurite outgrowth and growth cone motility. J Neurosci 13, 559–567 (1993).

42. Yun, J., et al. Neuronal Per Arnt Sim (PAS) domain protein 4 (NPAS4) regulates neurite outgrowth and phosphorylation of synapsin I. J Biol Chem 288, 2655–2664 (2013).

43. Hamaguchi, R., Takemori, K., Inoue, T., Masuno, K. & Ito, H. Short-term treatment of stroke-prone spontaneously hypertensive rats with an AT1 receptor blocker protects against hypertensive end-organ damage by prolonged inhibition of the renin-angiotensin system. Clin Exp Pharmacol Physiol 35, 1151–1155 (2008).

44. Oddie, C.J., Dilley, R.J. & Bobik, A. Long-term angiotensin II antagonism in spontaneously hypertensive rats: effects on blood pressure and cardiovascular amplifiers. Clin Exp Pharmacol Physiol 19, 392–395 (1992).

45. Rezacova, L., et al. Role of angiotensin II in chronic blood pressure control of heterozygous Ren-2 transgenic rats: Peripheral vasoconstriction versus central sympathoexcitation. Biomed Pharmacother 116, 108996 (2019).

46. Husken, B.C., Hendriks, M.G., Pfaffendorf, M. & Van Zwieten, P.A. Effects of aging and hypertension on the reactivity of isolated conduit and resistance vessels. Microvasc Res 48, 303–315 (1994).

47. Sepehrdad, R., et al. Amiloride reduces stroke and renalinjury in stroke-prone hypertensive rats. Am J Hypertens 16, 312–318 (2003).

48. Pirici, D., et al. Common impact of chronic kidney disease and brain microhemorrhages on cerebral Aβ pathology in SHRSP. Brain Pathol 27, 169–180 (2017).

49. Smeda, J.S. Renal function in stroke-prone rats fed a high-K+ diet. Can J Physiol Pharmacol 75, 796–806 (1997).

50. Fujiyabu, T., Toni, F., Li, X., Chung, U.I. & Sakai, T. Three cooperative diffusion coefficients describing dynamics of polymer gels. Chem Commun (Camb) 54, 6784–6787 (2018).

51. Min, S., et al. Arterial baroreceptors sense blood pressure through decorated aortic claws. Cell Rep 29, 2192–2201 e2193 (2019).

52. Furlan, R., et al. Oscillatory patterns in sympathetic neural discharge and cardiovascular variables during orthostatic stimulus. Circulation 101, 886–892 (2000).

53. Milutinovic, S., Murphy, D. & Japundzic-Zigon, N. Central cholinergic modulation of blood pressure short-term variability. Neuropharmacology 50, 874–883 (2006).

54. Pagani, M., et al. Power spectral analysis of heart rate and arterial pressure variabilities as a marker of sympatho-vagal interaction in man and conscious dog. Circ Res 59, 178–193 (1986).

55. Billman, G.E. The LF/HF ratio does not accurately measure cardiac sympatho-vagal balance. Front Physiol 4, 26 (2013).

56. Reyes del Paso, G.A., Langewitz, W., Mulder, L.J., van Roon, A. & Duschek, S. The utility of low frequency heart rate variability as an index of sympathetic cardiac tone: a review with emphasis on a reanalysis of previous studies. Psychophysiology 50, 477–487 (2013).

57. Joyner, M.J. & Green, D.J. Exercise protects the cardiovascular system: effects beyond traditional risk factors. J Physiol 587, 5551–5558 (2009).

58. Jancovski, N., et al. Angiotensin type 1A receptor expression in C1 neurons of the rostral ventrolateral medulla contributes to the development of angiotensin-dependent hypertension. Exp Physiol 99, 1597–1610 (2014).

59. Guo, F., et al. Astroglia are a possible cellular substrate of angiotensin(1-7) effects in the rostral ventrolateral medulla. Cardiovasc Res 87, 578–584 (2010).

60. Kishi, T., et al. Exercise training causes sympathoinhibition through antioxidant effect in the rostral ventrolateral medulla of hypertensive rats. Clin Exp Hypertens 34, 278–283 (2012).

61. Liddelow, S.A. & Barres, B.A. Reactive Astrocytes: Production, Function, and Therapeutic Potential. Immunity 46, 957–967 (2017).

62. Maclullich, A.M., et al. Higher systolic blood pressure is associated with increased water diffusivity in normal-appearing white matter. Stroke 40, 3869–3871 (2009).

63. Bedussi, B., et al. Enhanced interstitial fluid drainage in the hippocampus of spontaneously hypertensive rats. Sci Rep 7, 744 (2017).

64. Iskratsch, T., Wolfenson, H. & Sheetz, M.P. Appreciating force and shape-the rise of mechanotransduction in cell biology. Nat Rev Mol Cell Biol 15, 825–833 (2014).

65. Hughes, J.W., Watkins, L., Blumenthal, J.A., Kuhn, C. & Sherwood, A. Depression and anxiety symptoms are related to increased 24-hour urinary norepinephrine excretion among healthy middle-aged women. J Psychosom Res 57, 353–358 (2004).

66. Tanaka, H., et al. Swimming training lowers the resting blood pressure in individuals with hypertension. J Hypertens 15, 651–657 (1997).

67. Matsusaki, M., et al. Influence of workload on the antihypertensive effect of exercise. Clin Exp Pharmacol Physiol 19, 471–479 (1992).

68. Li, S.C., et al. Tissue elasticity bridges cancer stem cells to the tumor microenvironment through microRNAs: Implications for a “watch-and-wait” approach to cancer. Curr Stem Cell Res Ther 12, 455–470 (2017).

69. Kishi, T., et al. Increased reactive oxygen species in rostral ventrolateral medulla contribute to neural mechanisms of hypertension in stroke-prone spontaneously hypertensive rats. Circulation 109, 2357–2362 (2004).

70. Luo, J., Borgens, R. & Shi, R. Polyethylene glycol improves function and reduces oxidative stress in synaptosomal preparations following spinal cord injury. J Neurotrauma 21, 994–1007 (2004).

71. Guest, J., Benavides, F., Padgett, K., Mendez, E. & Tovar, D. Technical aspects of spinal cord injections for cell transplantation. Clinical and translational considerations. Brain Res Bull 84, 267–279 (2011).

72. Peterson, S.L. & Armstrong, J.J. Muscarinic receptors mediate carbachol-induced inhibition of maximal electroshock seizures in the nucleus reticularis pontis oralis. Epilepsia 40, 20–25 (1999).

73. Faingold, C.L., et al. GABA in the inferior colliculus plays a critical role in control of audiogenic seizures. Brain Res 640, 40–47 (1994).

74. Shi, C., et al. Transportation in the interstitial space of the brain can be regulated by neuronal excitation. Sci Rep 5, 17673 (2015).

75. Li, G.Q., et al. Role of cannabinoid receptor type 1 in rostral ventrolateral medulla in high-fat diet-induced hypertension in rats. J Hypertens 36, 801–808 (2018).

76. Carlson, D.J., Dieberg, G., Hess, N.C., Millar, P.J. & Smart, N.A. Isometric exercise training for blood pressure management: a systematic review and meta-analysis. Mayo Clin Proc 89, 327–334 (2014).

77. Limpert, E., Stahel, W.A. & Abbt, M. Log-normal distributions across the sciences: Keys and clues: On the charms of statistics, and how mechanical models resembling gambling machines offer a link to a handy way to characterize log-normal distributions, which can provide deeper insight into variability and probability—normal or log-normal: That is the question. BioScience 51, 341–352 (2001).

78. Byun, S., Hecht, V.C. & Manalis, S.R. Characterizing cellular biophysical responses to stress by relating density, deformability, and size. Biophys J 109, 1565–1573 (2015).

79. ISO TS 21362:2018(E). Nanotechnologies − Analysis of nano-objects using asymmetrical-flow and centrifugal field flow fractionation (2018)

80. Carman, P.C. Fluid flow through granular beds. Chemical Engineering Research and Design 75, S32–S48 (1997).

81. Xu, P. & Yu, B. Developing a new form of permeability and Kozeny–Carman constant for homogeneous porous media by means of fractal geometry. Advances in Water Resources 31, 74–81 (2008).

82. Pedersen, J.A., Boschetti, F. & Swartz, M.A. Effects of extracellular fiber architecture on cell membrane shear stress in a 3D fibrous matrix. J Biomech 40, 1484–1492 (2007).

83. Yetkin, F., et al. Cerebrospinal fluid viscosity: a novel diagnostic measure for acute meningitis. South Med J 103, 892–895 (2010).

84. Bloomfield, I.G., Johnston, I.H. & Bilston, L.E. Effects of proteins, blood cells and glucose on the viscosity of cerebrospinal fluid. Pediatr Neurosurg 28, 246–251 (1998).

85. Tarbell, J.M. & Shi, Z.-D. Effect of the glycocalyx layer on transmission of interstitial flow shear stress to embedded cells. Biomech Model Mechanobiol 12, 111–121 (2013).

86. Sakai, K., et al. Overexpression of eNOS in NTS causes hypotension and bradycardia in vivo. Hypertension 36, 1023–1028 (2000).

87. Kishi, T., et al. Overexpression of eNOS in the RVLM causes hypotension and bradycardia via GABA release. Hypertension 38, 896–901 (2001).

88. Paxinos, G. & Watson, C. The rat brain in stereotaxic coordinates, (Academic Press, 1998).

89. Hirooka, Y., Polson, J.W. & Dampney, R.A. Pressor and sympathoexcitatory effects of nitric oxide in the rostral ventrolateral medulla. J Hypertens 14, 1317–1324 (1996).

90. Shinohara, Y., et al. Effects of neutralizing antibody production on AAV-PHP.B-mediated transduction of the mouse central nervous system. Mol Neurobiol 56, 4203–4214 (2019).

91. Shinohara, Y., Ohtani, T., Konno, A. & Hirai, H. Viral vector-based evaluation of regulatory regions in the neuron-specific enolase (NSE) promoter in mouse cerebellum in vivo. Cerebellum 16, 913–922 (2017).

92. Konno, A., et al. Mutant ataxin-3 with an abnormally expanded polyglutamine chain disrupts dendritic development and metabotropic glutamate receptor signaling in mouse cerebellar Purkinje cells. Cerebellum 13, 29–41 (2014).

93. Matsuzaki, Y., et al. Neurotropic Properties of AAV-PHP.B are shared among diverse inbred strains of mice. Mol Ther 27, 700–704 (2019).

94. Sakitani, N., et al. Application of consistent massage-kike perturbations on mouse calves and monitoring the resulting intramuscular pressure changes. J Vis Exp (2019).

95. Diaspro, A., et al. Multi-photon excitation microscopy. Biomed Eng Online 5, 36 (2006).

96. Lei, Y., Han, H., Yuan, F., Javeed, A. & Zhao, Y. The brain interstitial system: Anatomy, modeling, in vivo measurement, and applications. Prog Neurobiol 157, 230–246 (2017).

97. Goerzen, D., et al. An MRI-derived neuroanatomical atlas of the Fischer 344 rat brain. Sci Rep 10, 6952 (2020).

98. Liu, C., De Luca, A., Rosso, A. & Talon, L. Darcy’s law for yield stress fluids. Phys Rev Lett 122, 245502 (2019).

99. Johnson, A.K., Yarin, A.L. & Mashayek, F. Packing density and the Kozeny-Carman equation. Neurosurgery 71, E1064–1065; author reply E1065-1066 (2012).

100. de Vellis, J. & Cole, R. Preparation of mixed glial cultures from postnatal rat brain. Methods Mol Biol 814, 49–59 (2012).

101. Nagao, M., et al. Coordinated control of self-renewal and differentiation of neural stem cells by Myc and the p19ARF-p53 pathway. J Cell Biol 183, 1243–1257 (2008).

102. Sato, T., et al. New simple methods for isolating baroreceptor regions of carotid sinus and aortic depressor nerves in rats. Am J Physiol 276, H326–332 (1999).

103. Umemura, S., et al. The Japanese Society of Hypertension guidelines for the management of hypertension (JSH 2019). Hypertens Res 42, 1235–1481 (2019).

104. Oppenheim, A.V., Schafer, R.W., Buck, J.R., Buck, J.R. & Schafer, R.W. Discrete-time signal processing, (Prentice Hall, Upper Saddle River, N.J, 1999).

105. King, C.E., et al. Postoperative continuous non-invasive cardiac output monitoring on the ward: a feasibility study. J Clin Monit Comput 35, 1349–1356 (2021).

106. Ogata, H., Kayaba, M., Kaneko, M., Ogawa, K. & Kiyono, K. Evaluation of sleep quality in a disaster evacuee environment. Int J Environ Res Public Health 17(2020).

